# Does where you live influence what you are made of? Spatial correlates of chemical traits across commonly occurring boreal plants

**DOI:** 10.1101/2021.01.26.428320

**Authors:** Travis R Heckford, Shawn J. Leroux, Eric Vander Wal, Matteo Rizzuto, Juliana Balluffi-Fry, Isabella C. Richmond, Yolanda F. Wiersma

## Abstract

**Context:** Spatially explicit drivers of foliar chemical traits link plants to ecosystem processes to reveal landscape functionality. Specifically, foliar elemental, stoichiometric, and phytochemical (ESP) compositions represent key indicator traits.

**Objectives:** Here, we investigate the spatial drivers of foliar ESP at the species level and across species at the trait level for five commonly occurring boreal forest understory plants.

**Methods:** On the island of Newfoundland, Canada, we collected foliar material from four chronosequenced forest grids. Using response variables of foliar elemental (C, N, P, percent and quantity), stoichiometric (C:N, C:P, N:P), and phytochemical (terpenoids) composition, we tested multiple competing hypotheses using spatial predictors of land cover (e.g., coniferous, deciduous, mixedwood), productivity (e.g., enhanced vegetation index), biotic (e.g., stand age/height, canopy closure) and abiotic (e.g., elevation, aspect, slope) factors.

**Results:** We found evidence to support spatial relationships of foliar ESP for most species (mean R^2^ = 0.22, max = 0.65). Spatial variation in elemental quantity traits of C, N, P were related to land cover along with biotic and abiotic factors for 2 of 5 focal species. Notably, foliar C, C:P, and sesquiterpene traits between different species were related to abiotic factors. Similarly, foliar terpenoid traits between different species were related to a combination of abiotic and biotic factors (mean R^2^ = 0.26).

**Conclusions:** Spatial-trait relationships mainly occur at the species level, with some commonalities occurring at the trait level. By linking foliar ESP traits to spatial predictors, we can map plant chemical composition patterns that influence landscape-scale ecosystem processes.

## 1. Introduction

Environmental factors are known to influence the foliar traits of plants. For instance, differences in overstory vegetation (i.e., landcover; Hallett & Hornbeck, 1997), productivity (Radwan & Harrington, 2011), community structure (Sedio et al., 2017), and topographic conditions (Müller et al., 2017) may influence foliar chemical, physiological, and morphological traits (Poorter & Bongers, 2006). Chemical traits such as elemental concentration (% and quantity Carbon, Nitrogen, and Phosphorus), stoichiometric ratios (elemental concentrations on a biomass basis, specifically molar C:N, N:P, and C:P ratios), and secondary carbon based compounds (terpenoids, phenols) are often useful indicators of ecosystem processes. These traits can be indicators of processes such as decomposition (Diaz et al., 2004), carbon sequestration/primary production (Harpole et al., 2011; Hessen et al., 2004), evapotranspiration (Liu et al., 2019), and trophic interactions (Bryant et al., 1983; Hunter, 2016). Environmental factors vary across the landscape and thus, species level intraspecific trait variability (ITV) mapped in response to spatial gradients of varying environmental conditions may reveal how underlining ecological processes contribute to spatial patterns that define landscape functionality (Harvey et al., 2017; Schmitz et al., 2018). As well, across species, common environmental factors may drive interspecific trait variability, and if so, this provides room to devise community-level generalities of landscape function (Santiago et al., 2004). Here, we investigate which environmental factors drive the spatial variability of foliar elemental (E), stoichiometric (S) and phytochemical (P) traits (hereafter labelled “ESP traits”) at a species level for common boreal plants, and we compare these factors across the five species to determine if there are shared community-level drivers of traits.

Across the landscape, differing environmental conditions influence plant trade-offs of resource acquisition and use, and as such the ITV of foliar traits (Lavorel et al., 2011). For instance, plants growing under different overstory vegetation (e.g., deciduous, coniferous, and mixedwood land cover types), which experience varied light conditions via canopy vertical and horizontal composition, may redistribute foliar N and P resources to optimize growth while stabilizing for competitive interactions (Hassell et al., 1994). As well, nutrient recycling pathways may vary by landcover types via litter inputs and canopy temperature/precipitation controls (Barron-Gafford et al., 2012; Philben et al., 2016) which can influence soil productivity (Krishna & Mohan, 2017) N and P resource availability (Gartner & Cardon, 2004; Knops et al., 2002), and plant N and P use efficiencies (Ashton et al., 2010). Moreover, topographic gradients of elevation, aspect, and slope further define temperature, precipitation, and solar insolation inputs (Macek et al., 2019) and as such can influence resource allocation (Müller et al., 2017). Indeed, many factors likely influence resource trade-offs by plants and their foliar ESP traits, with the range of ITV constrained by a species resource strategy (Grime & Pierce, 2012). Spatial gradients of environmental conditions create a landscape of resource trade-offs where the ITV of foliar ESP traits provides mapped heterogeneity of inferred ecosystem processes.

Identifying the spatial covariates of traits linked to ecosystem processes is an important topic in landscape ecology (Pickett & Cadenasso, 1995; Turner, 1989). For instance, the distribution and movement of energy and matter is a central focus for understanding landscape functionality via pattern and process relationships (Lavorel et al., 2011; Shen et al., 2011; Monica Goigel Turner, 2005). Foliar ESP traits provide a direct link to thermodynamics and entropy processes at landscape extents (Elser & Hamilton, 2007; Vranken et al., 2015). For example, foliar N and P concentration and N:P ratios have been linked to primary productivity (Elser et al., 2010), while stoichiometric traits have been associated with nutrient limitation and community structure processes (Harpole et al., 2011; Urbina et al., 2017). Phytochemical defense traits have been linked to trophic interactions, spatial flows of energy and matter, and nutrient recycling processes (Hunter, 2016). At the landscape level, spatial covariates of land cover, productivity, forest structure, and topography are known drivers of foliar ESP trait variability. However, different covariates likely influence different foliar traits between species. For example, balsam fir (*Abies balsamea*) and red spruce (*Picea rubens*) foliar N and P follow elevational gradients (Richardson, 2004), while Scots pine (*pinus sylvestris*) shifts foliar stoichiometric content in response to soil nutrients (i.e., site level productivity; He et al., 2019), and eucalyptus (*Eucalyptus urophylla*) foliar P decreases with stand age (Fan et al., 2015). Thus, a species level approach to identifying spatial covariates of foliar ESP traits will allow us to obtain refined estimates that are comparable across species and traits to derive potential generalities.

Here, we use spatially explicit covariates to investigate correlates of foliar ESP traits for five commonly occurring juvenile boreal forest species. Our spatial predictors of land cover (i.e., coniferous, deciduous, mixedwood), productivity (i.e., enhanced vegetation index), biotic factors (i.e., structural conditions of stand age, height, and canopy closure), and abiotic factors (i.e., elevation, aspect, and slope), represent known and/or suggested drivers of foliar ESP traits (see Table 1). Our aim is to investigate the spatial relationships influencing foliar ESP traits by interrogating covariate selection for generalities at the trait and species level. Our integrative approach investigates multiple components of foliar elemental and nutritional traits and their spatial drivers. This allows us to link spatial patterns to ecosystem processes that contribute to landscape function.

**Table 1.**
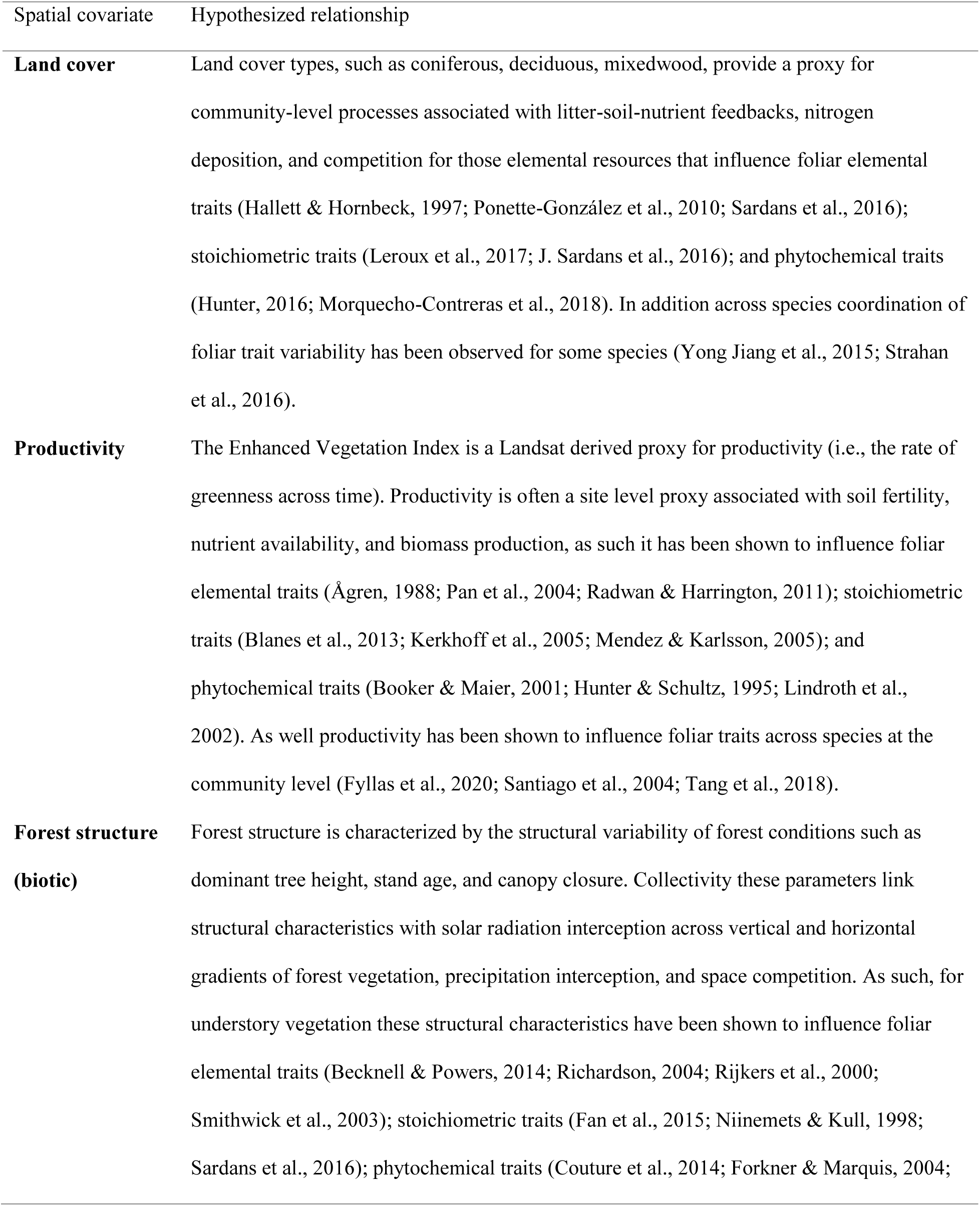
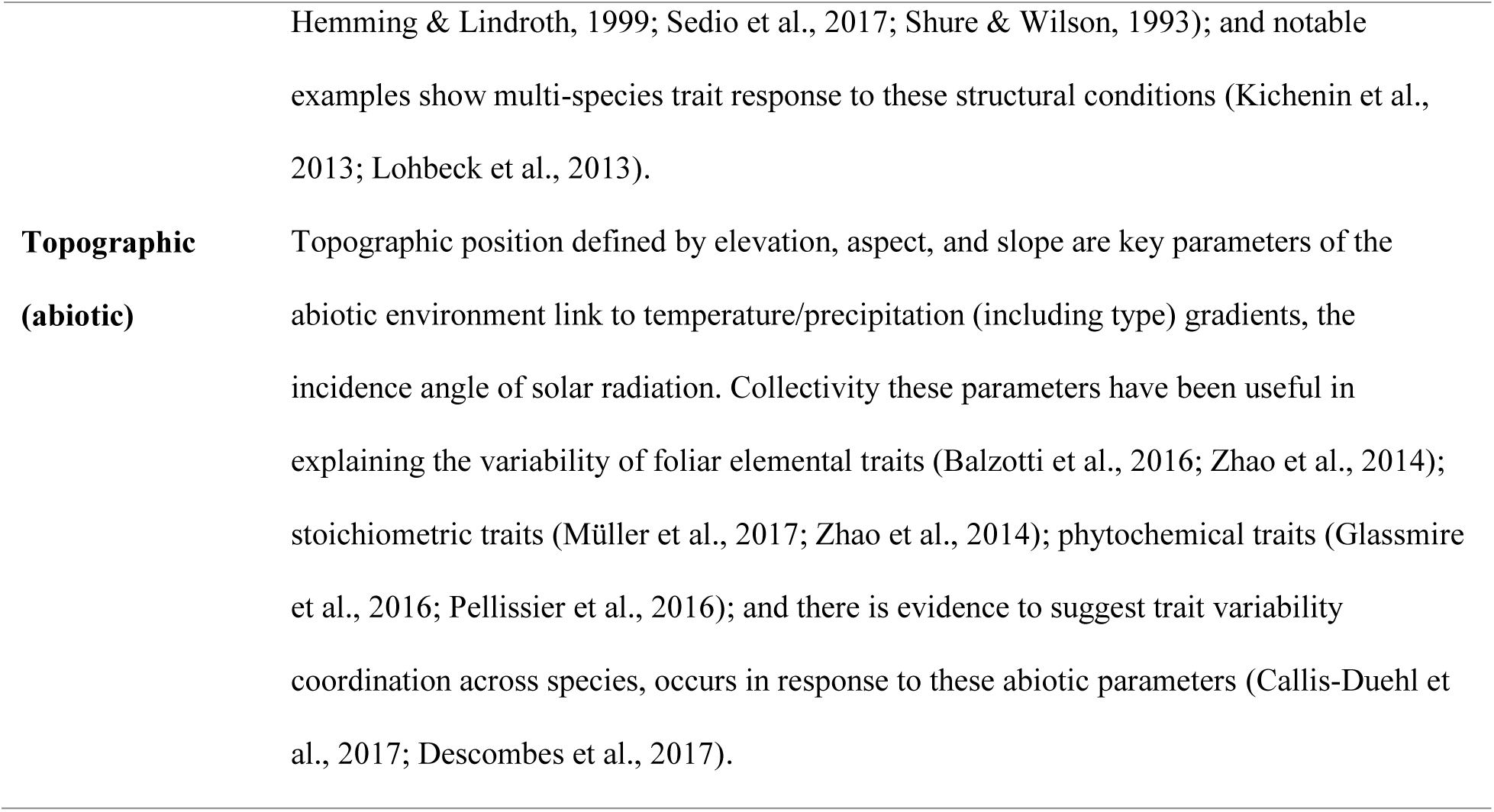
Hypotheses for land cover, productivity, biotic (forest structure: age, height, canopy closure) and abiotic (elevation, aspect, slope) spatial covariates relationship to the variability of foliar elemental, stoichiometric, and phytochemical traits. For each spatial covariate we provide references to foliar ESP traits and to community level coordination of trait variability. Our approach does not consider a community weighted assessment of foliar ESP traits across species, instead we compare spatial covariates at the trait and species level to investigate potential commonalities.

| Spatial covariate | Hypothesized relationship |
| --- | --- |
| **Land cover** | Land cover types, such as coniferous, deciduous, mixedwood, provide a proxy for community-level processes associated with litter-soil-nutrient feedbacks, nitrogen deposition, and competition for those elemental resources that influence foliar elemental traits (Hallett & Hornbeck, 1997; Ponette-González et al., 2010; Sardans et al., 2016); stoichiometric traits (Leroux et al., 2017; J. Sardans et al., 2016); and phytochemical traits (Hunter, 2016; Morquecho-Contreras et al., 2018). In addition across species coordination of foliar trait variability has been observed for some species (Yong Jiang et al., 2015; Strahan et al., 2016). |
| **Productivity** | The Enhanced Vegetation Index is a Landsat derived proxy for productivity (i.e., the rate of greenness across time). Productivity is often a site level proxy associated with soil fertility, nutrient availability, and biomass production, as such it has been shown to influence foliar elemental traits (Ågren, 1988; Pan et al., 2004; Radwan & Harrington, 2011); stoichiometric traits (Blanes et al., 2013; Kerkhoff et al., 2005; Mendez & Karlsson, 2005); and phytochemical traits (Booker & Maier, 2001; Hunter & Schultz, 1995; Lindroth et al., 2002). As well productivity has been shown to influence foliar traits across species at the community level (Fyllas et al., 2020; Santiago et al., 2004; Tang et al., 2018). |
| **Forest structure (biotic)** | Forest structure is characterized by the structural variability of forest conditions such as dominant tree height, stand age, and canopy closure. Collectively these parameters link structural characteristics with solar radiation interception across vertical and horizontal gradients of forest vegetation, precipitation interception, and space competition. As such, for understory vegetation these structural characteristics have been shown to influence foliar elemental traits (Becknell & Powers, 2014; Richardson, 2004; Rijkers et al., 2000; Smithwick et al., 2003); stoichiometric traits (Fan et al., 2015; Niinemets & Kull, 1998; Sardans et al., 2016); phytochemical traits (Couture et al., 2014; Forkner & Marquis, 2004; |
|  | Hemming & Lindroth, 1999; Sedio et al., 2017; Shure & Wilson, 1993); and notable examples show multi-species trait response to these structural conditions (Kichenin et al., 2013; Lohbeck et al., 2013). |
| **Topographic**<br>**(abiotic)** | Topographic position defined by elevation, aspect, and slope are key parameters of the abiotic environment link to temperature/precipitation (including type) gradients, the incidence angle of solar radiation. Collectivity these parameters have been useful in explaining the variability of foliar elemental traits (Balzotti et al., 2016; Zhao et al., 2014); stoichiometric traits (Müller et al., 2017; Zhao et al., 2014); phytochemical traits (Glassmire et al., 2016; Pellissier et al., 2016); and there is evidence to suggest trait variability coordination across species, occurs in response to these abiotic parameters (Callis-Duehl et al., 2017; Descombes et al., 2017). |

## 2. Methods

### 2.1. Study site and focal species description

Our study area is located on the eastern side of the island and Newfoundland, Canada (Fig. 1a; a detailed description of Fig. 1, is provided in Appendix 1). Here, the bedrock is generally a mixture of crystalline Paleozoic strata with upland dominated by hummocky to ridged sandy morainal depositions (South, 1983). The vegetative cover is dominated primarily by intermediate-aged, closed canopy, forest stands of balsam fir (*Abies balsamea*) and black spruce (*Picea mariana*) on steep, moist, upland areas. Alternatively, disturbed areas are dominated by paper birch (*Betula papyrifera*), trembling aspen (*Populus tremuloides*), and black spruce with drier sites consisting of black spruce and heaths of kalmia (*Kalmia angustifolia*) (South, 1983). On average this region experiences annual temperature of 4.5°C, with a summer and winter mean of 12.5°C and −3.5°C, and mean annual precipitation of 100-300 cm (South, 1983).

**Figure 1.**
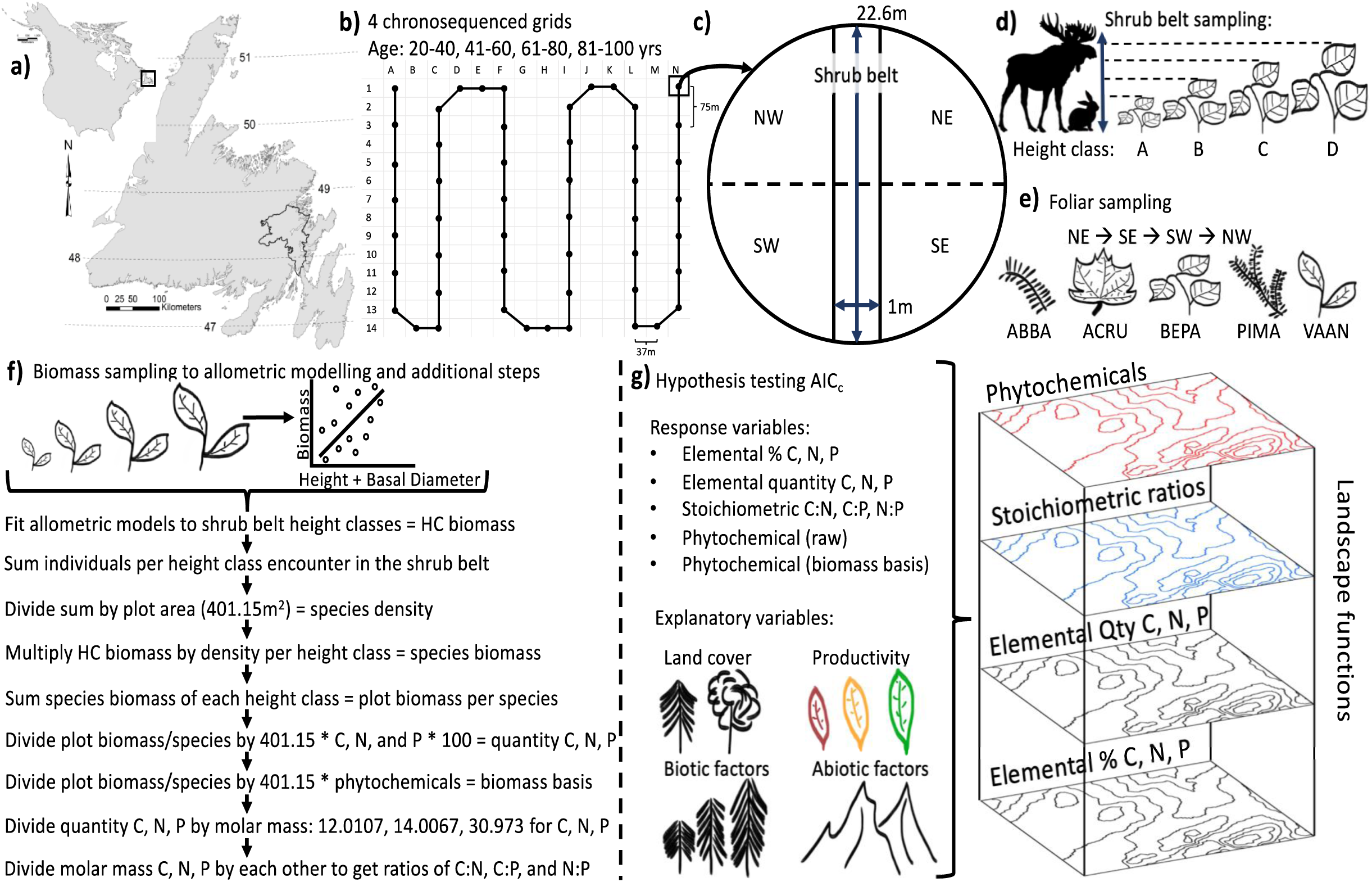
The roadmap of our methods adapted from Leroux et al. 2017. Our study location occurred on the island of Newfoundland, Canada (a) where we set up four chronosequenced meandering transect grids each consisting of 50 sampling locations (b). At each sample location we set up 22.6 m diameter circular plots (c), and along a 22.6 m long, 1 m wide shrub belt (c) we collected density measures of our study species for a max of five per height class: 0-50 cm, 51-100 cm, 101-150 cm, 151-200 cm, coded as A, B, C, and D, respectively (d). We collected foliar samples in each intercardinal corner of the sample plot, starting in the NE corner and moving clockwise until a sufficient and representative sample was acquired (e). Species codes used: balsam fir (ABBA), red maple (ACRU), white birch (BEPA), black spruce (PIMA), and lowbush blueberry (VAAN) (e). We collected biomass samples (i.e., all new growth foliar material) on the periphery of the grids from approximately fifty individuals distributed across height classes (f). Allometric models were fit using biomass as a function of height and basal diameter, from which we parameterized shrub belt correlates to acquire plot level biomass estimates. We used these estimates to determine foliar elemental quantity, stoichiometric ratios, and phytochemical (biomass) traits relativized to biomass density at the plot level. We fit 16 models, including a null model, for response variables of foliar elemental (percent and quantity), stoichiometric, and phytochemical traits using spatially explicit covariates of land cover, productivity, biotic (stand age, height, canopy closure) and abiotic (elevation, aspect, and slope) factors (g). Using top model coefficient estimates and or average coefficients for competing top models, we constructed spatial surfaces of foliar ESP trait surfaces that link physiological properties to ecosystem processes at the landscape extent.

Our understory focal species consist of two coniferous species: balsam fir (*Abies balsamea*), black spruce (*Picea mariana*), two deciduous species: red maple (*Acer rubrum*), white birch (*Betula papyrifera*), and one herbaceaous plant: lowbush blueberry (*Vaccinium angustifolia*). Our focal species commonly occur across the study region and are largely co-distributed geographically across North America. Moreover, our focal species represent common forage for the dominant herbivores within the boreal system: moose (*Alces alces*) and snowshoe hare (*Lepus americanus*). As such, their foliar traits provide us with a useful measure of resource distribution by which we can infer spatial patterns of herbivory (Balluffi-Fry et al., 2020; Rizzuto et al., 2019).

For each of our study species we assessed foliar traits of elemental concentration (i.e., percent and quantity C, N, and P) and stoichiometric ratio (i.e., C:N, C:P, and N:P). For our coniferous species, balsam fir and black spruce that have constituent phytochemical defence strategies, we assessed foliar phytochemical traits of terpene, monoterpene, monoterpenic alcohol, monoterpenic ester, sesquiterpene, and phytochemical diversity.

### 2.2. Data collection

The following sections describe how we collected shrub belt, foliar material, and biomass data.

#### 2.2.1. Sampling design

In black spruce leading stands, which is the predominant forest type for this region (South, 1983), we set up four chronosequenced meandering transect grids (25 ha), differing in age by 20 year intervals (Fig. 1b; centroid locations for each grid: Bloomfield 48.34°N, −53.98°W; Unicorn 48.63°N, −54.01°W; Terra Nova North 48.62°N, −53.97°W; Dunphy’s Pond 48.49°N, −54.05°W). Although heterogeneity in forest structure does exist across our grids, including differences in tree age, height and canopy density, our sampling locations were designed to capture a representative snapshot of forest structure in this region (see Appendix 2 for a comparison of forest structure sampled versus available). These grids were originally designed for snowshoe hare live trapping, to investigate animal spatial ecology related to spatially variable foliar ESP resources. Each grid is comprised of 50 sampling locations (Fig. 1b).

#### 2.2.2. Shrub belt

At each sample location, we set up a 22.6 m diameter circular plot (Fig. 1c). Within each plot, we collected density estimates for each of our study species along a 22.6 m long and 1 m wide shrub belt transect (Fig. 1c). Moving in a north to south direction, along the belt, for each of our study species encountered, we measured height and basal diameter, and the distance at which it was encountered, for a maximum of five individuals per height class: 0-50 cm, 51-100 cm, 101-150 cm, and 151-200 cm, denoted as A, B, C, and D respectively (Fig. 1d). We restricted our sampling to species within 0-2 m heights as these individuals represent the available forage for common boreal herbivores such as moose and snowshoe hare.

#### 2.2.3. Foliar material

Within our circular plots, we collected representative foliar material from each intercardinal corner. Starting in the NE corner, we clipped foliar material (i.e., terminal and lateral leaves) and then moved to the next corner and clipped a similar amount of foliar material, we continued this process, moving clockwise between the plot corners, until we acquired an approximately 20 g foliar sample. We also measured height and basal diameter (used for augmenting shrub belt data described below) of each individual sampled. Samples of balsam fir (n = 95), black spruce (n = 157), red maple (n = 91), white birch (n = 71), and lowbush blueberry (n = 160) were frozen at −20C until they were sent for foliar elemental analysis at the Agriculture Food Lab (AFL) at the University of Guelph Ontario, Canada. Foliar C and N was determined using an Elementar Vario Macro Cube. Foliar P was determined using a microwave acid digestion CEM MARSxpress microwave system and brought to volume using Nanopure water. The clear extract supernatant was further diluted by 10 to accurately fall within calibration range and reduce high level analyte concentration entering the inductively coupled plasma mass spectrometry (ICP-MS) detector (Poitevin, 2016). Foliar phytochemical analysis for balsam fir (n = 104) and black spruce (n = 163) was performed at the Laboratorie PhytoChemia Inc. in Quebec, Canada, foliar terpenoid composition was determined using a gas chromatography solvent extraction with an internal standard and a correction factor (Cachet et al., 2016).

Elemental/stoichiometric and phytochemical samples differ due to the amount of foliar material needed for each analysis. Less foliar material is needed to perform the phytochemical analysis; thus, we were able to have more samples processed. See Appendix 3 for a complete list of individual terpenoid compounds found in our balsam fir and black spruce foliar samples.

#### 2.2.4. Biomass

To determine the foliar biomass of new growth material for our focal species we collected all of the new growth foliar material from approximately 50 individuals. We collected these individuals along the periphery of our study grids, in randomly selected locations to avoid destructive sampling of foliage in our long-term monitoring grids. We sampled individuals evenly across height classes to obtain a representative sample. In addition, we measured the height and basal diameter for each individual sampled (Fig. 1d). Biomass samples were dried at 50^°^C for 2-3 days. We used the resulting dry weights to perform allometric modelling (described below).

### 2.3. Constructing foliar ESP response variables

Following Leroux et al. (2017), we used three pieces of information to construct foliar ESP distribution models; shrub belt data to determine plot level species density, foliar material to extract elemental percentages (i.e., % C, N, and P) and phytochemical composition (raw basis mg/g), and biomass data to fit allometric models. We fit allometric models using biomass as a function of height and basal diameter for each of our study species (goodness of fit adjusted R^2^ for balsam fir (0.82), black spruce (0.80), red maple (0.83), white birch (0.79), and lowbush blueberry (0.47); see Appendix 4). The estimates of allometric correlates allow us to parameterize shrub belt density data and predict plot level biomass estimates based on density of species in their respective height classes (Fig. 1d,f). We then summed height class biomass estimates at the plot level. In the few instances where we did not encounter a species on the shrub belt but had collected foliar material within that plot, we augmented shrub belt data by adding the total number of individuals sampled for foliar material as ceiling estimate of abundance for a given height class in that plot (see Appendix 5 for details). To acquire foliar elemental quantity traits, we divided plot level biomass by the plot area (401.15 m^2^) multiplied by foliar elemental percentages. To acquire foliar stoichiometric traits, we divided foliar elemental quantity traits of C, N and P by their respective molar masses and divided the resulting values together to get ratios of C:N, C:P, and N:P (Fig. 1f). Similarly, to acquire phytochemical traits, we divided plot level biomass by the plot area (401.15 m^2^) multiplied by our phytochemical raw measures.

### 2.4. Statistical analyses

Data processing and statistical analyses were done using R and Esri software (*Esri*, 2020; R Core Team, 2020). Based on *a priori* reasons we used spatially explicit covariates of land cover, productivity, biotic and abiotic factors, at a resolution of 30 m, to predict ESP trait distribution across the study area (see Table 1 for hypothesis rationale). We investigated the relationship between all possible combinations of the four a priori covariate including a null model (n = 16 total models per response variable, Table 2 for complete model list). In addition, we confirmed the absence of collinearity among our spatial covariates. Our land cover covariate was derived from the Commission for Environmental Cooperation (*Land Cover Map of North American at 30 m Resolution*, 2017) and consists of three categorical conditions coniferous, deciduous and mixed wood. We used the Enhanced Vegetation Index (EVI, 30 m resolution) as a proxy of productivity, which does not saturate as easily as the Normalized Difference Vegetation Index under wet boreal forest conditions (Vermote et al., 2016). Using Forest Resource Inventory (FRI, originally digitized at a 1:12,500 scale and rasterized to a 30 m resolution) spatial datasets provided by the Provincial Government of Newfoundland and the Federal Government of Canada we derived three biotic covariates of stand height, age, and canopy closure, each having four factor levels. Our abiotic factors were derived from a Canadian Digital Elevation Model (*Canadian Digital Elevation Model: Product Specifications-Edition 1.1.*, 2016, originally a 20 m resolution rasterized to a 30 m resolution) and includes covariates of elevation, aspect, and slope (see Appendices 6 and 7 for spatial covariate description and processing).

**Table 2.** List of models used to assess spatial covariates of foliar trait distribution. Land cover and productivity are derived from Landsat 8 scenes. The land cover dataset was acquired from the Commission for Environmental Cooperation and provides general classification of habitat types, i.e., coniferous, deciduous, mixedwood forests, as well as others. Our proxy for productivity was acquired from Landsat 8 as the Enhanced Vegetation Index spectral product. Our biotic factors include the grouped covariates of forest age, height, and canopy density. These variables were derived from Forest Resource Inventory datasets supplied by the Provincial Government of Newfoundland and Labrador and from the Federal Government of Canada’s Park agency. These variables are grouped as their designation of these three measures are contained within a single polygon and represents associated conditions. Similarly, our abiotic factors include the grouped covariates of elevation, aspect, and slope derived from a single source, a Digital Canadian Elevation Model.

| **Model Number** | **Explanatory Variables** |
| --- | --- |
| **1** | ~ Land Cover + Productivity + Biotic + Abiotic |
| **2** | ~ Productivity + Biotic + Abiotic |
| **3** | ~ Land Cover + Biotic + Abiotic |
| **4** | ~ Land Cover + Productivity + Abiotic |
| **5** | ~ Land Cover + Productivity + Biotic |
| **6** | ~ Land Cover + Biotic |
| **7** | ~ Productivity + Biotic |
| **8** | ~ Land Cover + Abiotic |
| **9** | ~ Productivity + Abiotic |
| **10** | ~ Biotic + Abiotic |
| **11** | ~ Land Cover + Productivity |
| **12** | ~ Land Cover |
| **13** | ~ Productivity |
| **14** | ~ Biotic |
| **15** | ~ Abiotic |
| **16** | ~ Null |

We fit General Linear Models (GLM) with the response variables of foliar percent elemental traits (C, N, P as a %), quantity elemental traits (C, N, P as g/m^2^), stoichiometric traits (molar ratios C:N, C:P, and N:P), phytochemical traits for our coniferous species which includes terpene, monoterpene, monoterpenic alcohol, monoterpenic ester, sesquiterpene, and phytochemical diversity on a raw (mg/g) and biomass basis (mg/m^2^). Our terpene variable is the sum of all phytochemical compounds at the plot level. Phytochemical diversity is calculated using a Shannon Diversity Index for all compounds identified per species (i.e., using our balsam fir phytochemical matrix, sites x by individual phytochemical compounds, we calculated alpha diversity; this was performed again for black spruce). We ranked models based on Akaike Information Criterion corrected for small sample size (AIC_c_) and only considered models within < 2 ΔAIC_c_ and those above the null model as having evidence to support a spatial relationship. In addition, we removed models with uninformative variables (Leroux, 2019). If more than one model was within a < 2 ΔAIC_c_ we averaged model coefficients and extracted full coefficient estimates for use in the construction of distribution models (Burnham & Anderson, 2002).

## 3. Results

We begin each section below by reporting patterns and pseudo R^2^ assessments of top ranked models (ΔAIC_c_ < 2, excluding the null model) across all five species and sub-components of foliar traits: elemental (%C, %N, %P, and quantity C, N, and P), stoichiometric (C:N, C:P, N:P ratios), and phytochemical (terpene, monoterpene, monoterpenic alcohol, monoterpenic ester, sesquiterpene, and diversity). In addition, for each section, we report patterns of top ranked models at the species level. Additional supporting results are reported in the appendix, including an AIC_c_ table (Appendix 8), table of coefficient slopes and significance (Appendix 9), distribution plots of pseudo R^2^ for traits (Appendix 10), a comparison of observed versus spatially predicted values (Appendix 11), and model coefficient estimate tables for top ranked models of traits %C, %N, and %P (Appendices 12-14), quantity C, N, and P (Appendix 15), stoichiometric ratios of C:N, C:P and N:P (Appendices 16-18) and phytochemical groups (terpene and monoterpene (Appendix 19), monoterpenic alcohol and ester (Appendix 20), and sesquiterpenes and phytochemical diversity (Appendix 21)). We include the predictive distribution maps of only a subset of the models (Fig. 5); as there were 41 combinations of species-ESP trait models.

### 3.1. Foliar percent elemental traits

Across all species for foliar percent elemental traits (Fig. 2a), eleven models supported the data (R^2^ min = 0.046, max = 0.646, mean = 0.286). At the trait level (Fig. 2a), four models explain foliar percent carbon data (R^2^ min = 0.092, max = 0.646, mean = 0.372), five models explain foliar percent nitrogen data (R^2^ min = 0.071, max = 0.360, mean = 0.233) and six models explain foliar percent phosphorus data (R^2^ min = 0.046, max = 0.472, mean = 0.242).

**Figure 2.**
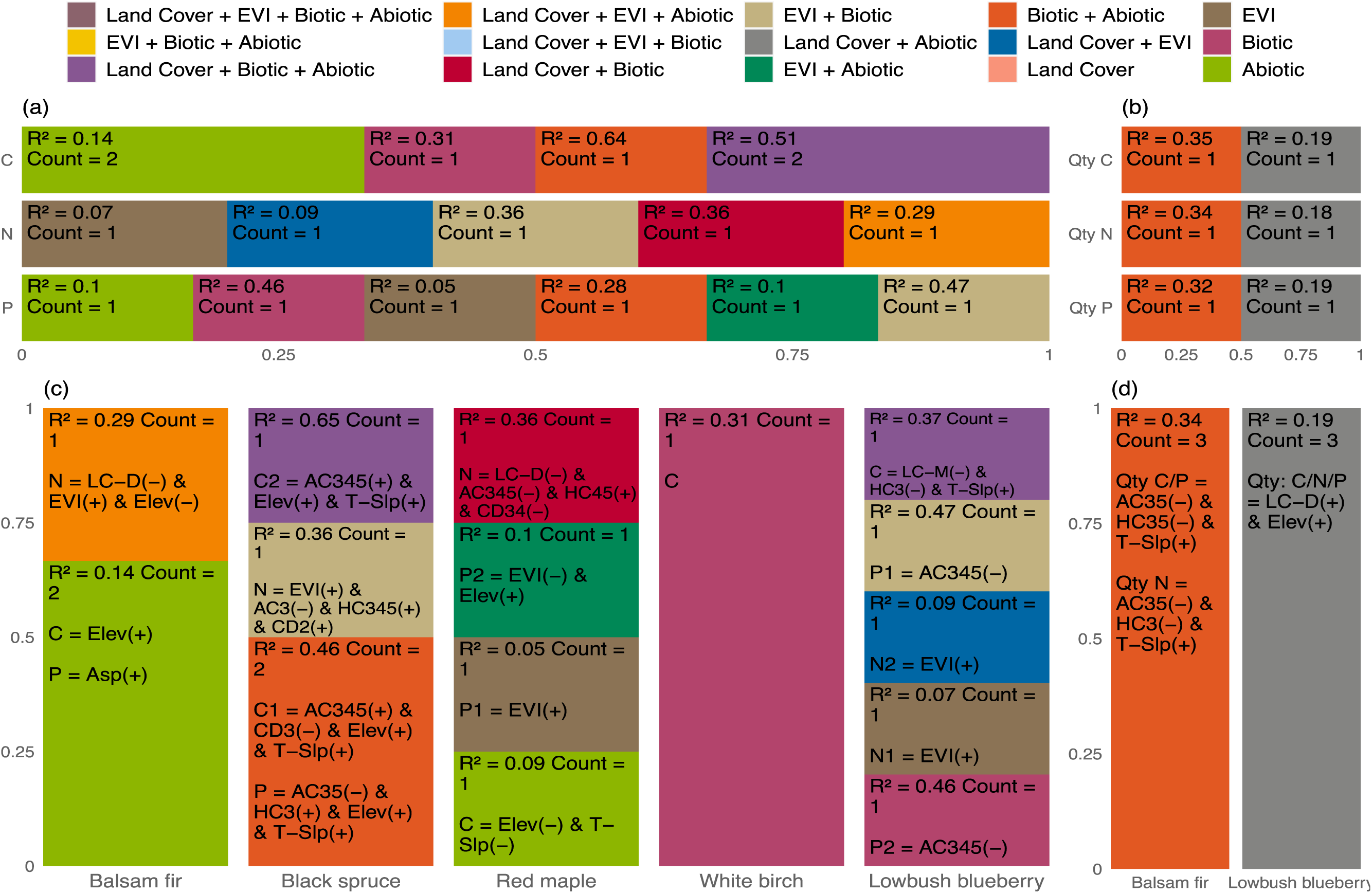
Top ranked model results (i.e., models ΔAIC_c_ < 2) at the trait level (a, b) and species level (c, d) for foliar percent elemental (a, c) and foliar quantity elemental (b, d) traits. Results are organized to show patterns of evidence to support spatial relationships between response and explanatory variables. Superimposed descriptive text on each portion of the stacked bar graphs includes the averaged pseudo R^2^ values for top models if the count > 1, if count is = 1 then only the R^2^ for that model is present. In addition, at the species level (c, d) for our response variables (i.e., C, N, and P) superimposed text indicates significant coefficients and their sign (+/-) for our explanatory variables of land cover, EVI, biotic, and abiotic. Coded values for explanatory variables represent their comprised variables and factor levels. For land cover, LC-C, LC-D, and LC-M indicate coniferous, deciduous, and mixed, respectively. EVI represents the Enhanced Vegetation Index. For biotic variables, AC indicates age class with 3, 4, 5 representing factor levels of 41-60, 61-80, and 81-100 years, respectively. HC indicates height class with 3, 4, 5 representing factor levels of 6.6-9.5, 9.6-12.5, 12.6-15.5 metres, respectively. CD indicates canopy density with 2, 3, 4 representing factor levels of 51-75, 26-50, 10-25 percent closed. For abiotic variables, Elev, Asp, T-Slp represent elevation, aspect, and slope, respectively. If a response variable is supported by more than one top model, a sequential numbering is used to indicate the rank of that model added as a suffix to the response variable text (i.e., C2 indicates the second top ranked model in support of foliar percent carbon). The asterisk symbol (*) is used to indicate that the null model was within ΔAIC_c_< 2. See Appendix 9 for a coefficient signs (+/-) and Appendix 12-15 for coefficient estimates, standard deviations, and confidence intervals.

Although top ranked models vary across species (Fig. 2c), we found trait spatial relationships for all species except white birch foliar percent N and P. Notably, there are different patterns of top ranked models between species and coefficient relationships. For balsam fir, our abiotic model explained foliar percent C and P while N is explained by the combination model of land cover, EVI, and abiotic. For black spruce, our biotic and abiotic model explained foliar percent C and P although our land cover, biotic, and abiotic model is within ΔAIC_c_ < 2 for foliar percent C (model averaged trait distribution map is shown in Fig. 5b). In addition, we found evidence for EVI and biotic model to explain black spruce foliar N. For red maple, foliar percent C is explained by our abiotic model, foliar percent N by our land cover and biotic model, and foliar percent P by two competing top models (1) EVI, and (2) EVI and abiotic. Only white birch foliar percent C is explained by our biotic model. For lowbush blueberry, foliar percent C is explained by our land cover, biotic, and abiotic model. In contrast foliar percent N is explained by two competing top models of (1) EVI, and (2) land cover and EVI, and foliar percent P by is explained by two competing top models of (1) EVI and biotic, and (2) biotic.

### 3.2. Foliar quantity elemental traits

Across all species (Fig. 2b) for foliar elemental quantity traits, two out of the fifteen potential models explain foliar elemental quantity traits (across all traits R^2^ min = 0.183, max = 0.350, mean = 0.263) of C (R^2^ min = 0.193, max = 0.350, mean = 0.271), N (R^2^ min = 0.183, max = 0.345, mean = 0.264), and P (R^2^ min = 0.188, max = 0.321, mean = 0.254). This is, however, only for balsam fir and lowbush blueberry (Fig. 2d). At the species level, balsam fir foliar quantity C, N, and P is explained by our biotic and abiotic model. For lowbush blueberry, foliar quantity C, N, and P is explained by our land cover and abiotic model.

### 3.3. Foliar stoichiometric traits

Across all species (Fig. 3a) twelve of the potential fifteen models explain foliar stoichiometric traits (across all traits R^2^ min = 0.070, max = 0.427, mean = 0.262). At the trait level (Fig. 3a), foliar C:N is explained by five top ranked models (R^2^ min = 0.089, max = 0.385, mean = 0.253). Foliar C:P is explained by four top ranked models (R^2^ min = 0.070, max = 0.336, mean = 0.234). Foliar N:P is explained by six top ranked models (R^2^ min = 0.076, max = 0.427, mean = 0.284).

**Figure 3.**
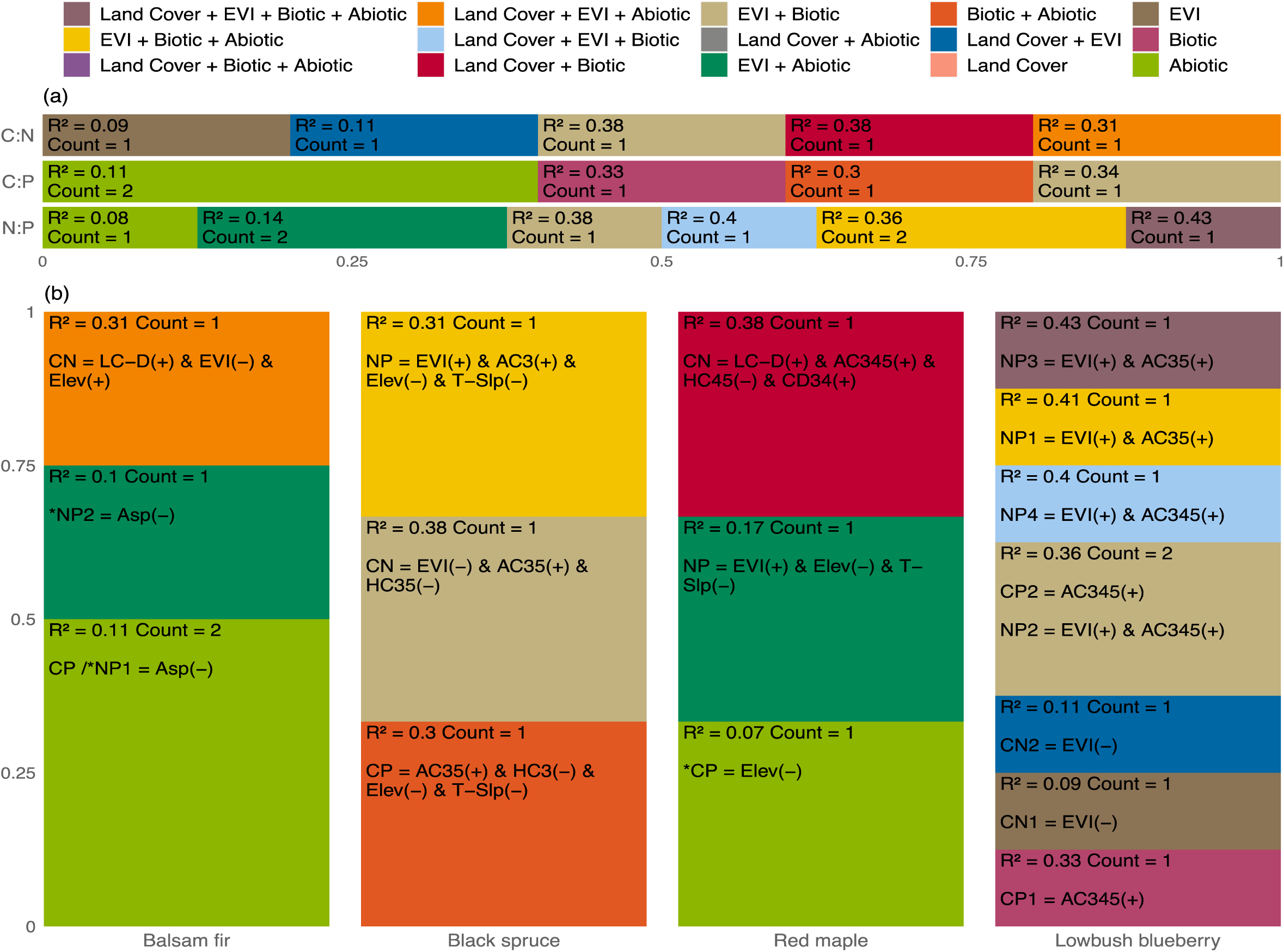
Top ranked model results (i.e., models ΔAIC_c_< 2) at the trait level (a) and species level (b) for foliar stoichiometric traits (i.e., CN, CP, NP). All specifications as in Figure 2. See Appendix 9 for a coefficient signs (+/-) and Appendix 16-18 for coefficient estimates, standard deviations, and confidence intervals.

Again, model specificity is variable at the species level (Fig. 3b), although some geographic commonalities exist in terms of top model covariates and coefficient relationships. For balsam fir, foliar C:N is explained by our land cover, EVI, and abiotic combination model, foliar C:P by our abiotic model, and foliar N:P by two top models (1) abiotic, and (2) EVI and abiotic combination although the null model here was within ΔAIC_c_ < 2. For black spruce, foliar C:N is explained by our EVI and biotic model (model averaged predictive model is shown in Fig. 5b), foliar C:P by our biotic and abiotic model, and foliar N:P by our EVI, biotic and abiotic model. For red maple, foliar C:N is explained by our land cover and biotic model, while our abiotic model explains foliar C:P, however the null model here was within ΔAIC_c_ < 2. In addition, red maple foliar N:P is explained by our land cover and biotic model. For lowbush blueberry, foliar C:N is explained by our EVI model, foliar C:P by competing models of (1) biotic, and (2) EVI and biotic, and foliar N:P by four competing top models of (1) EVI, biotic and abiotic, (2) EVI and biotic, (3) land cover, EVI, biotic and abiotic, and (4) land cover, EVI and biotic. For white birch, the null model was the top ranked for all foliar stoichiometric traits.

### 3.4. Foliar phytochemical traits

Across all species (Fig. 4a) eight of the potential fifteen models explain foliar phytochemical traits on a raw and biomass basis (across all traits R^2^ min = 0.017, max = 0.272, mean = 0.138). At the trait level, terpene raw is explained by three top ranked models (R^2^ min = 0.047, max = 0.270, mean = 0.191), in comparison terpene on a biomass basis is explained by one top ranked model (R^2^ = 0.270). Monoterpene raw is explained by four top ranked models (R^2^ min = 0.041, max = 0.244, mean = 0.121), in comparison monoterpene on a biomass basis is explained by one top ranked model (R^2^ = 0.272). Monoterpenic alcohol raw is explained by two top ranked model (R^2^ min = 0.046, max = 0.233, mean = 0.139). Monoterpenic ester raw is explained by one top ranked model (R^2^ = 0.265), and monoterpenic ester on a biomass basis is also explained by one top ranked model (R^2^ = 0.265). Sesquiterpene raw is explained by seven top ranked models (R^2^ min = 0.040, max = 0.194, mean = 0.098), while sesquiterpene on a biomass basis is explained by two top ranked models (R^2^ min = 0.023, max = 0.242, mean = 0.132). Phytochemical diversity on a raw basis is supported by four top ranked models (R^2^ min = 0.017, max = 0.122, mean = 0.060).

**Figure 4.**
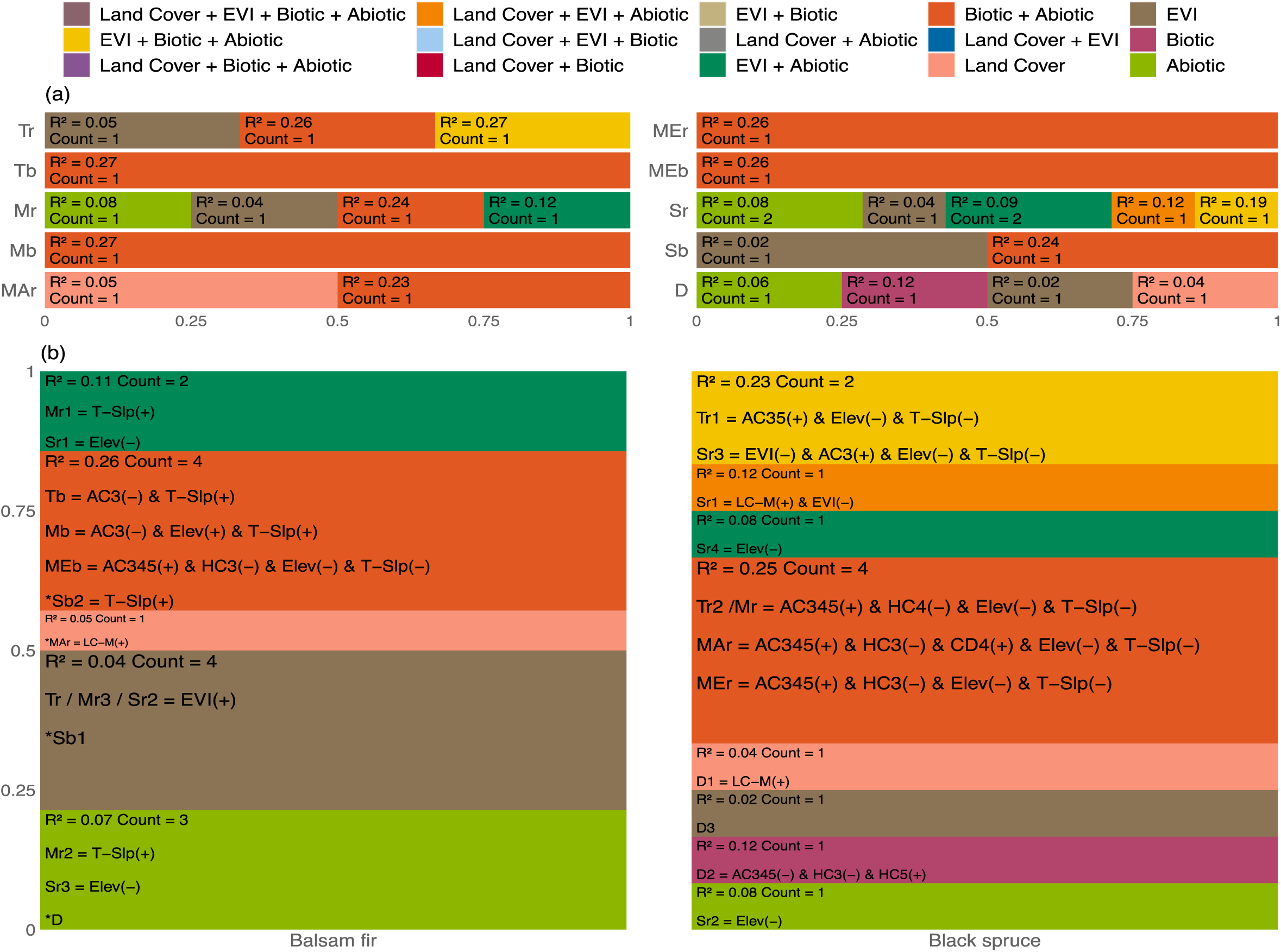
Top ranked model results (i.e., models ΔAIC_c_< 2) at the trait level (a) and species level (b) for foliar phytochemical traits. Coded values are supplied for response variables as with upper case letters representing the trait and lower case letter representing either raw (r) or biomass basis (b). For response variables, T, M, MA, ME, S, and D indicate terpene, monoterpene, monoterpenic alcohol, monoterpenic ester, sesquiterpene, and phytochemical diversity, respectively. All specifications as in Figure 2. See Appendix 9 for a coefficient signs (+/-) and Appendix 19-21 for coefficient estimates, standard deviations, and confidence intervals.

At the species level (Fig. 4b), balsam fir and black spruce share some geographic commonalities in terms of top model covariates and coefficient relationships. For balsam fir, foliar terpene (raw) is explained by our EVI model and terpene on a biomass basis by our biotic and abiotic model. In comparison black spruce foliar terpene (raw) is explained by two competing top models of (1) EVI, biotic and abiotic, and (2) biotic and abiotic (model averaged predictive model shown in Fig. 5d). Three competing top models of (1) EVI and abiotic, (2) abiotic, and (3) EVI explain balsam fir foliar monoterpene (raw), while our biotic and abiotic model explain monoterpene on a biomass basis. In comparison, black spruce foliar monoterpene (raw) is explained by our biotic and abiotic model. Balsam fir foliar monoterpenic alcohol (raw), although the null model is within ΔAIC_c_ < 2, is explained by our land cover model, while black spruce foliar monoterpenic alcohol is explained by our biotic and abiotic combination model. Balsam fir foliar monoterpenic ester on a biomass basis is explained by our biotic and abiotic combination model. While black spruce foliar monoterpenic ester (raw) is explained by our biotic and abiotic combination model. Balsam fir foliar sesquiterpene (raw) is explained by three competing top models of (1) EVI and abiotic, (2) EVI, and (3) abiotic. Balsam fir sesquiterpene on a biomass basis is explained by two competing top models of (1) EVI, and (2) biotic and abiotic, although the null model is within ΔAIC_c_ < 2. In contrast, black spruce foliar sesquiterpene is explained by four competing top models of (1) land cover, EVI and abiotic, (2) abiotic, (3) biotic and abiotic, and (4) EVI and abiotic. Lastly, balsam fir foliar phytochemical diversity is explained by our abiotic model, although the null model is within ΔAIC_c_ < 2, while black spruce foliar phytochemical diversity is explained by three competing top models of (1) land cover, (2) biotic, and (3) EVI.

**Figure 5.**
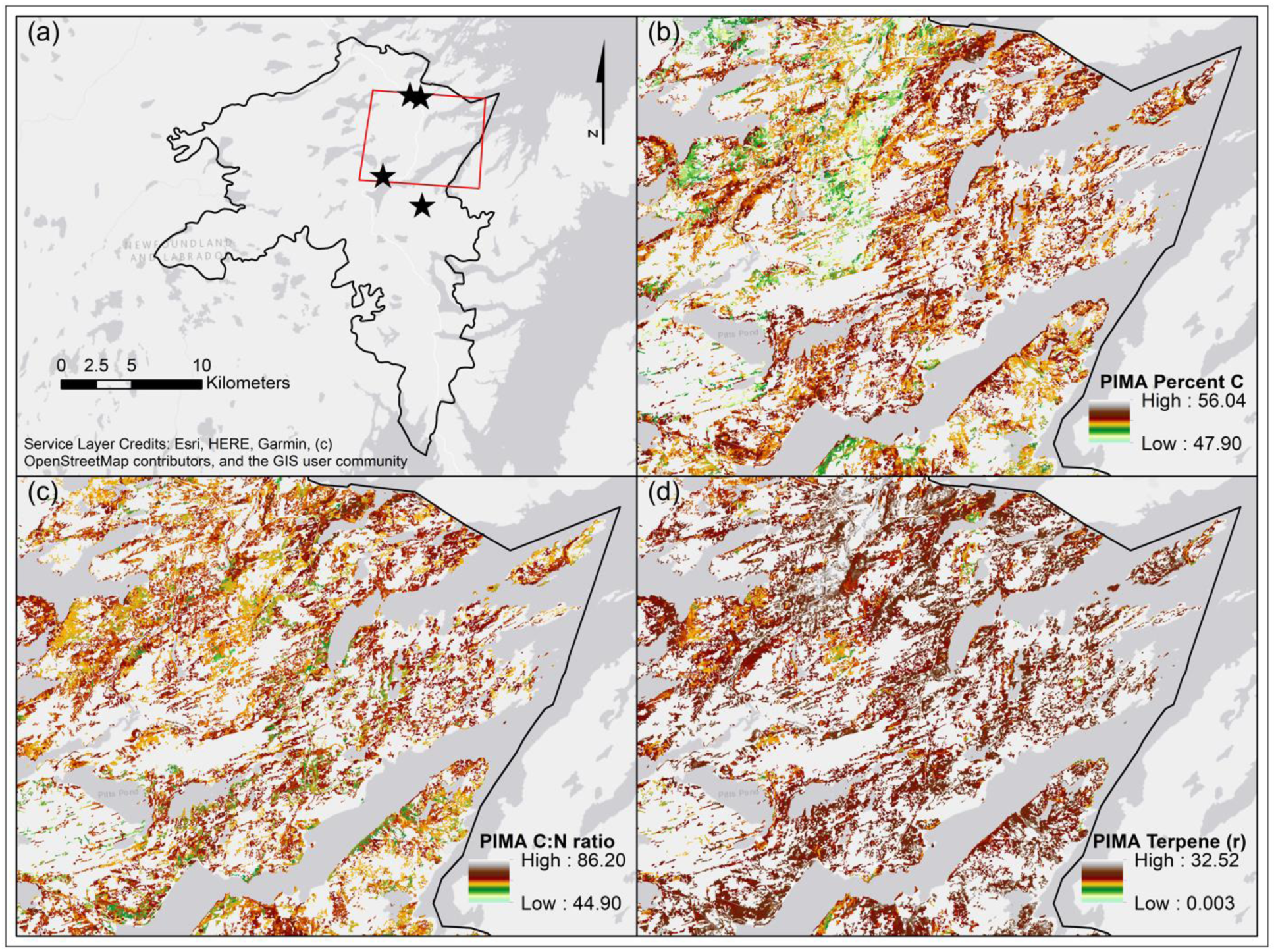
Example of spatially explicit foliar ESP trait distribution models. In (a) we show our spatial area of interest as the black outlined region. Our grid locations are denoted in panel a using the star outline. The red box shown in panel a, is the extent of the subsequent maps provided in this figure, a close up view of spatial foliar ESP patterns for black spruce (PIMA). Foliar percent carbon (b) ranges from 47.9 to 56.04 and is predicted using spatial correlates of land cover, biotic and abiotic factors (pseudo R^2^ = 0.65). Foliar stoichiometric C:N ranges from 44.9 to 86.2 and is predicted using spatial correlates of EVI and biotic factors (pseudo R^2^ = 0.38). Foliar terpene (raw) ranges from 0.003 to 32.52 and is predicted using spatial correlates of biotic and abiotic factors (pseudo R^2^ = 0.26). Although these traits are predicted using different spatial correlates, emerging spatial patterns in trait variability suggest different processes are acting on trait expressions in different areas. For instance, high foliar C areas may relate to community type (land cover), forest structure (biotic), and topographic conditions (abiotic), however, patterns of C:N forest structure (biotic) and site productivity (EVI) indicate nutrient limitation areas with lower values have higher foliar nitrogen content. Moreover, foliar terpene patterns provide contours from which higher herbivore interactions results in increased terpene production. When overlaid with C and C:N we can gleam spatial patterns on the allocation of C to terpene production in terms of nutrient limitation constraints.

## 4. Discussion

Identifying spatial explicit drivers of foliar ESP traits may be a starting point to study ecosystem processes at the landscape extent. Our approach allows us to obtain spatially explicit estimates of heterogeneity through the development of foliar ESP trait distribution models (e.g., Fig. 5, Pickett and Cadenasso, 1995; Shen et al., 2011; Turner, 1989). Foliar ESP traits are often useful indicators of primary productivity, community structure, nutrient cycling, and trophic interactions (Brauer et al., 2012; Hunter, 2016). Here, we use differing combinations of spatially explicit covariates: land cover, productivity (EVI), biotic (forest structure: age, height, canopy closure), and abiotic (elevation, aspect, slope) factors, to identify which combinations of these factors drive ESP traits for our focal species at the landscape extent. In addition, we compare trait drivers across species to determine if there are commonalities. We find that not all traits, across species, are driven by the same spatial covariates. Although many studies have demonstrated community level coordination of foliar traits (Callis-Duehl et al., 2017; Descombes et al., 2017; Fyllas et al., 2020; Jiang et al., 2016), our findings suggest that trait spatial patterns are largely species specific. Thus, pattern-to-process relationships act at the species level to create landscapes of plant trait spatial heterogeneity and provides us with a new lens to evaluate landscape function. For instance, the spatial co-location of foliar resource convergence and divergence likely influence where, how, and why herbivores make foraging trade-offs decision between multiple forage species (Balluffi-Fry et al., 2020; Haynes & Cronin, 2004; Hunter, 2016). By developing spatial distribution models for multiple species and their traits (see Fig. 5 for an example) these maps can aid us in identifying resource hot spots of ecosystem services (Bernhardt et al., 2017; Lavorel et al., 2011; McClain et al., 2003), which in turn can inform herbivore foraging and pollinator (Filipiak, 2018) strategies and trade-off decisions (Shepard et al. 2013; see also Appendix 20 Fig A6 for a spatial correlation matrix of observed versus predicted ESP surfaces).

### 4.1. Foliar percent elemental traits

At the foliar percent elemental trait level, C, N, and P, we find mixed support for general patterns, as our results support species-specific spatial covariate trait relationships. For instance, abiotic covariates occurred more often as a top model, reinforcing a Humboltian perspective of plant distributions influence by soil and climate (Pausas & Bond, 2019). Other top ranked models, however, with biotic components, suggest that land cover type, site productivity, and forest structure have an influence on the spatial variability of foliar percent elemental traits. Across species, the EVI covariate did not occur in top ranked models for foliar percent carbon, although land cover, biotic and abiotic correlates did. Foliar percent C, N and P are often a useful measure of site level productivity, and EVI is a measure of productivity from space, however, a difference in scale here is likely why EVI is not a spatial driver of these foliar traits. Our results suggest that land cover and biotic factors of forest structure, likely have more of an influence on these foliar traits at the landscape extent (Rijkers et al., 2000). However, we did find that different combinations of EVI, biotic, and abiotic correlates influence foliar percent P at the trait level; suggesting that land cover type may not regulate phosphorus pathways. The weathering of rocks and soil particles contribute to soil P availability (i.e., EVI as a proxy for productivity/soil fertility) and P acquisition and nutrient uplift likely depends on competitive interactions determined by community types (i.e., biotic factors), and soil and water movement that facilitate anion and cation exchanges from soils particles to roots (Smith et al., 2000).

At the species level, general drivers of foliar percent C, N, P composition are more evident. For example, our models of (1) abiotic and (2) land cover, biotic and abiotic were the top models for foliar percent carbon in red maple and balsam fir and for lowbush blueberry and black spruce, respectively. This may suggest that between species with differing life histories that operate on different ends of the leaf spectrum (i.e., long lived versus seasonal foliar material); similar spatial predictors influence foliar percent carbon. Moreover, red maple foliar percent N content showed specificity to deciduous land cover and open canopy conditions, which may suggest increased N use efficiency in areas where deciduous leaf litter feedbacks ameliorate microbial community associated with plant functional types (Hobbie, 2015). These patterns provide evidence that biotic interactions may have important consequences for intraspecific variability of plant traits. Not all correlates within top models were, however, significant drivers. Notably, elevation and slope were important for species foliar percent carbon, supported by models with abiotic correlates. Together elevation and slope often have an influence on soil nutrient retention due to drainage properties (Müller et al., 2017). In addition, age classes (a biotic correlate) was important for black spruce foliar percent carbon, thus as the dominant tree species in this area, optimal carbon sequestration potential may occur under black spruce canopy community types across various seral stages (Dunn et al., 2009). We failed to find evidence to support models for foliar percent N and P of white birch. White birch is a clonal species with ramets that connect neighbouring individuals and facilitate the sharing of elemental resources to enhance collective nutrient use efficiencies (Bittebiere et al., 2019; Cornelissen and Cornwell, 2014). Ramet nutrient sharing, coupled with high plasticity of intraspecific variability in foliar percent elemental traits likely explain why we failed to detect a spatial signal with our covariates for white birch (Pyakurel and Wang, 2014; Wam et al., 2018). Overall, on the landscape, different drivers of foliar resource quality (i.e., C, N, and P), result in spatially heterogeneous species-specific resource hot spots. This may have far reaching implications for consumer dynamics and ecosystem processes (Haynes and Cronin, 2004; Wam et al., 2018).

### 4.2 Foliar quantity elemental traits

We only found support for drivers of foliar quantity elemental traits for two out of our five study species; balsam fir and lowbush blueberry. In each case, a single but different model explained all foliar quantity elemental traits. Collectively, these covariate combinations suggest that community type along with the structural properties of community conditions and abiotic factors highly determine the amount of foliar quantity C, N, and P resources. Across the landscape, these spatial covariates allow us to map the distribution of foliar quantity C, N, and P to detect areas of plant performance (i.e., optimal growth), resource abundance, and biogeochemical hot spots associated with nutrient uplift and storage (McClain et al., 2003; Tang et al., 2018). In addition, foliar quantity C is often related to leaf dry matter content, where increased dry matter correlates with decreased leaf palatability (Adler et al., 2014) and as such is a suggested driver of herbivore foraging trade-offs between quantity and quality (Champagne et al., 2018; Wam et al., 2018). The lack of evidence, however, to support foliar quantity elemental traits in our other study species constrains our ability to form generalizations of species spatial patterns and the processes that drive them, and as such suggests that these traits are either driven by different covariates or that inference may be limited to smaller spatial extents (Smithwick et al., 2003).

### 4.3 Foliar stoichiometric traits

Across species at the trait level, we have limited evidence to support generalizations of spatial foliar stoichiometric relationships. More notable are the foliar stoichiometric patterns that emerge at the species level. For instance, foliar C:P and N:P between balsam fir and red maple share similar predictors. However, for red maple, elevation and slope were determined to be key correlates, in comparison, aspect was a significant correlate for balsam fir. This suggests, that although these traits share similar predictors, the impact of these correlates differ, likely due to species and community level differences of nutrient co-limitation (Brauer et al., 2012). In contrast, lowbush blueberry and black spruce share a similar predictor for foliar N:P and similar responses to significant correlates of EVI, age class (i.e., biotic factor), elevation, and slope. Here, although, lowbush blueberry and black spruce occupy different ecological niches, they appear to respond to similar constraints of nutrient co-limitation, and thus may be nutrient limited under similar conditions. Similar to foliar percent and quantity elemental traits, we did not find evidence of a spatial covariate relationship for white birch foliar stoichiometric traits. Although communities are often spatially structured by nutrient co-limitation (Harpole et al., 2011), clonal strategies of ramet nutrient transfer may diminish these effects and as such constrain our ability to detect spatial predictors of foliar C:N, C:P, and N:P in white birch (Alpert, 1991; Li et al., 2004; Zhang and He, 2009). Collectively, this information is vital to informing resource hot spots, and mechanisms of nutrient co-limitation that structure biological communities (Gimona & van der Horst, 2007; Harpole et al., 2011). For instance, foliar N:P range maps for balsam fir and red maple provide nutrient use efficiency contours from which we can make spatial comparisons of species interactions that scale to the community structure level and aid us in identifying multi-species foliar resource hot spots. Moreover, by describing the spatial patchiness of resources we can inform herbivore foraging decisions and begin to make novel spatially explicit predictions associated with movement and behavioural trade-offs (Balluffi-Fry et al., 2020; Leroux et al., 2017; Rizzuto et al., 2019).

### 4.4. Foliar phytochemical traits

Across species, at the trait level we potentially have support to form generalization of geographic commonalities of foliar phytochemical traits. For all traits, except foliar sesquiterpene and phytochemical diversity, the biotic and abiotic model was determined to be an important spatial driver. This may suggest that structural properties of habitats (i.e., stand age, tree heights, and canopy conditions) and topographic conditions interact to determine foliar phytochemical traits. This is, to some extent expected, given that phytochemical traits are influenced by the spatial association of other species and their responses to the presence of herbivores (Champagne et al., 2018). On the island of Newfoundland, moose often forage on balsam fir and not black spruce (Gosse et al., 2011). Given the presented commonalities, consumption of balsam fir may elicit a non-consumptive phytochemical response in black spruce, thus further decreasing its potential palatability and aligning their foliar phytochemical composition (however, see Hussain et al., 2019).

At the species level, general patterns of foliar phytochemical trait correlates are less evident. Given the predominance of our phytochemical groups in both balsam fir and black spruce, we expected that similar spatial covariates should yield similar results between species. Our results, however, suggest foliar phytochemical traits exhibit species specificity to many different correlates. For instance, balsam fir and black spruce foliar terpene had differing predictors and differing significant correlates. Although some similarities between these two species exist, they are for traits on a different basis. For example, balsam fir foliar monoterpene on a biomass basis and black spruce foliar monoterpene on raw basis shared predictors; however, their response to specific correlates differed. For balsam fir, EVI as a remotely sensed proxy for productivity correlates to foliar terpene and monoterpene traits, suggesting optimal nutrient conditions may invoke a strong defence position (Lindroth et al., 2002). However, there are potential confounding effects. Increased phytochemical production, in species with constituent strategies (i.e., maintained baseline phytochemical production), may occur in response to the presence and or interaction of an herbivore (Kessler, 2015), which in turn influence top-down nutrient dynamics (Hunter, 2016) in positive or negative ways depending on the soil condition and litter feedbacks (Hemming & Lindroth, 1999; Hobbie, 2015). As well, when we relativized phytochemical variables on a biomass basis, for balsam fir, support for foliar terpene, monoterpene, and monoterpenic ester traits was explained by the same combination of spatial covariates; abiotic and biotic. In contrast, we had no evidence to support spatial relationships of black spruce foliar phytochemical traits on a biomass basis. More notably, between the two species, abiotic covariates appear to influence foliar sesquiterpene. Here, the intraspecific variability of phytochemical groups and measure of compound diversity are often used as a proxy to indicate plant-herbivore interactions, herbivore diversity, and trophic specialization (Richards et al., 2015). From our results, we find evidence to map phytochemical terpene groups and diversity, with some similarities in covariate specificity between two species with similar life histories. The spatial variability of foliar phytochemical composition provides us with a spatially explicit way to unravel the consequences and species interactions of herbivore foraging patterns with links to nutrient cycling processes (i.e., soil trampling, nutrient transfer, and changes in plant species composition Champagne et al., 2018; Gosse et al., 2011; Hunter, 2016).

### 4.5 Implications of ESP spatial trait distributions beyond the boreal

Foliar ESP traits represent a common currency of species (Elser & Hamilton, 2007). These traits are often used as indicators for differing ecological conditions with consequences that reach across levels of biological organization (Fajardo & Siefert, 2018). For instance, global patterns of N and P are associated with latitudinal gradients, with northern plants having higher concentrations of N and P related to plants at the equator (Reich & Oleksyn, 2004). By identifying the spatially explicit drivers of foliar N and P, we can map resource hot spots and compare how the distribution of these resources influence primary production (Smithwick et al., 2003), nutrient uplift (Jobbágy & Jackson, 2004), herbivore space use and forage selection (Duparc et al., 2020), and community assembly processes (Harpole et al., 2011; Jung et al., 2010) in different ecosystems. Moreover, we can begin evaluate the spatial flows of elements across the landscape (Shen et al., 2011). Indeed, many studies have identified spatial drivers of foliar ESP traits in differing ecosystems (see Table 1 for a non-exhaustive list of studies) however, a spatially explicit approach is needed to derive predictions from which we can map these resource distributions and obtain estimates of spatial heterogeneity.

## 5. Conclusion

By identifying spatially explicit covariates for foliar ESP traits at the species level, we can develop distribution models of intraspecific trait variability across a boreal landscape (for an example see Fig. 5). These distribution models, allow us to explore the consequences of trait spatial heterogeneity and the processes that drive them with implications for landscape functionality (Harvey et al., 2019). For example, we can test hypotheses about herbivore resource selection across scales (Balluffi-Fry et al., 2020), infer landscape functionality via pattern and process relationships (Turner, 1989), or explore how the spatial distribution of matter and energy feedbacks on landscape structure with implications for the management of biogeochemical processes (Lovell & Johnston, 2009; Shen et al., 2011). In addition, our work described here may be of use to carbon modelling approaches which largely focus on sequestration and storage, or Net Ecosystem Production (NEP), and overlook carbon dynamics at the interface of ecological interactions (Schmitz et al., 2018). Knowing how much carbon is sequestered, lost through respiration, or through pathways of non-photosynthetic carbon, foliar carbon reabsorption, and foliar carbon loss through consumptive activities allows for the refinement of carbon cycling models (Dirnböck et al., 2020). Given the importance of the circumboreal in carbon cycles, our work here can help understand how carbon dynamics may manifest in other parts of the boreal. Here, we investigated the drivers of foliar ESP traits for commonly occurring, geographically widespread boreal species using accessible spatial covariates. We found some geographic commonalities in spatial covariates at the trait and species level from which we can make generalities about physiological links to ecosystem processes and landscape function (Hobbie, 2015; Li et al., 2004; McClain et al., 2003; Poorter and Bongers, 2006). There are specificities in spatial predictors at the species level that suggest plants respond differently to environmental conditions and that ideas of resources hot spots are likely species specific. How different species of plants respond in different parts of the world merits further work like this that combines landscape ecology, spatially modelling, and plant stoichiometry.

## Acknowledgements

This research was funded by the Government of Newfoundland and Labrador Centre for Forest Science Innovation (CFSI); Memorial University of Newfoundland SEEDS funding to SJL, EVW, YFW; Mitacs Accelerate Grant to YFW, SJL and EVW; Canada Foundation for Innovation funding to YFW, and the Natural Sciences and Engineering Research Council of Canada (Discovery Grant to YFW). In-kind support was provided by Parks Canada – Terra Nova National Park and the CFSI, with thanks to Janet Feltham and Blair Adams. In addition, we would also like to thank the Landscape Ecology and Spatial Analysis lab at Memorial University of Newfoundland. Thank you to three anonymous reviewers for helpful comments that helped to substantially improve an earlier version of the manuscript.

## Funding

This research was funded by the Government of Newfoundland and Labrador Centre for Forest Science and Innovation to YFW (Grant #221273), SJL (Grant #221274), and EVW (Grant #221275), Government of Newfoundland and Labrador Innovate NL Leverage R&D to EVW & SJL (Grant #5404.1884.102) and Ignite R&D to SJL (Grant #5404.1696.101) programs, Mitacs Accelerate Graduate Research Internship program to YW, EVW, & SJL (Grant #IT05904), the Canada Foundation for Innovation John R. Evans Leaders Fund to EVW & SJL (Grant #35973), and a Natural Science and Engineering Research Council Discovery Grant to YFW (Grant #RGPIN-2015-05799).

## Conflicts of interest

The authors have no conflicts of interest to declare that are relevant to the content of this article.

## Ethics approval

n/a

## Consent to participate

n/a

## Consent for publication

there are no restrictions to publish

## Availability of data and material

The datasets and associated R code used to conduct research presented in this manuscript are available on Figshare.com at https://doi.org/10.6084/m9.figshare.11911455.v1. The Forest Resource Inventory datasets provided by the Provincial Government of Newfoundland and Labrador and the Federal Government of Canada may be made available upon request, however additional permission through these agencies may be required.

## Code availability

The R code used to conduct research presented in this manuscript are available on Figshare.com at https://doi.org/10.6084/m9.figshare.11911455.v1.

## Appendices

## Appendix 1

Detailed description of Fig. 1: the roadmap of our methods. Our study area (a) is location on the eastern side of the island of Newfoundland, North America, Canada, as shown by the outlined area. Generally, bounded between the 47^th^ and 48^th^ latitude this biogeographical area is composed of boreal forest conditions primarily dominated by intermediate-aged, closed canopy, forest stands of black spruce (*Picea mariana*), balsam fir (*Abies balsamea*), white birch (*Betula papyrifera*), and trembling aspen (*Populus tremuloides*) (Ecological Stratification Working Group, 1996; South, 1983). Within this area we set up four chronosequenced grids, consisting of connected meandering transects. Age classes and grid layout shown in panel b. Grids were originally designed for snowshoe hare (*Lepus americanus*) trapping and to allow us to relate foliar resource quality to hare home range size and ecology. Each grid is comprised of 50 sampling locations, spaced equally apart by 75 m with closer sample location rounding the corners (b). At each sample location we set up 22.6 m diameter circular plots (c). Within each plot we collected density estimates for each of our study species along a 22.6 m long and 1 m wide shrub belt transect (c/d). Moving in a north to south direction, along the belt, for each of our study species encountered we measured their height and basal diameter, and the distance at which it was encountered, for a maximum of five individuals per height class: 0-50 cm, 51-100 cm, 101-150 cm, and 151-200 cm, denoted as A, B, C, and D respectively (d). We restricted our sampling to species within 0-2 m heights (d) as these individuals represent the available forage for common boreal herbivores, moose (*Alces alces*) and snowshoe hare. Within each plot, starting in the NE corner (e), we moved in a clockwise direction and collected foliar samples of our study species, as well we measured their height and basal diameter. In panel e, we use coded names for our study species, balsam fir (ABBA), red maple (ACRU), white birch (BEPA), black spruce (PIMA), and lowbush blueberry (VAAN); see Appendix 2 for a description of our study species. We collect foliar material for our study species until we had a sufficient sample size of approximately 10-20 g. Using foliar samples for each of our study species, we combined representative units of foliar material until a wet weight sample of 10 g and 4 g was amassed – the amount required for elemental and phytochemical analysis, respectively. At the Agriculture Food Lab (AFL) at the University of Guelph Ontario, Canada the carbon and nitrogen composition of foliar material determined using an Elementar Vario Macro Cube. Foliar phosphorus content was determined using a microwave acid digestion CEM MARSxpress microwave system and brought to volume using Nanopure water. The clear extract supernatant was further diluted by 10 to accurately fall within calibration range and reduce high level analyte concentration entering the inductively coupled plasma mass spectrometry (ICP-MS) detector (Poitevin, 2016). This provides us with a measure of percent foliar C, N, and P. At the Laboratorie PhytoChemia Inc in Quebec, Canada, the phytochemical composition of balsam fir and black spruce foliar samples were determined using a gas chromatography solvent extraction with an internal standard and a correction factor (Cachet et al., 2016). This procedure produced mg/g measures of individual terpene compounds, see Appendix 7 Table A2 for a complete list of identified terpene compounds and groups. In addition, along the periphery of our study grids and outside of the sample plots, in randomly selected locations we collected all new growth foliar material for each of our study species from approximately 50 individuals, the number of samples distributed across the height classes listed above (f). As well, we measured the height and basal diameter for each individual sampled. The foliar material was dried, providing a measure of biomass from which we fit linear allometric models using covariates of height and basal diameter (f). Using coefficient estimates from our allometric models we predicted biomass estimates for our study species per height class from shrub belt measurements. In the few instances where we had obtained foliar samples but did not encounter individuals on the shrub belt we augmented the total number of individuals per height class as the total number of foliar samples in that height class. We subsequently summed biomass estimates per height class for each of our study species and divided this measure by the area of the circular plot (401.15 m^2^) to get a density estimate. We then multiplied biomass by density for each height class to get a species biomass estimate, which was summed together, providing a plot level biomass estimate per species. To obtain elemental quantity estimates we divided biomass by the plot area multiplied by the foliar percentage of carbon, nitrogen, and phosphorus. As well, we did the same for phytochemicals to obtain a plot level biomass basis estimate of foliar phytochemicals. To determine stoichiometric ratios, we divided quantity C, N, and P estimates by their corresponding molar mass and then divided the resulting value together to get foliar C:N, C:P, and N:P for each study species. Using response variables of foliar percent elemental, quantity elemental, stoichiometric, and phytochemical we constructed sixteen plausible model combinations with spatially explicit covariates of land cover, productivity, abiotic, and biotic factors and used Akaike Information Criterion to determine plausible explanations (g). We then assessed top models and extracted coefficient estimates for use in constructing distribution models of foliar elemental, stoichiometric, and phytochemical traits which provides us surfaces to inform landscape function (g).

## Appendix 2

**Figure A1.**
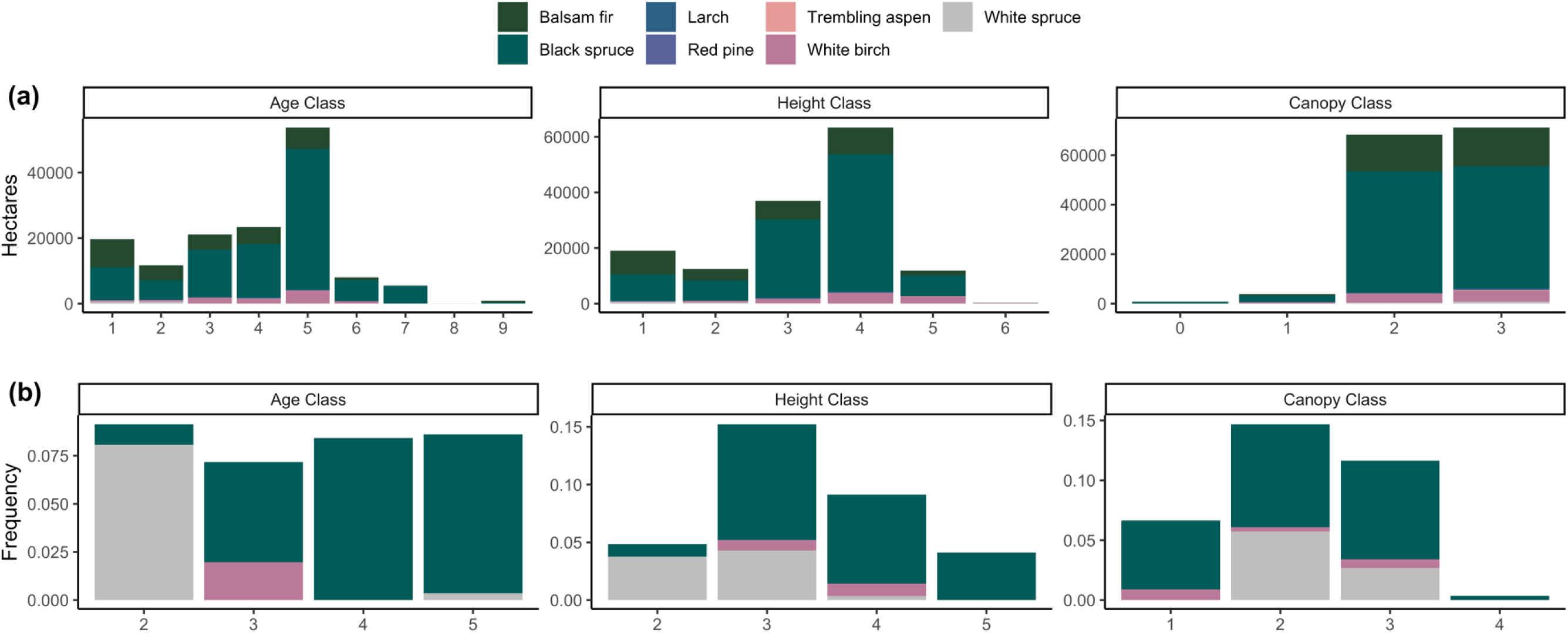
The top (a) shows the total number of hectares for each dominant species forest type within our landscape area of interest, for stand metrics of age, height, and canopy class. Age class codes represent 20-year intervals ranging from 1 (0-20 years) to 9 (161+ years). Height class codes represent 3.5 m intervals of tree heights ranging from 1 (0-3.5 m) to 6 (15.6-18.5 m). Canopy class codes represent 25 % intervals of canopy closeness where 0 indicates a regenerating stand that is 100 % closed and 4 indicates a 10 −25% closed canopy conditions. The bottom (b) shows the frequency in which these dominant species forest stands were sampled for foliar ESP traits of our study species. Here we show that although our sampling design is not ideal for spatial distribution modelling, we sampled within representative units of forest types available on the landscape, thus strengthening our inference for the spatial distribution of foliar ESP traits on this landscape.

## Appendix 3

**Table A1.** A complete list of phytochemical compounds and classes for terpenes identified in balsam fir and black spruce foliar samples. Only common terpene groups between these two coniferous species were used: terpene (includes all compounds identified), monoterpene, monoterpenic alcohol, monoterpenic ester, sesquiterpene, and diversity (computed using all compounds identified).

| Group | Balsam fir | Black spruce |
| --- | --- | --- |
|  | *Chemical Name* | *Chemical Name* |
| Monoterpene | Tricyclene | Tricyclene |
| | α-Pinene | α-Pinene |
|  | Camphene | Camphene |
| | β-Pinene (main) + Sabinene | β-Pinene (main) + Sabinene |
|  | Myrcene | Myrcene |
| | Δ-Carene | Δ-Carene |
| | Limonene (main) + β-Phellandrene | α-Phellandrene |
| | γ-Terpinene | Limonene (main) + 1,8-cineole |
|  | Terpinolene | Terpinolene |
| Monoterpenic | Linalool | Linalool |
| Alcohol | Camphene hydrate | Camphene hydrate |
|  | Borneol | Citronellol |
| | α-Terpineol | α-Terpineol |
|  | Thymol | --- |
| **Monoterpenic Ester** | Bornyl acetate | Unknown "PIMA 6" |
|  |  | Bornyl acetate (main) + Isobornyl acetate |
|  |  | endo-Fenchyl acetate |
|  |  | trans-Pinocarvyl acetate |
|  |  | cis-Piperityl acetate |
|  |  | Geranyl acetate |
| **Sesquiterpene** | Longifolene | β-Elemene |
| | β-Caryophyllene | β-Caryophyllene |
| | α-Humulene | α-Humulene |
| | (E)-β-Farnesene | Germacrene D |
| | Unknown sesquiterpene | γ-Cadinene (main) + Cubebol |
| | α-Muurolene | α-Muurolene |
| | (Z)-α-Bisabolene | δ-Cadinene |
| | β-Bisabolene | Unknown sesquiterpene |
| | (E)-α-Bisabolene | (E)-α-Bisabolene |
| **Sesquiterpenic alcohol** | -- | Germacrene D-4-ol |
| | | τ-Cadinol + τ-Muurolol (approx 1:1) |
| | | α-Cadinol |
|  |  | Oplopanone |
| **Sesquiterpenic ether** | Caryophyllene oxide |  |
| **Monoterpenic aldehyde** | -- | α-Campholenal |
| **Monoterpenic ketone** | Piperitone | -- |
| **Maltol** | Maltol | -- |
| **Oxygenated sesquiterpene** | -- | Unknown "PIMA 18" |
| **Unknown** | -- | Unknown "PIMA 9" |

## Appendix 4

**Figure A2.**
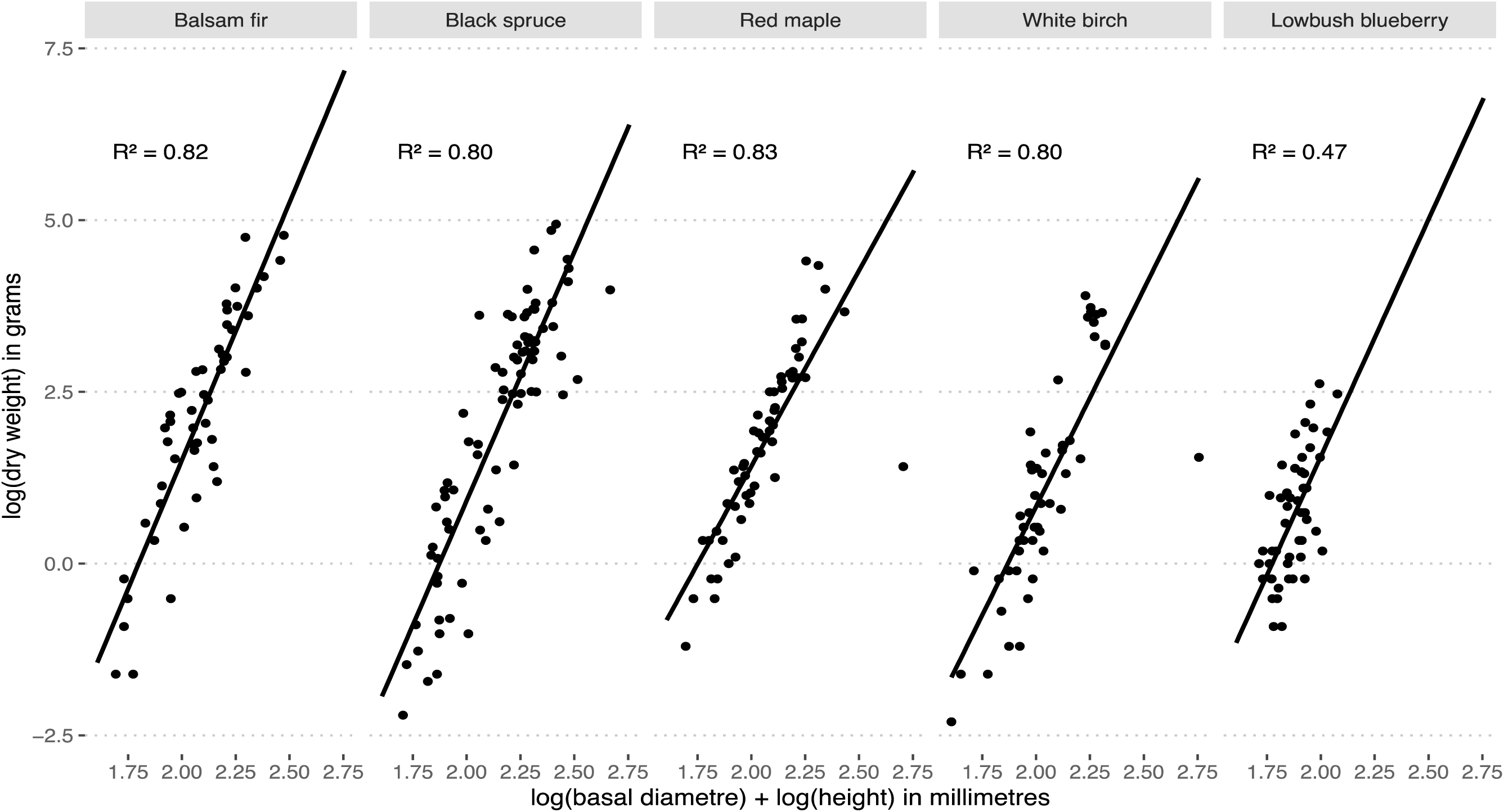
Allometric modelling of biomass in terms of basal diameter and height for each of our study species, balsam fir, black spruce, red maple, white birch, and lowbush blueberry. The goodness of fit (adjusted R^2^) is superimposed on each species regression plot.

## Appendix 5

**Table A2.**
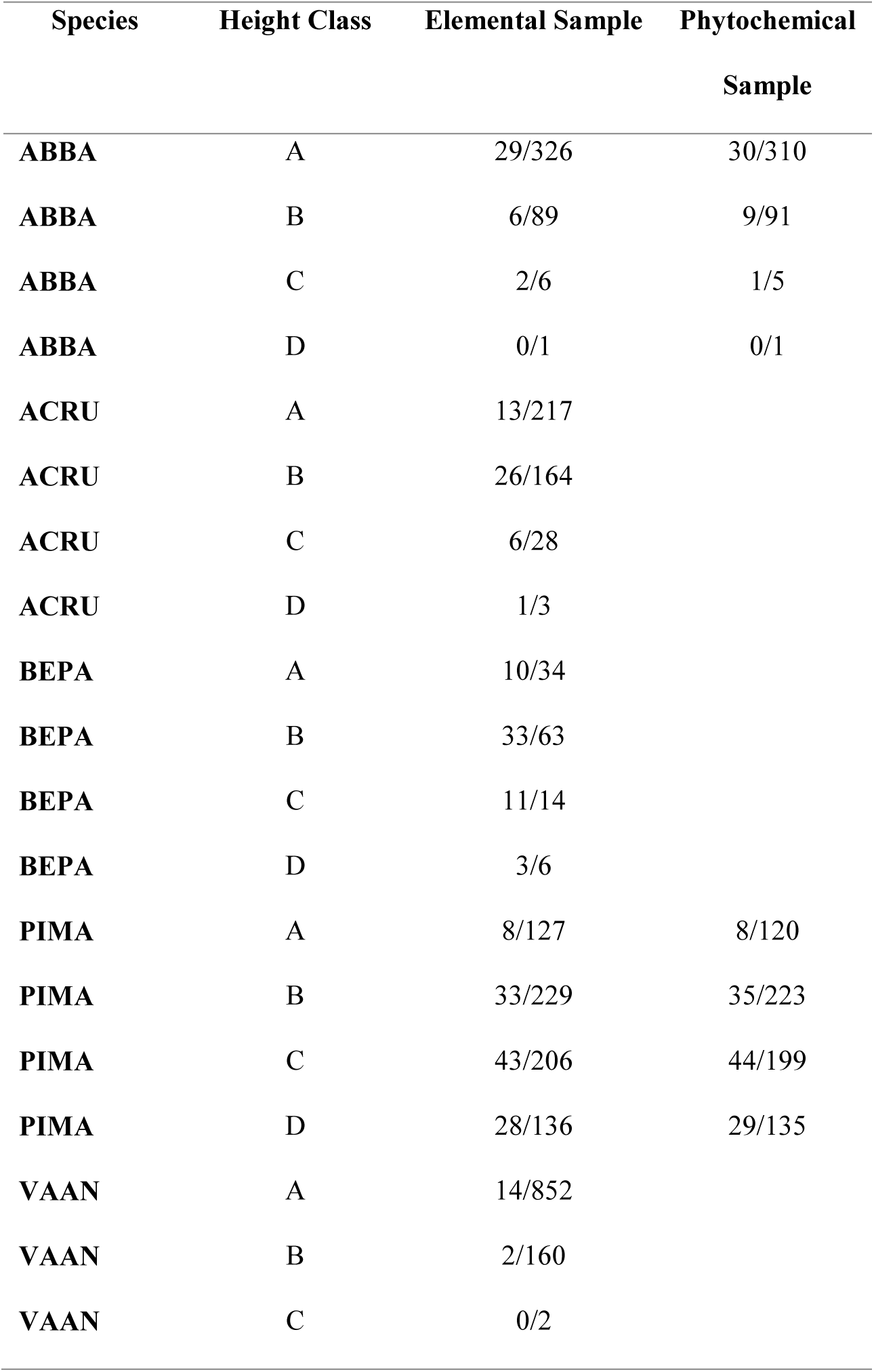
The number of individuals that we augmented using foliar samples to obtain density measures when individuals of that species were not encountered on the shrub belt. Numbers are shown for each species per height class relative to the total number of individuals used in that height class. Height class is coded as A = 0-50 cm, B = 51-100 cm, C = 101-150 cm, and D = 151-200 cm.

| Species | Height Class | Elemental Sample | Phytochemical<br>Sample |
| --- | --- | --- | --- |
| ABBA | A | 29/326 | 30/310 |
| ABBA | B | 6/89 | 9/91 |
| ABBA | C | 2/6 | 1/5 |
| ABBA | D | 0/1 | 0/1 |
| ACRU | A | 13/217 |  |
| ACRU | B | 26/164 |  |
| ACRU | C | 6/28 |  |
| ACRU | D | 1/3 |  |
| BEPA | A | 10/34 |  |
| BEPA | B | 33/63 |  |
| BEPA | C | 11/14 |  |
| BEPA | D | 3/6 |  |
| PIMA | A | 8/127 | 8/120 |
| PIMA | B | 33/229 | 35/223 |
| PIMA | C | 43/206 | 44/199 |
| PIMA | D | 28/136 | 29/135 |
| VAAN | A | 14/852 |  |
| VAAN | B | 2/160 |  |
| VAAN | C | 0/2 |  |

## Appendix 6

Our spatial resolution was constrained by our coarsest dataset, Landsat 8, i.e., 30 m resolution. In ArcGIS, we resampled elevation and our Digital Elevation Model from a 20 m to a 30 m resolution. The Forest Resource Inventory vector dataset was rasterized at a 30 m resolution.

### Enhanced Vegetation Index (EVI)

Landsat 8 satellite imagery was acquired from the Earth Resources Observation (EROS) and Science Centre Science Processing Architecture (ESPA). There were three Landsat 8 scenes available during our 2016 sampling time period; June 28, August 15, and September 16, 2016 with 0.46%, 20.18%, 4.39% land cloud cover respectively. As a standard product, Landsat 8 acquisitions contain a preprocessed EVI surface reflectance scene. Newfoundland boreal forest demonstrably receives a greater amount of precipitation and experiences shorter growing seasons due to Atlantic Ocean influence creating colder climatic conditions compared to continental boreal forest conditions (South, 1983). Under these conditions, the EVI as a measure of biological productivity performs better than the Normalized Difference Vegetation Index which commonly saturates early in the season and does not account for the structural complexity of vegetative canopies (Muraoka et al., 2013; Requena-Mullor et al., 2017; Waring et al., 2006). Using the Landsat Quality Assurance ArcGIS toolbox, publicly accessible software from the U.S. Geological Survey, we extracted the following cloud coded bits from the pixel QA band: cloud shadow, snow, cloud, high cloud confidence and high cirrus confidence (Jones et al., 2013; U.S. Geological Survey, 2017). Using the ‘Extract by Mask’ ArcGIS function we removed cloudy pixels from our EVI scenes. In R, we rescaled EVI scenes by dividing by 0.0001. Using the ‘approxNA’ function from the ‘raster’ R package (Hijmans, 2020), we computed a linear interpolation across our temporal scenes to fill cloud removed pixels, see Appendix 7 Fig. 3, for before and after interpolation maps and pixel histograms. We average our temporal EVI scene to obtain an estimated seasonal measure of productivity. Using the ‘raster.transformation’ function from the ‘spatialEco’ R package, we standardized the EVI annual productivity scene by subtracting the scene mean from each pixel and dividing by the scene standard deviation (Evans, 2020).

### Elevation, Aspect, Slope and Land Cover

A Canadian Digital Elevation Model (DEM) was retrieved from Natural Resources Canada. Using ArcGIS, we combined DEM images together to create a seamless raster. In ArcGIS, using the ‘Clip’ function we limited our DEM raster to our AOI. Using the ‘terrain’ function from the ‘raster’ R package we constructed aspect and slope raster. We normalized our aspect raster by replacing any value > 180 by subtracting −180 (e.g., an aspect of 240 is now an aspect of 60; changing the scale from 0-360 to 0-180). We used the base R ‘subs’ function with a legend of corresponding values to normalize the aspect raster. As we did for the EVI raster, we standardized elevation, aspect, and slope rasters using the ‘raster.transformation’ function from the ‘spatialEco’ R package. In addition, we used the freely accessible Commission for Environmental Cooperation Land Cover dataset; derived from Landsat images, to obtain categorical values of forest type: coniferous, deciduous, mixed coniferous and deciduous.

### Forest Resource Inventory

our AOI covers a national park, Terra Nova National Park (TNNP) and public land. Spatial information regarding forest stand attributes, Forest Resource Inventory (spatial vector), were supplied to us from two sources: Parks Canada and the Provincial Government of Newfoundland and Labrador. Using unique forest polygon identifiers, we attributed spatial covariates to the FRI datasets (attributes also contained non-interest covariates). To construct a seamless FRI layer across our AOI we combined the two sets of Forest Resource Inventory together. In ArcGIS, using the ‘clip’ function we constrained the geographic extents of the two FRI datasets to our AOI; to alleviate spatial data processing time. Using the ‘erase’ function in ArcGIS we removed any spatially overlapping boundaries between the two FRI datasets. Using the ‘merge’ ArcGIS function we create a single FRI dataset by spatially joining the two FRI datasets together. In R, we subset the FRI dataset to only include covariates of interest: forest stand age class, height class, and crown density – categorical properties that likely influence growing conditions and thus the elemental and phytochemical properties of our plants. In R, we further cleaned the FRI dataset by removing any non-intention ‘white space’ in the text of the categorical data. For each co-variate we extracted unique values and re-coding text values as integers. Using the ‘rasterize’ function from the ‘raster’ R package, we convert our FRI vector data into a raster for each co-covariate, using the integer values as a coded legend for our categories. In addition, we created binary layers for each factor in the age class, height class, and crown density variables. Binary layers were used when model average estimates were extracted as the predict function in the ‘raster’ package is limited to single model objects.

### Inference Mask

Using species composition codes derived from the FRI dataset for each of our sample points, we create a vector mask of forest polygons types for which we have spatial inference. These codes represent community types dominated by either black spruce, white spruce, and white birch. In R, we used the ‘mask’ function from the ‘raster’ package to clip spatial covariate surfaces.

### Spatial Data Extraction

At each sample location, using the ‘extract’ function from the ‘raster’ R package we spatially extracted pixel values from each of our raster datasets: elevation, aspect, slope, and land cover. We used the ‘intersect’ function from the ‘raster’ R package to extract polygon forest stand attributes from the FRI dataset: age class, height class, and crown density. At some sample locations the FRI was either inaccurate or our sample location was within a wetland type area with no attributes. For these instances, we attributed our sample locations with the values from the closest forest stand polygon. In total there were 14, 3, and 5 incorrect spatial designations for age class, height class, and crown density.

## Appendix 7

**Figure A3.**
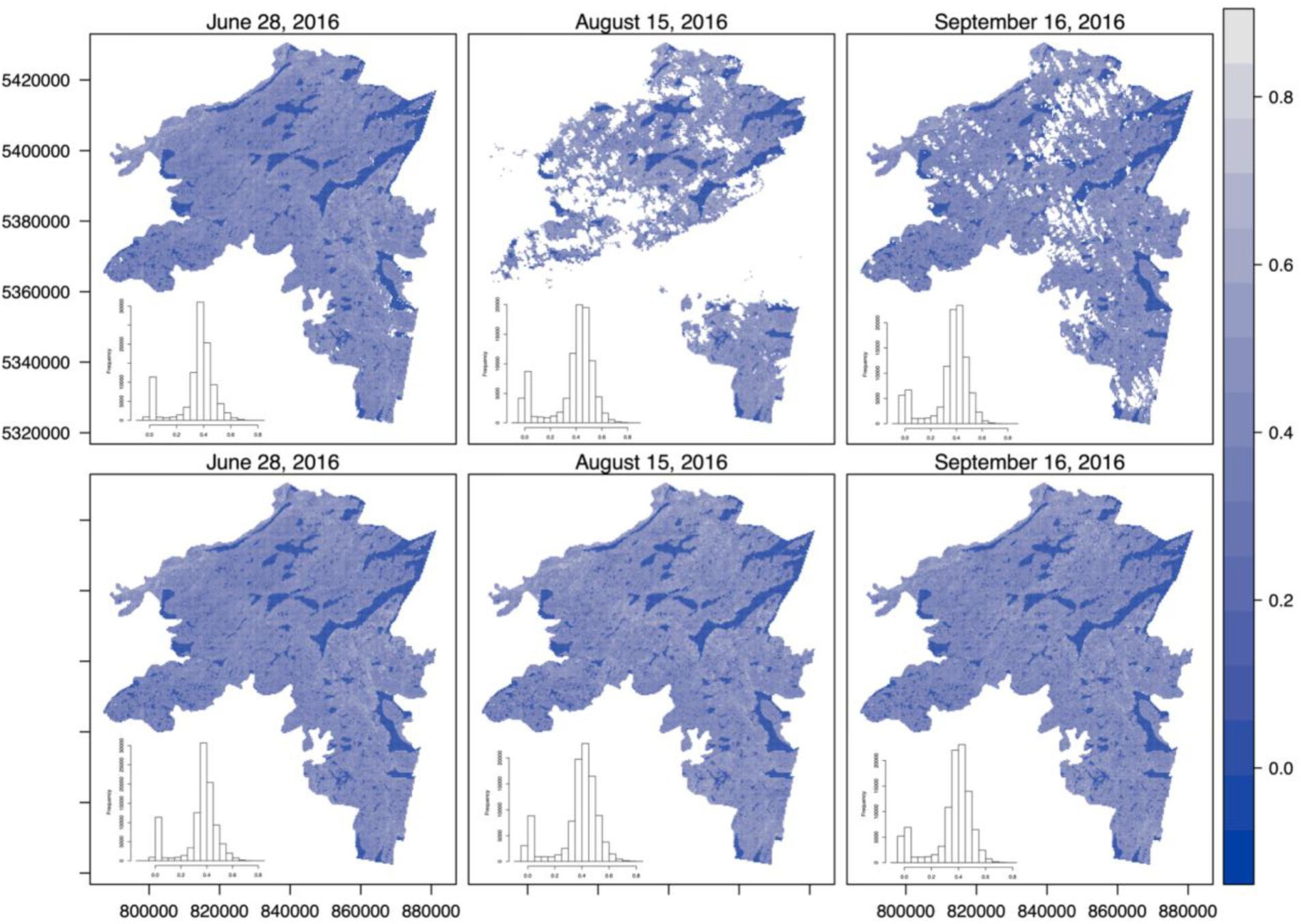
Using the ‘approxNA’ function from the ‘raster’ package in R, we performed a linear temporal interpolation to determine pixel values for areas of cloud cover for our three Enhanced Vegetation Index scenes, June 28, August 15, and September 16, 2016. The top panel shows each scene before interpolation and the bottom panel shows each scene after interpolation. Accompanying histograms are provided for each EVI scene, demonstrating the change in pixel value distribution after interpolation.

## Appendix 8

**Table A3.** AIC_c_ results for foliar elemental (percent and quantity), stoichiometric, and phytochemical traits. Explanatory variables include land cover, EVI, biotic, and abiotic. Land cover is a categorical variable with three factor levels which include coniferous, deciduous, and mixed. EVI is the Enhanced Vegetation Index and performs better than NDVI (Normalized Difference Vegetation Index) under wet conditions. Our biotic variable represents forest structural conditions and is comprised of three variables, age class, height class, and canopy density, each containing four factors levels of increasing age, height, and canopy density. Abiotic is comprised of three continuous variables for elevation, aspect, and slope. Results are shown for models within 2 delta AIC_c_, K is the number of parameters, LL represents the model log likelihood, ΔAIC_c_ for the interpretation of model ranking, ωAIC_c_ for model weights, and R^2^ is presented as Efron’s goodness of fit. Pretending variables are denoted with an asterisk and were removed from any model averaging. Biomass basis phytochemical models are identified with (bm).

| Species | Explanatory Variables | K | LL | ΔAIC<sub>c</sub> | ωAIC<sub>c</sub> | R<sup>2</sup> |
| --- | --- | --- | --- | --- | --- | --- |
| **Elemental: percent carbon** |  |  |  |  |  |  |
| **ABBA** | Abiotic | 5 | -47.96 | 0.00 | 0.52 | 0.19 |
|  | EVI* + Abiotic | 6 | -47.36 | 1.08 | 0.31 | 0.20 |
| **ACRU** | Abiotic | 5 | -51.43 | 0.00 | 0.37 | 0.09 |
|  | EVI* + Abiotic | 6 | -50.54 | 0.52 | 0.28 | 0.11 |
| **BEPA** | Biotic | 10 | -72.14 | 0.00 | 0.48 | 0.31 |
| **PIMA** | Biotic + Abiotic | 14 | -202.13 | 0.00 | 0.38 | 0.64 |
|  | Land Cover + Biotic + Abiotic | 16 | -199.88 | 0.42 | 0.31 | 0.65 |
| **VAAN** | Land Cover + Biotic + Abiotic | 16 | -92.50 | 0.00 | 0.44 | 0.37 |
|  | Land Cover + EVI* + Biotic + Abiotic | 17 | -91.78 | 1.06 | 0.26 | 0.37 |
| **Elemental: percent nitrogen** |  |  |  |  |  |  |
| **ABBA** | Land Cover + EVI + Abiotic | 8 | 48.53 | 0.00 | 0.62 | 0.29 |
| **ACRU** | Land Cover + Biotic | 13 | 20.02 | 0.00 | 0.36 | 0.36 |
|  | Land Cover + EVI* + Biotic | 14 | 21.05 | 0.75 | 0.25 | 0.37 |
| **BEPA** | Intercept | 2 | -45.13 | 0.00 | 0.41 | 0.00 |
|  | Abiotic | 5 | -42.47 | 1.43 | 0.20 | 0.07 |
|  | EVI | 3 | -44.96 | 1.85 | 0.16 | 0.00 |
| **PIMA** | EVI + Biotic | 12 | 112.91 | 0.00 | 0.68 | 0.36 |
| **VAAN** | EVI | 3 | 33.08 | 0.00 | 0.45 | 0.07 |
|  | Land Cover + EVI | 5 | 34.82 | 0.76 | 0.31 | 0.09 |
| **Elemental: percent phosphorus** |  |  |  |  |  |  |
| **ABBA** | Abiotic | 5 | 199.02 | 0.00 | 0.52 | 0.10 |
| **ACRU** | EVI | 3 | 134.27 | 0.00 | 0.39 | 0.05 |
|  | EVI + Abiotic | 6 | 136.95 | 1.35 | 0.20 | 0.10 |
| **BEPA** | Intercept | 2 | 94.94 | 0.00 | 0.51 | 0.00 |
|  | EVI | 3 | 95.44 | 1.19 | 0.28 | 0.01 |
| **PIMA** | Biotic + Abiotic | 14 | 345.69 | 0.00 | 0.42 | 0.28 |
|  | EVI* + Biotic + Abiotic | 15 | 346.63 | 0.57 | 0.32 | 0.29 |
| **VAAN** | EVI + Biotic | 12 | 419.42 | 0.00 | 0.60 | 0.47 |
|  | Biotic | 11 | 417.36 | 1.78 | 0.25 | 0.46 |
| **Elemental: quantity carbon** |  |  |  |  |  |  |
| **ABBA** | Biotic + Abiotic | 14 | -158.93 | 0.00 | 0.68 | 0.35 |
| **ACRU** | Intercept | 2 | -190.97 | 0.00 | 0.60 | 0.00 |
|  | EVI | 3 | -190.84 | 1.87 | 0.23 | 0.00 |
| **BEPA** | Intercept | 2 | 10.10 | 0.00 | 0.60 | 0.00 |
| **PIMA** | Intercept | 2 | -497.69 | 0.00 | 0.43 | 0.00 |
|  | Abiotic | 5 | -495.20 | 1.34 | 0.22 | 0.03 |
| **VAAN** | Land Cover + Abiotic | 7 | -444.06 | 0.00 | 0.56 | 0.19 |
| **Elemental: quantity nitrogen** |  |  |  |  |  |  |
| **ABBA** | Biotic + Abiotic | 14 | 227.46 | 0.00 | 0.68 | 0.35 |
| **ACRU** | Intercept | 2 | 121.64 | 0.00 | 0.58 | 0.00 |
|  | EVI | 3 | 121.87 | 1.67 | 0.25 | 0.01 |
| **BEPA** | Intercept | 2 | 251.05 | 0.00 | 0.62 | 0.00 |
| **PIMA** | Intercept | 2 | 146.79 | 0.00 | 0.38 | 0.00 |
|  | Abiotic | 5 | 149.56 | 0.79 | 0.25 | 0.03 |
|  | EVI | 3 | 146.97 | 1.74 | 0.16 | 0.00 |
| **VAAN** | Land Cover + Abiotic | 7 | 166.00 | 0.00 | 0.54 | 0.18 |
|  | Land Cover + EVI* + Abiotic | 8 | 166.19 | 1.84 | 0.22 | 0.19 |
| **Elemental: quantity phosphorus** |  |  |  |  |  |  |
| **ABBA** | Biotic + Abiotic | 14 | 449.29 | 0.00 | 0.53 | 0.32 |
| **ACRU** | Intercept | 2 | 334.15 | 0.00 | 0.61 | 0.00 |
|  | EVI | 3 | 334.24 | 1.96 | 0.23 | 0.00 |
| **BEPA** | Intercept | 2 | 433.74 | 0.00 | 0.62 | 0.00 |
| **PIMA** | Intercept | 2 | 470.41 | 0.00 | 0.48 | 0.00 |
| **VAAN** | Land Cover + Abiotic | 7 | 581.16 | 0.00 | 0.72 | 0.19 |
| **Stoichiometric: C:N ratio** |  |  |  |  |  |  |
| **ABBA** | Land Cover + EVI + Abiotic | 8 | -364.16 | 0.00 | 0.72 | 0.31 |
| **ACRU** | Land Cover + Biotic | 13 | -267.39 | 0.00 | 0.40 | 0.39 |
|  | Land Cover + EVI* + Biotic | 14 | -266.49 | 0.99 | 0.24 | 0.40 |
| **BEPA** | Intercept | 2 | -291.69 | 0.00 | 0.48 | 0.00 |
|  | EVI | 3 | -291.22 | 1.25 | 0.26 | 0.01 |
| **PIMA** | EVI + Biotic | 12 | -559.84 | 0.00 | 0.70 | 0.38 |
| **VAAN** | EVI | 3 | -539.51 | 0.00 | 0.44 | 0.09 |
|  | Land Cover + EVI | 5 | -537.79 | 0.79 | 0.30 | 0.11 |
| **Stoichiometric: C:P ratio** |  |  |  |  |  |  |
| **ABBA** | Abiotic | 5 | -755.06 | 0.00 | 0.63 | 0.14 |
| **ACRU** | Abiotic | 5 | -665.73 | 0.00 | 0.26 | 0.07 |
|  | Intercept | 2 | -669.06 | 0.07 | 0.25 | 0.00 |
|  | EVI + Abiotic | 6 | -664.89 | 0.61 | 0.19 | 0.09 |
|  | EVI | 3 | -668.32 | 0.75 | 0.18 | 0.02 |
| **BEPA** | Intercept | 2 | -568.29 | 0.00 | 0.49 | 0.00 |
|  | EVI | 3 | -567.62 | 0.85 | 0.32 | 0.02 |
| **PIMA** | Biotic + Abiotic | 14 | -1095.54 | 0.00 | 0.59 | 0.30 |
|  | EVI* + Biotic + Abiotic | 15 | -1095.19 | 1.75 | 0.25 | 0.30 |
| **VAAN** | Biotic | 11 | -1189.78 | 0.00 | 0.40 | 0.33 |
|  | EVI + Biotic | 12 | -1188.63 | 0.03 | 0.39 | 0.34 |
| **Stoichiometric: N:P ratio** |  |  |  |  |  |  |
| **ABBA** | Abiotic | 5 | -331.11 | 0.00 | 0.31 | 0.08 |
|  | EVI + Abiotic | 6 | -330.06 | 0.18 | 0.28 | 0.10 |
|  | Intercept | 2 | -334.87 | 0.99 | 0.19 | 0.00 |
| **ACRU** | EVI + Abiotic | 6 | -324.45 | 0.00 | 0.54 | 0.17 |
| **BEPA** | Intercept | 2 | -245.07 | 0.00 | 0.44 | 0.00 |
|  | EVI | 3 | -244.16 | 0.36 | 0.37 | 0.03 |
| **PIMA** | EVI + Biotic + Abiotic | 15 | -410.79 | 0.00 | 0.73 | 0.31 |
| **VAAN** | EVI + Biotic + Abiotic | 15 | -541.26 | 0.00 | 0.32 | 0.41 |
|  | EVI + Biotic | 12 | -544.89 | 0.04 | 0.31 | 0.39 |
|  | Land Cover + EVI + Biotic + Abiotic | 17 | -539.24 | 0.94 | 0.20 | 0.43 |
|  | Land Cover + EVI + Biotic | 14 | -543.20 | 1.44 | 0.15 | 0.40 |
| **Phytochemical: terpene (raw)** |  |  |  |  |  |  |
| **ABBA** | EVI | 3 | -269.49 | 0.00 | 0.52 | 0.05 |
| **PIMA** | EVI + Biotic + Abiotic | 15 | -471.87 | 0.00 | 0.47 | 0.27 |
|  | Biotic + Abiotic | 14 | -473.22 | 0.28 | 0.41 | 0.26 |

| Phytochemical: terpene (bm) |  |  |  |  |  |  |
| --- | --- | --- | --- | --- | --- | --- |
| **ABBA** | Biotic + Abiotic | 14 | -516.39 | 0.00 | 0.54 | 0.27 |
| **PIMA** | Intercept | 2 | -1126.03 | 0.00 | 0.40 | 0.00 |
|  | Abiotic | 5 | -1123.40 | 1.05 | 0.24 | 0.03 |
|  | EVI | 3 | -1125.99 | 2.00 | 0.15 | 0.00 |
| Phytochemical: monoterpene (raw) |  |  |  |  |  |  |
| **ABBA** | EVI + Abiotic | 6 | -229.81 | 0.00 | 0.44 | 0.12 |
|  | Abiotic | 5 | -231.81 | 1.75 | 0.18 | 0.08 |
|  | EVI | 3 | -234.08 | 1.93 | 0.17 | 0.04 |
| **PIMA** | Biotic + Abiotic | 14 | -290.67 | 0.00 | 0.67 | 0.24 |
| Phytochemical: monoterpene (bm) |  |  |  |  |  |  |
| **ABBA** | Biotic + Abiotic | 14 | -461.32 | 0.00 | 0.57 | 0.27 |
| **PIMA** | Intercept | 2 | -944.97 | 0.00 | 0.41 | 0.00 |
|  | Abiotic | 5 | -942.37 | 1.12 | 0.23 | 0.03 |
| Phytochemical: monoterpenic alcohol (raw) |  |  |  |  |  |  |
| **ABBA** | Land Cover | 4 | 42.49 | 0.00 | 0.26 | 0.05 |
|  | Intercept | 2 | 40.04 | 0.61 | 0.19 | 0.00 |
|  | Land Cover + EVI | 5 | 43.01 | 1.17 | 0.15 | 0.06 |
|  | Land Cover + Abiotic | 7 | 45.16 | 1.42 | 0.13 | 0.09 |
|  | Abiotic | 5 | 42.64 | 1.90 | 0.10 | 0.05 |
| **PIMA** | Biotic + Abiotic | 14 | 149.34 | 0.00 | 0.53 | 0.23 |
| Phytochemical: monoterpenic alcohol (bm) |  |  |  |  |  |  |
| **ABBA** | Intercept | 2 | -179.59 | 0.00 | 0.30 | 0.00 |
|  | EVI | 3 | -178.87 | 0.67 | 0.21 | 0.01 |
|  | Biotic + Abiotic | 14 | -165.76 | 0.94 | 0.19 | 0.23 |
| **PIMA** | Intercept | 2 | -487.60 | 0.00 | 0.42 | 0.00 |
|  | Abiotic | 5 | -485.08 | 1.27 | 0.22 | 0.03 |
|  | EVI | 3 | -487.49 | 1.86 | 0.16 | 0.00 |
| **Phytochemical: monoterpenic ester (raw)** |  |  |  |  |  |  |
| **ABBA** | Intercept | 2 | -115.95 | 0.00 | 0.21 | 0.00 |
|  | Land Cover | 4 | -113.88 | 0.13 | 0.19 | 0.04 |
|  | Land Cover + EVI | 5 | -112.80 | 0.19 | 0.19 | 0.06 |
|  | Abiotic | 5 | -113.31 | 1.22 | 0.11 | 0.05 |
|  | EVI | 3 | -115.73 | 1.67 | 0.09 | 0.00 |
| **PIMA** | Biotic + Abiotic | 14 | -310.53 | 0.00 | 0.57 | 0.27 |
|  | EVI* + Biotic + Abiotic | 15 | -309.93 | 1.23 | 0.31 | 0.27 |
| **Phytochemical: monoterpenic ester (bm)** |  |  |  |  |  |  |
| **ABBA** | Biotic + Abiotic | 14 | -332.19 | 0.00 | 0.47 | 0.27 |
| **PIMA** | Intercept | 2 | -926.76 | 0.00 | 0.42 | 0.00 |
|  | Abiotic | 5 | -924.24 | 1.26 | 0.22 | 0.03 |
| **Phytochemical: sesquiterpene (raw)** |  |  |  |  |  |  |
| **ABBA** | EVI + Abiotic | 6 | -61.73 | 0.00 | 0.33 | 0.10 |
|  | EVI | 3 | -65.31 | 0.54 | 0.25 | 0.04 |
|  | Abiotic | 5 | -63.51 | 1.31 | 0.17 | 0.07 |
| **PIMA** | Land Cover + EVI + Abiotic | 8 | 4.46 | 0.00 | 0.28 | 0.12 |
|  | Abiotic | 5 | 0.93 | 0.50 | 0.22 | 0.08 |
|  | EVI + Biotic + Abiotic | 15 | 11.98 | 1.29 | 0.15 | 0.19 |
|  | EVI + Abiotic | 6 | 1.53 | 1.45 | 0.13 | 0.08 |
| **Phytochemical: sesquiterpene (bm)** |  |  |  |  |  |  |
| **ABBA** | EVI | 3 | -284.44 | 0.00 | 0.23 | 0.02 |
|  | Biotic + Abiotic | 14 | -271.25 | 0.08 | 0.22 | 0.24 |
|  | Intercept | 2 | -285.69 | 0.36 | 0.19 | 0.00 |
| **PIMA** | Intercept | 2 | -566.61 | 0.00 | 0.38 | 0.00 |
|  | Abiotic | 5 | -563.89 | 0.86 | 0.25 | 0.03 |
|  | EVI | 3 | -566.55 | 1.95 | 0.15 | 0.00 |
| **Phytochemical: diversity** |  |  |  |  |  |  |
| **ABBA** | Abiotic | 5 | 113.25 | 0.00 | 0.31 | 0.06 |
|  | Intercept | 2 | 109.95 | 0.11 | 0.30 | 0.00 |
|  | EVI | 3 | 110.17 | 1.79 | 0.13 | 0.00 |
|  | EVI + Abiotic | 6 | 113.45 | 1.85 | 0.12 | 0.07 |
| **PIMA** | Land Cover | 4 | 189.24 | 0.00 | 0.31 | 0.04 |
|  | Biotic | 11 | 196.43 | 1.11 | 0.18 | 0.12 |
|  | EVI | 3 | 187.20 | 1.98 | 0.11 | 0.02 |

## Appendix 9

**Table A4.** Shows the coefficient sign (+/-) for all for top ranked models. Top models are presented in order of rank with Efron pseudo R^2^ presented in the last column. We use red coloured coefficients signs to indicate statistical significance at alpha =0.05. For land cover, Decid, and Mix indicate, deciduous, and mixed cover types respectively. EVI represents the Enhanced Vegetation Index. For biotic variables, AC indicates age class with 3, 4, 5 representing factor levels of 41-60, 61-80, and 81-100 years, respectively. HC indicates height class with 3, 4, 5 representing factor levels of 6.6-9.5, 9.6-12.5, 12.6-15.5 metres, respectively. CD indicates canopy density with 2, 3, 4 representing factor levels of 51-75, 26-50, 10-25 percent closed. For abiotic variables, E, A, and S represent elevation, aspect, and slope, respectively.

| Species | Land cover |  | Prod | Biotic factors |  |  |  |  |  |  |  |  | Abiotic factors |  |  | R<sup>2</sup> |
| --- | --- | --- | --- | --- | --- | --- | --- | --- | --- | --- | --- | --- | --- | --- | --- | --- |
|  | Decid | Mix | EVI | AC3 | AC4 | AC5 | HC3 | HC4 | HC5 | CD2 | CD3 | CD4 | E | A | S |  |
| Elemental: percent carbon |  |  |  |  |  |  |  |  |  |  |  |  |  |  |  |  |
| ABBA |  |  |  |  |  |  |  |  |  |  |  |  | + | − | + | 0.19 |
| ACRU |  |  |  |  |  |  |  |  |  |  |  |  | − | − | − | 0.09 |
| BEPA |  |  |  | + | + | + | − | − | − | − | − |  |  |  |  | 0.31 |
| PIMA |  |  |  | + | + | + | − | − | − | + | − | − | + | + | + | 0.64 |
| PIMA | − | − |  | + | + | + | − | − | − | + | − | − | + | + | + | 0.65 |
| VAAN | − | − |  | + | + | + | − | − | − | − | + | + | + | − | + | 0.37 |
| Elemental: percent nitrogen |  |  |  |  |  |  |  |  |  |  |  |  |  |  |  |  |
| ABBA | − | − | + |  |  |  |  |  |  |  |  |  | − | + | + | 0.29 |
| ACRU | − | − |  | − | − | − | + | + | + | − | − | − |  |  |  | 0.36 |
| PIMA |  |  | + | − | − | − | + | + | + | + | + | + |  |  |  | 0.36 |
| VAAN |  |  | + |  |  |  |  |  |  |  |  |  |  |  |  | 0.07 |
| VAAN | − | − | + |  |  |  |  |  |  |  |  |  |  |  |  | 0.09 |
| Elemental: percent phosphorus |  |  |  |  |  |  |  |  |  |  |  |  |  |  |  |  |
| ABBA |  |  |  |  |  |  |  |  |  |  |  |  | + | + | + | 0.1 |
| ACRU |  |  | + |  |  |  |  |  |  |  |  |  |  |  |  | 0.05 |
| ACRU |  |  | - |  |  |  |  |  |  |  |  |  | + | + | + | 0.1 |
| PIMA |  |  |  | - | - | - | + | + | + | + | + | - | + | + | + | 0.28 |
| VAAN |  |  | - | - | - | - | - | - | + | - | - | - |  |  |  | 0.47 |
| VAAN |  |  |  | - | - | - | - | - | - | + | - | - |  |  |  | 0.46 |
| Elemental: quantity carbon |  |  |  |  |  |  |  |  |  |  |  |  |  |  |  |  |
| ABBA |  |  |  | - | - | - | - | - | - | - | + | + | + | - | + | 0.35 |
| VAAN | + | + |  |  |  |  |  |  |  |  |  |  | + | + | - | 0.19 |
| Elemental: quantity nitrogen |  |  |  |  |  |  |  |  |  |  |  |  |  |  |  |  |
| ABBA |  |  |  | - | - | - | - | - | - | - | + | + | + | - | + | 0.35 |
| VAAN | + | + |  |  |  |  |  |  |  |  |  |  | + | + | - | 0.18 |
| Elemental: quantity phosphorus |  |  |  |  |  |  |  |  |  |  |  |  |  |  |  |  |
| ABBA |  |  |  | - | - | - | - | - | - | + | + | + | + | + | + | 0.32 |
| VAAN | + | + |  |  |  |  |  |  |  |  |  |  | + | + | - | 0.19 |
| Stoichiometric: C:N ratio |  |  |  |  |  |  |  |  |  |  |  |  |  |  |  |  |
| ABBA | + | + | - |  |  |  |  |  |  |  |  |  | + | - | - | 0.31 |
| ACRU | + | + |  | + | + | + | - | - | - | + | + | + |  |  |  | 0.39 |
| PIMA |  |  | - | + | + | + | - | - | - | - | - | - |  |  |  | 0.38 |
| VAAN |  |  | - |  |  |  |  |  |  |  |  |  |  |  |  | 0.09 |
| VAAN | + | + | - |  |  |  |  |  |  |  |  |  |  |  |  | 0.11 |
| Stoichiometric: C:P ratio |  |  |  |  |  |  |  |  |  |  |  |  |  |  |  |  |
| ABBA |  |  |  |  |  |  |  |  |  |  |  |  | + | - | - | 0.14 |
| *ACRU |  |  |  |  |  |  |  |  |  |  |  |  | - | - | - | 0.07 |
| PIMA |  |  |  | + | + | + | - | - | - | - | - | + | - | - | - | 0.3 |
| VAAN |  |  |  | + | + | + | + | + | - | - | + | + |  |  |  | 0.33 |
| VAAN |  |  | + | + | + | + | + | + | - | - | + | + |  |  |  | 0.34 |
| Stoichiometric: N:P ratio |  |  |  |  |  |  |  |  |  |  |  |  |  |  |  |  |
| *ABBA |  |  |  |  |  |  |  |  |  |  |  |  | - | - | - | 0.08 |
| *ABBA |  |  | + |  |  |  |  |  |  |  |  |  | - | - | - |  |
| ACRU |  |  | + |  |  |  |  |  |  |  |  |  | - | - | - | 0.17 |
| PIMA |  |  | + | + | + | + | - | + | - | + | - | + | - | - | - | 0.31 |
| VAAN |  |  | + | + | + | + | - | + | - | - | - | - | - | - | + | 0.41 |
| VAAN |  |  | + | + | + | + | - | + | - | - | - | + |  |  |  | 0.39 |
| VAAN | - | - | + | + | + | + | - | + | - | - | - | + | - | - | + | 0.43 |
| VAAN | - | - | + | + | + | + | - | + | - | - | - | + |  |  |  | 0.4 |
| Phytochemical: terpene (raw) |  |  |  |  |  |  |  |  |  |  |  |  |  |  |  |  |
| ABBA |  |  | + |  |  |  |  |  |  |  |  |  |  |  |  | 0.05 |
| PIMA |  |  | - | + | + | + | - | - | - | + | + | + | - | - | - | 0.27 |
| PIMA |  |  |  | + | + | + | - | - | - | + | + | + | - | - | - | 0.26 |
| Phytochemical: terpene (bm) |  |  |  |  |  |  |  |  |  |  |  |  |  |  |  |  |
| ABBA |  |  |  | - | - | - | - | - | - | + | + | + | + | - | + | 0.27 |
| Phytochemical: monoterpene (raw) |  |  |  |  |  |  |  |  |  |  |  |  |  |  |  |  |
| ABBA |  |  | + |  |  |  |  |  |  |  |  |  | + | - | + | 0.12 |
| ABBA |  |  |  |  |  |  |  |  |  |  |  |  | + | - | + | 0.08 |
| ABBA |  |  | + |  |  |  |  |  |  |  |  |  |  |  |  | 0.04 |
| PIMA |  |  |  | + | + | + | - | - | - | + | + | + | - | - | - | 0.24 |
| Phytochemical: monoterpene (bm) |  |  |  |  |  |  |  |  |  |  |  |  |  |  |  |  |
| ABBA |  |  |  | - | - | - | - | - | - | + | + | + | + | - | + | 0.27 |
| Phytochemical: monoterpene alcohol (raw) |  |  |  |  |  |  |  |  |  |  |  |  |  |  |  |  |
| *ABBA | + | + |  |  |  |  |  |  |  |  |  |  |  |  |  | 0.05 |
| PIMA |  |  |  | + | + | + | - | - | - | + | + | + | - | - | - | 0.23 |
| Phytochemical: monoterpene ester (raw) |  |  |  |  |  |  |  |  |  |  |  |  |  |  |  |  |
| PIMA |  |  |  | + | + | + | - | - | - | + | + | + | - | - | - | 0.27 |
| Phytochemical: monoterpene ester (bm) |  |  |  |  |  |  |  |  |  |  |  |  |  |  |  |  |
| ABBA |  |  |  | - | - | - | - | - | - | + | + | + | + | - | + | 0.27 |
| Phytochemical: sesquiterpene (raw) |  |  |  |  |  |  |  |  |  |  |  |  |  |  |  |  |
| ABBA |  |  | + |  |  |  |  |  |  |  |  |  | - | - | - | 0.1 |
| ABBA |  |  | + |  |  |  |  |  |  |  |  |  |  |  |  | 0.04 |
| ABBA |  |  |  |  |  |  |  |  |  |  |  |  | - | - | - | 0.07 |
| PIMA | - | + | - |  |  |  |  |  |  |  |  |  | - | - | - | 0.12 |
| PIMA |  |  |  |  |  |  |  |  |  |  |  |  | - | - | - | 0.08 |
| PIMA |  |  | - | + | - | + | - | - | - | - | - | + | - | + | - | 0.19 |
| PIMA |  |  | - |  |  |  |  |  |  |  |  |  | - | - | - | 0.08 |
| Phytochemical: sesquiterpene (bm) |  |  |  |  |  |  |  |  |  |  |  |  |  |  |  |  |
| *ABBA |  |  | + |  |  |  |  |  |  |  |  |  |  |  |  | 0.02 |
| *ABBA |  |  |  | - | - | - | - | - | - | + | + | + | + | - | + | 0.24 |
| Phytochemical: diversity |  |  |  |  |  |  |  |  |  |  |  |  |  |  |  |  |
| *ABBA |  |  |  |  |  |  |  |  |  |  |  |  | - | - | + | 0.06 |
| PIMA | + | + |  |  |  |  |  |  |  |  |  |  |  |  |  | 0.04 |
| PIMA |  |  |  | - | - | - | - | + | + | - | - | - |  |  |  | 0.12 |
| PIMA |  |  | + |  |  |  |  |  |  |  |  |  |  |  |  | 0.02 |

## Appendix 10

**Figure A4.**
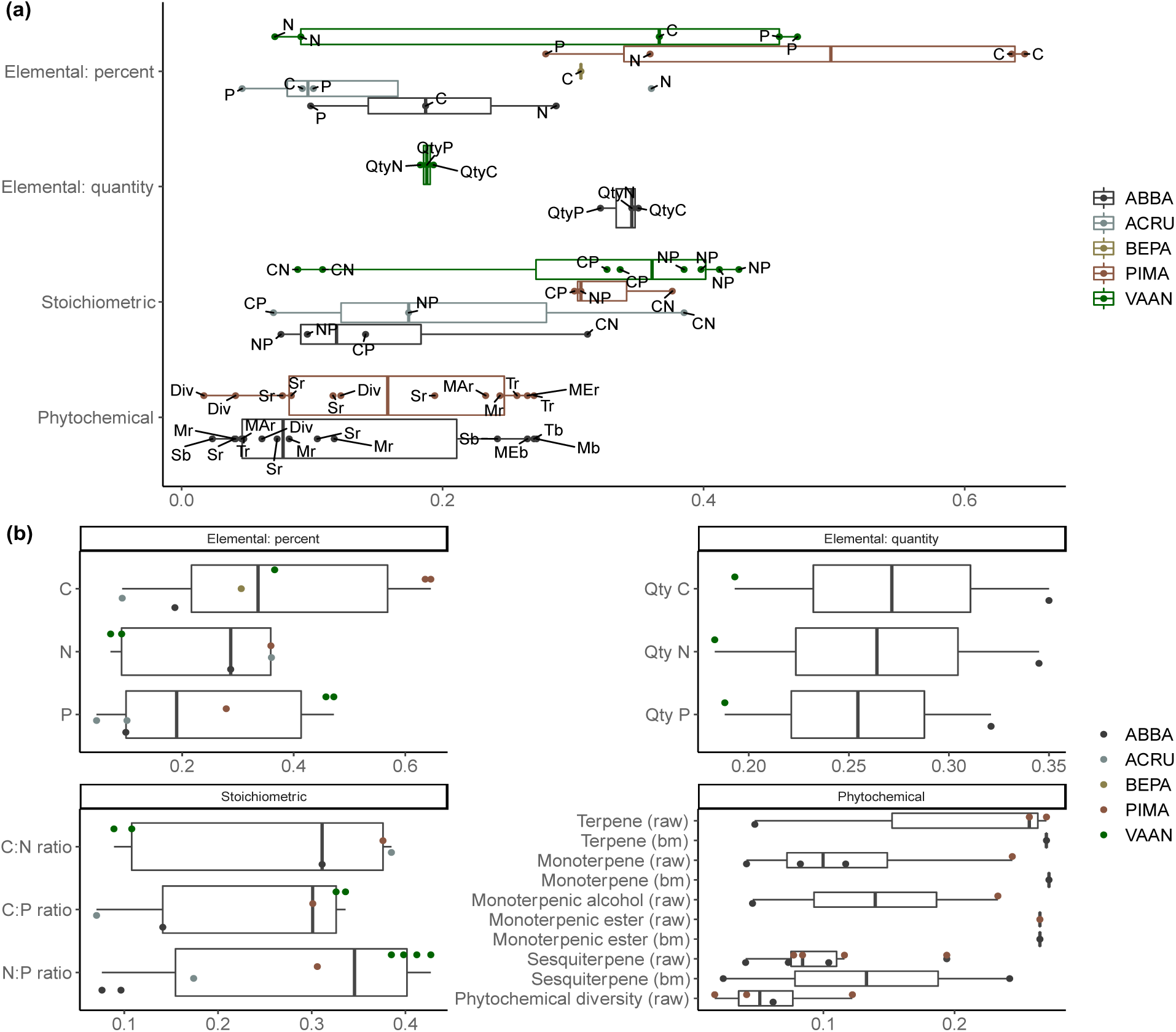
Distribution of pseudo R^2^ values across species, at the trait type level (a) and at the trait level (b), for all top ranked models. At trait type level, we show pseudo R^2^ values for element percent and quantity, stoichiometric, and phytochemical traits. At the trait level we show individual traits of percent elemental (i.e., %C, %N, and %P), quantity elemental (i.e., C, N, and P on a g/m^2^ biomass basis), stoichiometric ratios (i.e., C:N, C:P, and N:P), and phytochemical groups (terpene, monoterpene, monoterpenic alcohol, monoterpenic ester, sesquiterpene, and phytochemical diversity) on a raw of biomass basis, indicated as either (raw) or (bm) suffixes, respectively. Species bar and point colours are the same between plots. In addition, labels are provided in (a) to identify individual traits for a given species within a trait type.

## Appendix 11

**Figure A5.**
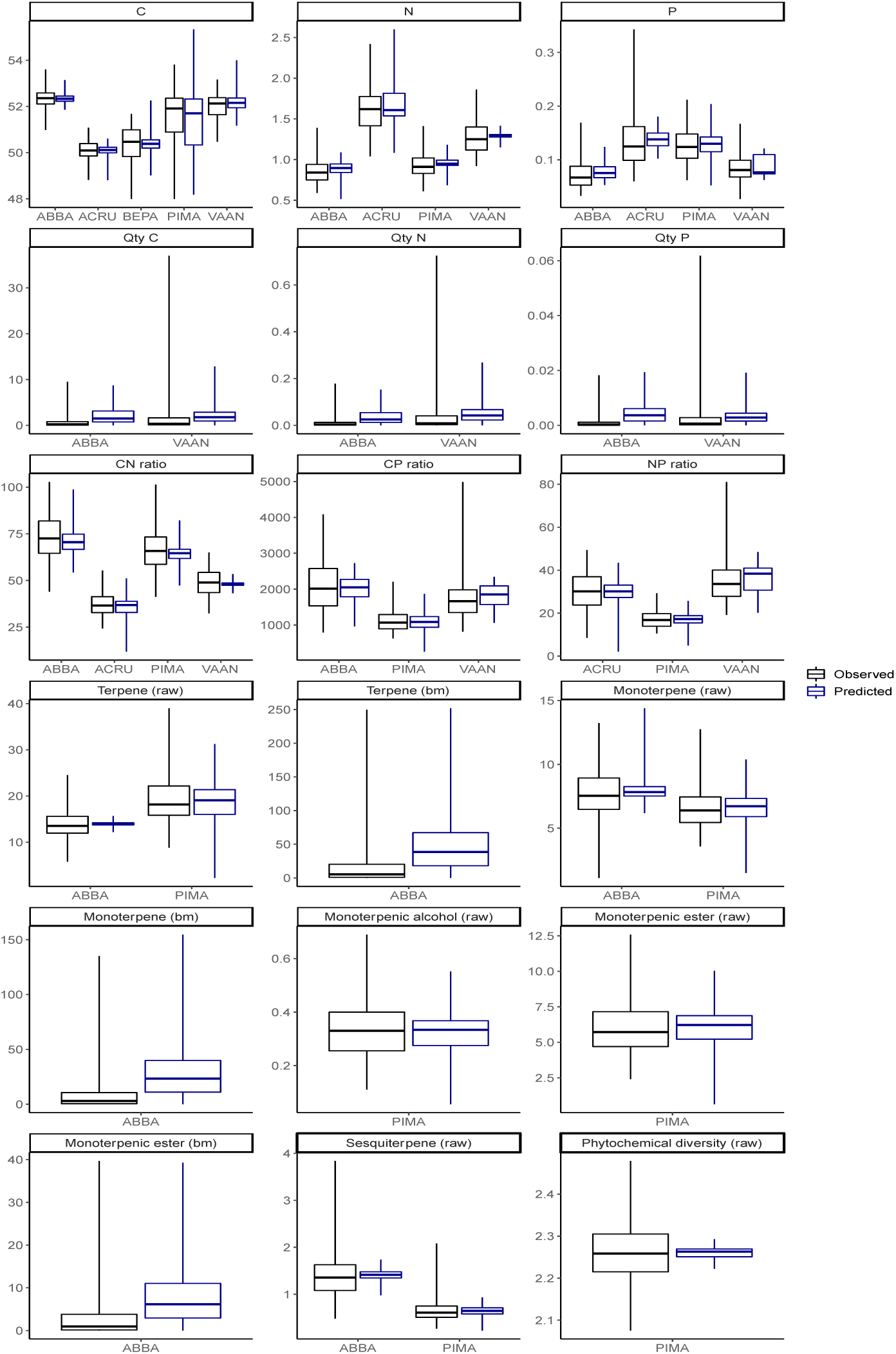
Comparison of observed (data) and predicted (raster values) data for foliar elemental, stoichiometric, and phytochemical traits for each of our study species where a plausible explanation was determined. Generally, medians are consistent between observed and predicted data, however, ranges differ with predicted often having a larger variance.

## Appendix 12

**Table A5.**
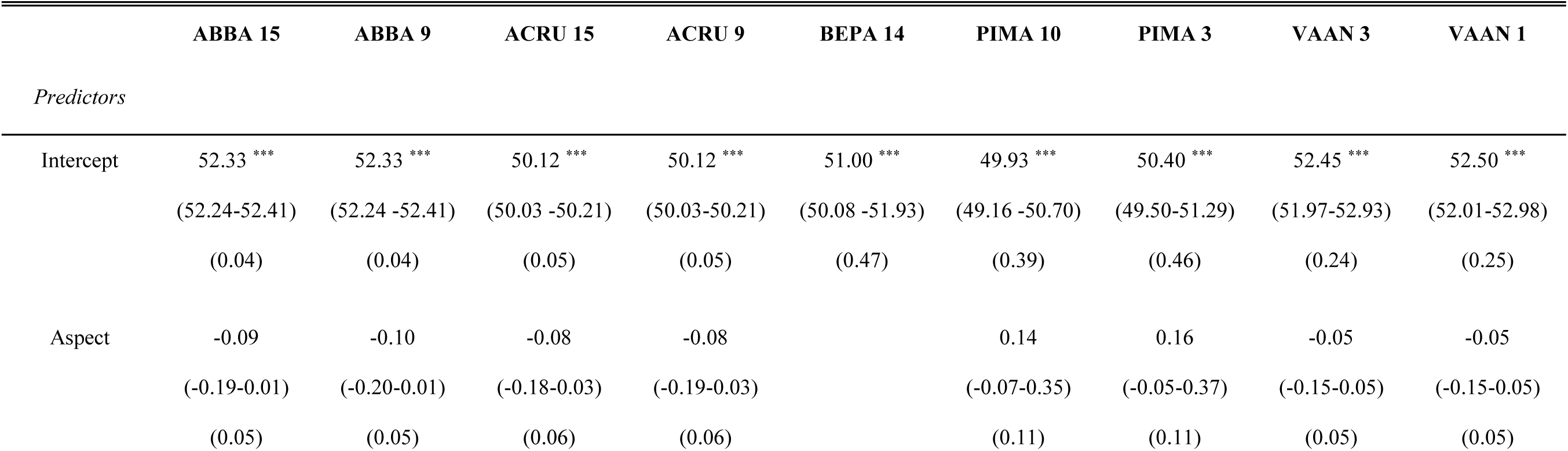
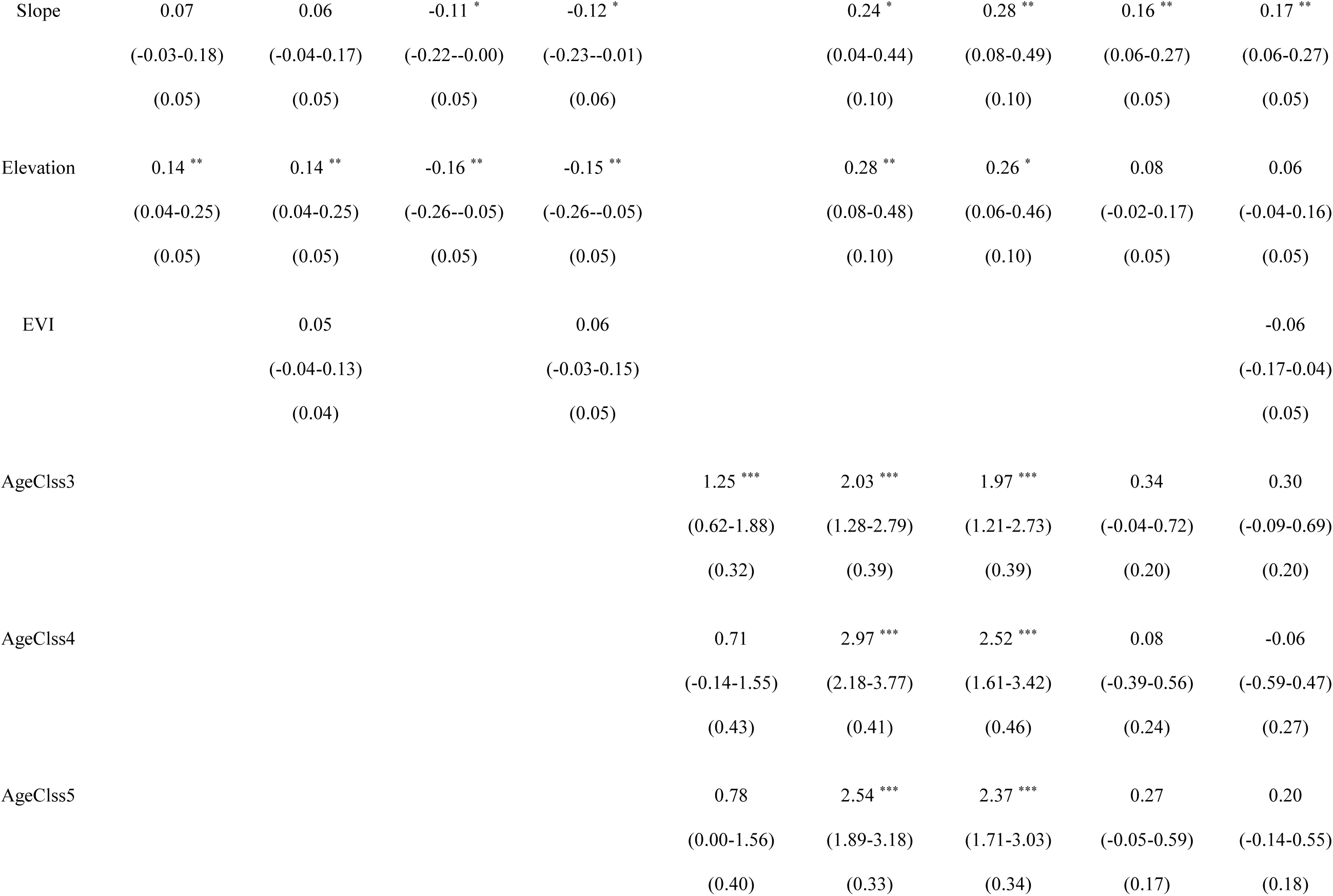
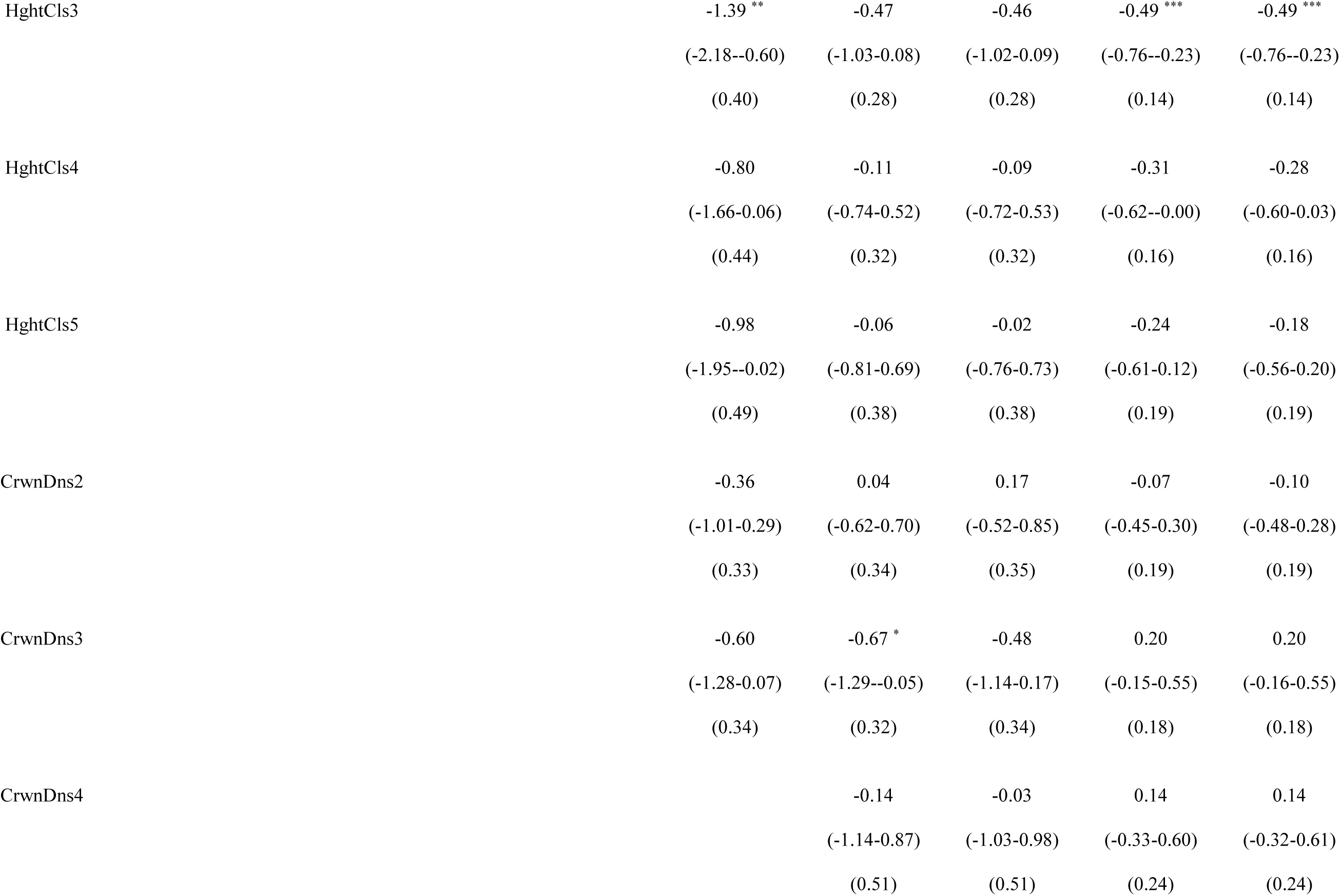
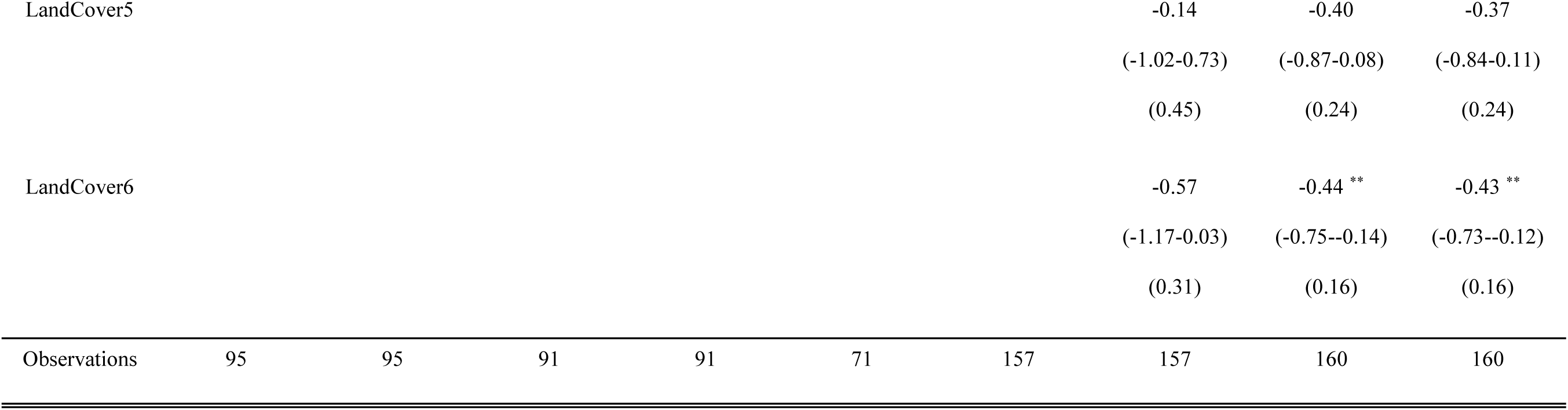
Foliar percent carbon trait coefficient estimates, confidence intervals, and standard error values for top ranked models (< 2 ΔAIC_c_). Species codes are used for balsam fir (ABBA), red maple (ACRU), white birch (BEPA), black spruce (PIMA, and lowbush blueberry (VAAN). If there is more than one top ranked model per species, we present in order of ΔAIC_c_ rank. Model numbers are supplied beside the species code in the top row (see Table 1 for model descriptions). Predictors include land cover (LandCover5 and LandCover6 represent deciduous and mix wood conditions), EVI (i.e., proxy for productivity), abiotic factors (aspect, slope, elevation), and biotic factors: AgeClss3 (41-60 years old), AgeClss 4 (61-80 years old), AgeClss5 (81-100 years old), HghtCls3 (6.6 −9.5 m), HghtCls4 (9.6-12.5 m), HghtCls5 (12.6-15.5m), CrwnDns2 (51-75 % closed), CrwnDns3 (26-50%), CrwnDn4 (10-25 % closed). Total number of observations are provided in the bottom row. In addition, asterisks are used to indicate coefficient significance as follows: *\* p<0.05 ** p<0.01 *** p<0.001*.

|  | ABBA 15 | ABBA 9 | ACRU 15 | ACRU 9 | BEPA 14 | PIMA 10 | PIMA 3 | VAAN 3 | VAAN 1 |
| --- | --- | --- | --- | --- | --- | --- | --- | --- | --- |
| *Predictors* |  |  |  |  |  |  |  |  |  |
| Intercept | 52.33 ***<br>(52.24-52.41)<br>(0.04) | 52.33 ***<br>(52.24 -52.41)<br>(0.04) | 50.12 ***<br>(50.03 -50.21)<br>(0.05) | 50.12 ***<br>(50.03-50.21)<br>(0.05) | 51.00 ***<br>(50.08 -51.93)<br>(0.47) | 49.93 ***<br>(49.16 -50.70)<br>(0.39) | 50.40 ***<br>(49.50-51.29)<br>(0.46) | 52.45 ***<br>(51.97-52.93)<br>(0.24) | 52.50 ***<br>(52.01-52.98)<br>(0.25) |
| Aspect | -0.09<br>(-0.19-0.01)<br>(0.05) | -0.10<br>(-0.20-0.01)<br>(0.05) | -0.08<br>(-0.18-0.03)<br>(0.06) | -0.08<br>(-0.19-0.03)<br>(0.06) |  | 0.14<br>(-0.07-0.35)<br>(0.11) | 0.16<br>(-0.05-0.37)<br>(0.11) | -0.05<br>(-0.15-0.05)<br>(0.05) | -0.05<br>(-0.15-0.05)<br>(0.05) |
| Slope | 0.07<br>(-0.03-0.18)<br>(0.05) | 0.06<br>(-0.04-0.17)<br>(0.05) | -0.11 *<br>(-0.22--0.00)<br>(0.05) | -0.12 *<br>(-0.23--0.01)<br>(0.06) |  | 0.24 *<br>(0.04-0.44)<br>(0.10) | 0.28 **<br>(0.08-0.49)<br>(0.10) | 0.16 **<br>(0.06-0.27)<br>(0.05) | 0.17 **<br>(0.06-0.27)<br>(0.05) |
| Elevation | 0.14 **<br>(0.04-0.25)<br>(0.05) | 0.14 **<br>(0.04-0.25)<br>(0.05) | -0.16 **<br>(-0.26--0.05)<br>(0.05) | -0.15 **<br>(-0.26--0.05)<br>(0.05) |  | 0.28 **<br>(0.08-0.48)<br>(0.10) | 0.26 *<br>(0.06-0.46)<br>(0.10) | 0.08<br>(-0.02-0.17)<br>(0.05) | 0.06<br>(-0.04-0.16)<br>(0.05) |
| EVI |  | 0.05<br>(-0.04-0.13)<br>(0.04) |  | 0.06<br>(-0.03-0.15)<br>(0.05) |  |  |  |  | -0.06<br>(-0.17-0.04)<br>(0.05) |
| AgeClss3 |  |  |  |  | 1.25 ***<br>(0.62-1.88)<br>(0.32) | 2.03 ***<br>(1.28-2.79)<br>(0.39) | 1.97 ***<br>(1.21-2.73)<br>(0.39) | 0.34<br>(-0.04-0.72)<br>(0.20) | 0.30<br>(-0.09-0.69)<br>(0.20) |
| AgeClss4 |  |  |  |  | 0.71<br>(-0.14-1.55)<br>(0.43) | 2.97 ***<br>(2.18-3.77)<br>(0.41) | 2.52 ***<br>(1.61-3.42)<br>(0.46) | 0.08<br>(-0.39-0.56)<br>(0.24) | -0.06<br>(-0.59-0.47)<br>(0.27) |
| AgeClss5 |  |  |  |  | 0.78<br>(0.00-1.56)<br>(0.40) | 2.54 ***<br>(1.89-3.18)<br>(0.33) | 2.37 ***<br>(1.71-3.03)<br>(0.34) | 0.27<br>(-0.05-0.59)<br>(0.17) | 0.20<br>(-0.14-0.55)<br>(0.18) |

|  |  |  |  |  |  |
| --- | --- | --- | --- | --- | --- |
| HghtCls3 | -1.39 ** | -0.47 | -0.46 | -0.49 *** | -0.49 *** |
|  | (-2.18--0.60) | (-1.03-0.08) | (-1.02-0.09) | (-0.76--0.23) | (-0.76--0.23) |
|  | (0.40) | (0.28) | (0.28) | (0.14) | (0.14) |
| HghtCls4 | -0.80 | -0.11 | -0.09 | -0.31 | -0.28 |
|  | (-1.66-0.06) | (-0.74-0.52) | (-0.72-0.53) | (-0.62--0.00) | (-0.60-0.03) |
|  | (0.44) | (0.32) | (0.32) | (0.16) | (0.16) |
| HghtCls5 | -0.98 | -0.06 | -0.02 | -0.24 | -0.18 |
|  | (-1.95--0.02) | (-0.81-0.69) | (-0.76-0.73) | (-0.61-0.12) | (-0.56-0.20) |
|  | (0.49) | (0.38) | (0.38) | (0.19) | (0.19) |
| CrwnDns2 | -0.36 | 0.04 | 0.17 | -0.07 | -0.10 |
|  | (-1.01-0.29) | (-0.62-0.70) | (-0.52-0.85) | (-0.45-0.30) | (-0.48-0.28) |
|  | (0.33) | (0.34) | (0.35) | (0.19) | (0.19) |
| CrwnDns3 | -0.60 | -0.67 * | -0.48 | 0.20 | 0.20 |
|  | (-1.28-0.07) | (-1.29--0.05) | (-1.14-0.17) | (-0.15-0.55) | (-0.16-0.55) |
|  | (0.34) | (0.32) | (0.34) | (0.18) | (0.18) |
| CrwnDns4 |  | -0.14 | -0.03 | 0.14 | 0.14 |
|  |  | (-1.14-0.87) | (-1.03-0.98) | (-0.33-0.60) | (-0.32-0.61) |
|  |  | (0.51) | (0.51) | (0.24) | (0.24) |
| LandCover5 |  |  |  |  |  |  | -0.14 | -0.40 | -0.37 |
|  |  |  |  |  |  |  | (-1.02-0.73) | (-0.87-0.08) | (-0.84-0.11) |
|  |  |  |  |  |  |  | (0.45) | (0.24) | (0.24) |
| LandCover6 |  |  |  |  |  |  | -0.57 | -0.44 ** | -0.43 ** |
|  |  |  |  |  |  |  | (-1.17-0.03) | (-0.75--0.14) | (-0.73--0.12) |
|  |  |  |  |  |  |  | (0.31) | (0.16) | (0.16) |
| Observations | 95 | 95 | 91 | 91 | 71 | 157 | 157 | 160 | 160 |

## Appendix 13

**Table A6.**
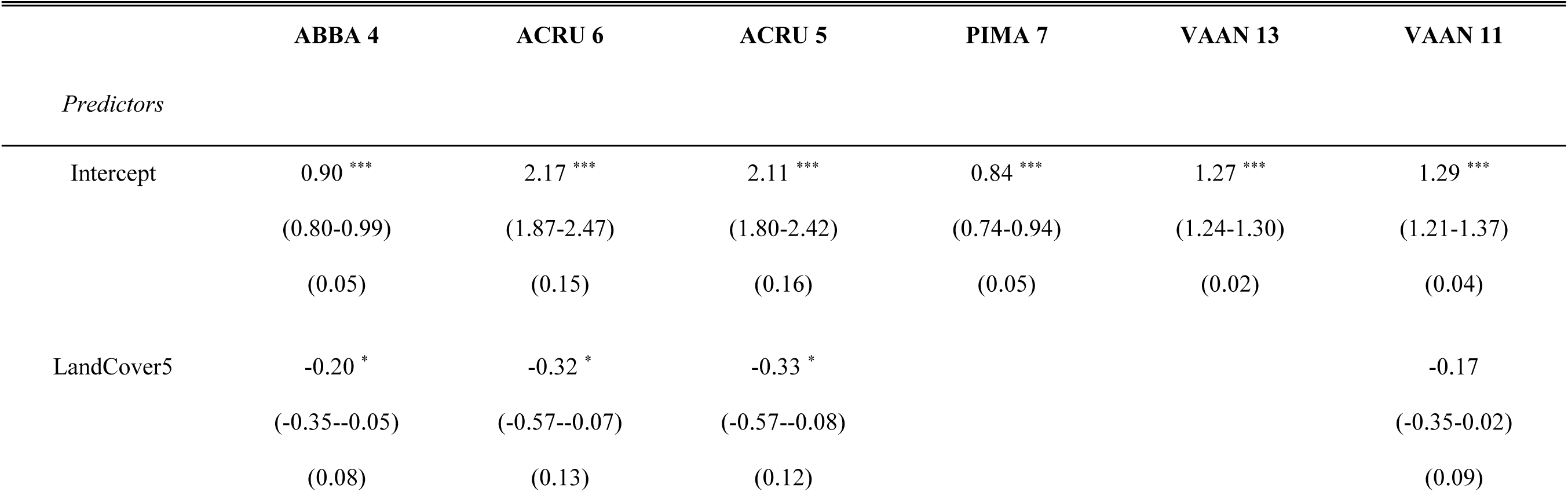
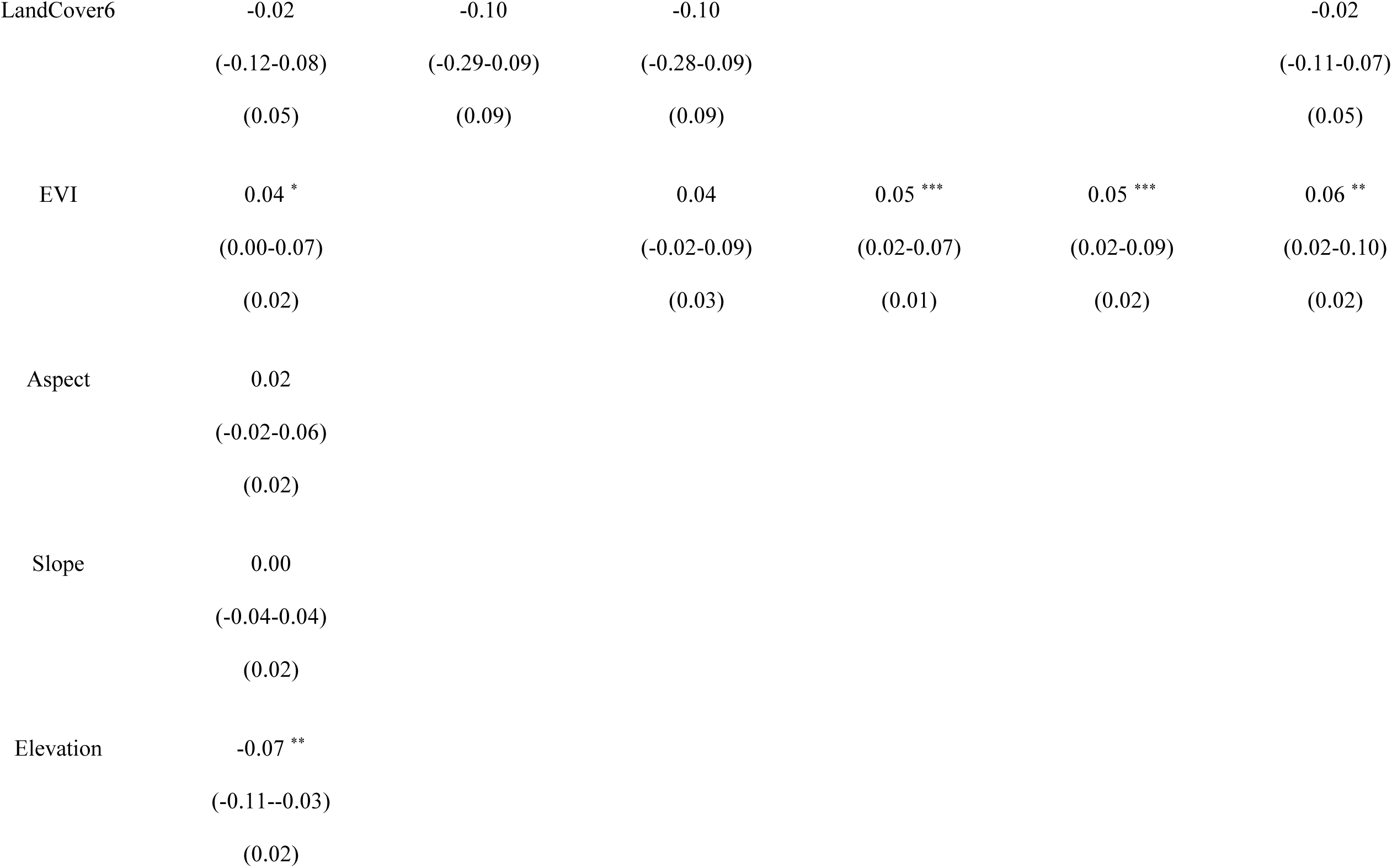
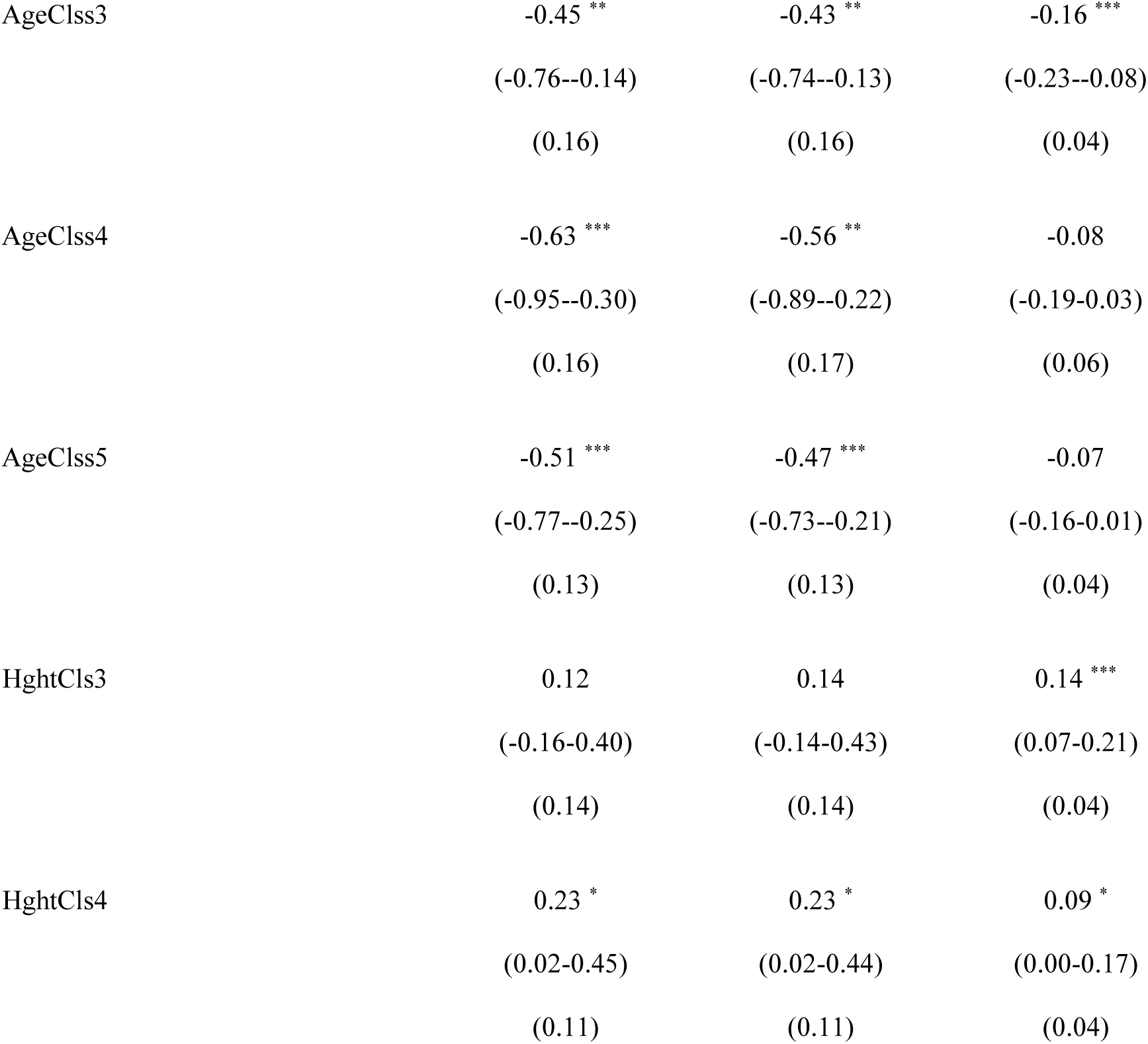
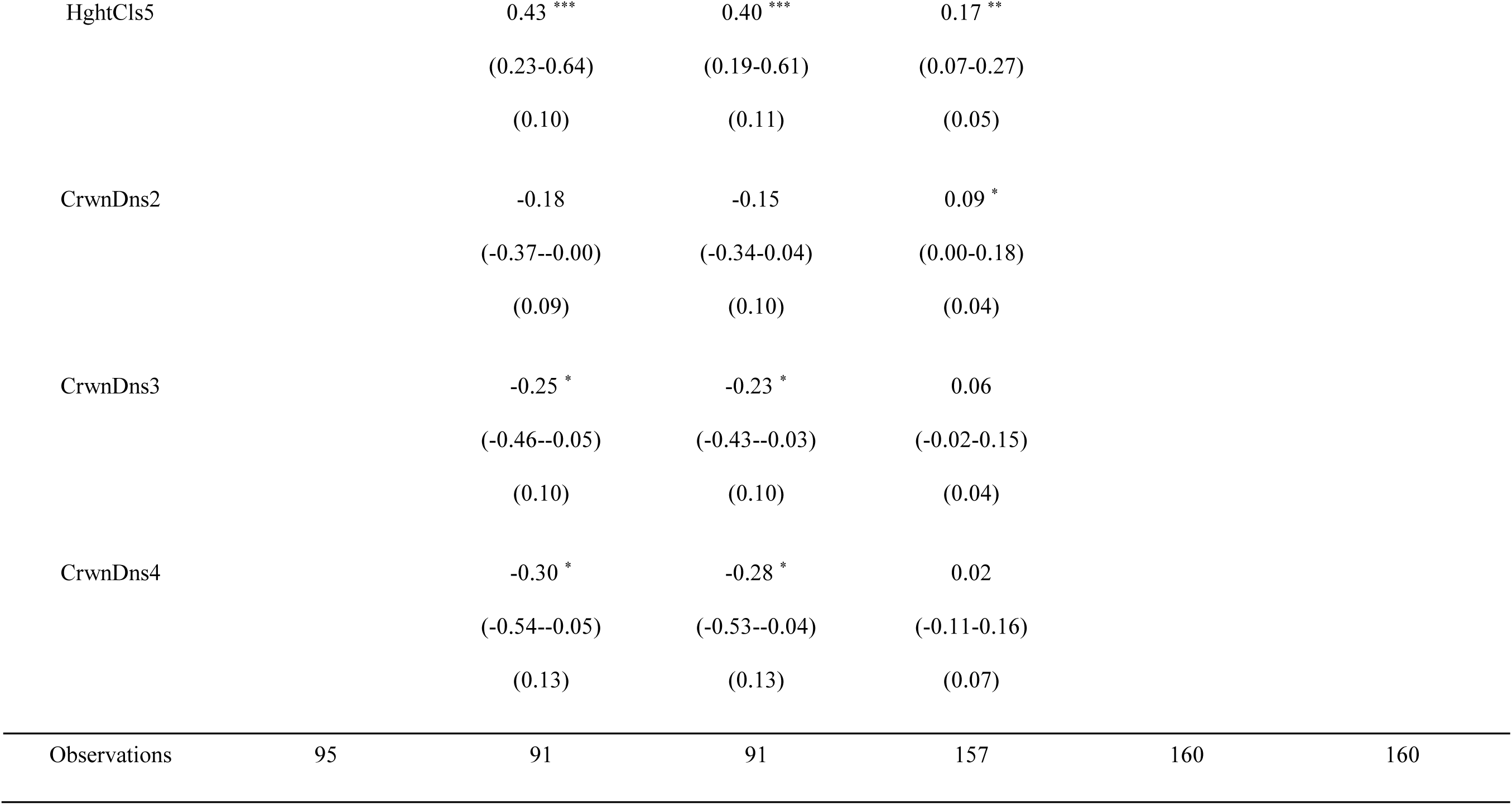
Foliar percent nitrogen trait coefficient estimates, confidence intervals, and standard error values for top ranked models (< 2 ΔAIC_c_). Species codes are used for balsam fir (ABBA), red maple (ACRU), white birch (BEPA), black spruce (PIMA, and lowbush blueberry (VAAN). If there is more than one top ranked model per species, we present in order of ΔAIC_c_ rank. Model numbers are supplied beside the species code in the top row (see Table 1 for model descriptions). Predictors include land cover (LandCover5 and LandCover6 represent deciduous and mix wood conditions), EVI (i.e., proxy for productivity), abiotic factors (aspect, slope, elevation), and biotic factors: AgeClss3 (41-60 years old), AgeClss 4 (61-80 years old), AgeClss5 (81-100 years old), HghtCls3 (6.6-9.5 m), HghtCls4 (9.6-12.5 m), HghtCls5 (12.6-15.5m), CrwnDns2 (51-75 % closed), CrwnDns3 (26-50%), CrwnDn4 (10-25 % closed). Total number of observations are provided in the bottom row. In addition, asterisks are used to indicate coefficient significance as follows: *\* p<0.05 ** p<0.01 *** p<0.001*.

|  | ABBA 4 | ACRU 6 | ACRU 5 | PIMA 7 | VAAN 13 | VAAN 11 |
| --- | --- | --- | --- | --- | --- | --- |
| *Predictors* |  |  |  |  |  |  |
| Intercept | 0.90 ***<br>(0.80-0.99)<br>(0.05) | 2.17 ***<br>(1.87-2.47)<br>(0.15) | 2.11 ***<br>(1.80-2.42)<br>(0.16) | 0.84 ***<br>(0.74-0.94)<br>(0.05) | 1.27 ***<br>(1.24-1.30)<br>(0.02) | 1.29 ***<br>(1.21-1.37)<br>(0.04) |
| LandCover5 | -0.20 *<br>(-0.35--0.05)<br>(0.08) | -0.32 *<br>(-0.57--0.07)<br>(0.13) | -0.33 *<br>(-0.57--0.08)<br>(0.12) |  |  | -0.17<br>(-0.35-0.02)<br>(0.09) |
| LandCover6 | -0.02<br>(-0.12-0.08)<br>(0.05) | -0.10<br>(-0.29-0.09)<br>(0.09) | -0.10<br>(-0.28-0.09)<br>(0.09) |  |  | -0.02<br>(-0.11-0.07)<br>(0.05) |
| EVI | 0.04 *<br>(0.00-0.07)<br>(0.02) |  | 0.04<br>(-0.02-0.09)<br>(0.03) | 0.05 ***<br>(0.02-0.07)<br>(0.01) | 0.05 ***<br>(0.02-0.09)<br>(0.02) | 0.06 **<br>(0.02-0.10)<br>(0.02) |
| Aspect | 0.02<br>(-0.02-0.06)<br>(0.02) |  |  |  |  |  |
| Slope | 0.00<br>(-0.04-0.04)<br>(0.02) |  |  |  |  |  |
| Elevation | -0.07 **<br>(-0.11--0.03)<br>(0.02) |  |  |  |  |  |

|  |  |  |  |
| --- | --- | --- | --- |
| AgeCls3 | -0.45 **<br>(-0.76--0.14)<br>(0.16) | -0.43 **<br>(-0.74--0.13)<br>(0.16) | -0.16 ***<br>(-0.23--0.08)<br>(0.04) |
| AgeCls4 | -0.63 ***<br>(-0.95--0.30)<br>(0.16) | -0.56 **<br>(-0.89--0.22)<br>(0.17) | -0.08<br>(-0.19-0.03)<br>(0.06) |
| AgeCls5 | -0.51 ***<br>(-0.77--0.25)<br>(0.13) | -0.47 ***<br>(-0.73--0.21)<br>(0.13) | -0.07<br>(-0.16-0.01)<br>(0.04) |
| HghtCls3 | 0.12<br>(-0.16-0.40)<br>(0.14) | 0.14<br>(-0.14-0.43)<br>(0.14) | 0.14 ***<br>(0.07-0.21)<br>(0.04) |
| HghtCls4 | 0.23 *<br>(0.02-0.45)<br>(0.11) | 0.23 *<br>(0.02-0.44)<br>(0.11) | 0.09 *<br>(0.00-0.17)<br>(0.04) |
| HghtCls5 | 0.43 *** | 0.40 *** | 0.17 ** |
|  | (0.23-0.64) | (0.19-0.61) | (0.07-0.27) |
|  | (0.10) | (0.11) | (0.05) |
| CrwnDns2 | -0.18 | -0.15 | 0.09 * |
|  | (-0.37--0.00) | (-0.34-0.04) | (0.00-0.18) |
|  | (0.09) | (0.10) | (0.04) |
| CrwnDns3 | -0.25 * | -0.23 * | 0.06 |
|  | (-0.46--0.05) | (-0.43--0.03) | (-0.02-0.15) |
|  | (0.10) | (0.10) | (0.04) |
| CrwnDns4 | -0.30 * | -0.28 * | 0.02 |
|  | (-0.54--0.05) | (-0.53--0.04) | (-0.11-0.16) |
|  | (0.13) | (0.13) | (0.07) |
| Observations | 95 | 91 | 91 | 157 | 160 | 160 |

## Appendix 14

**Table A7.**
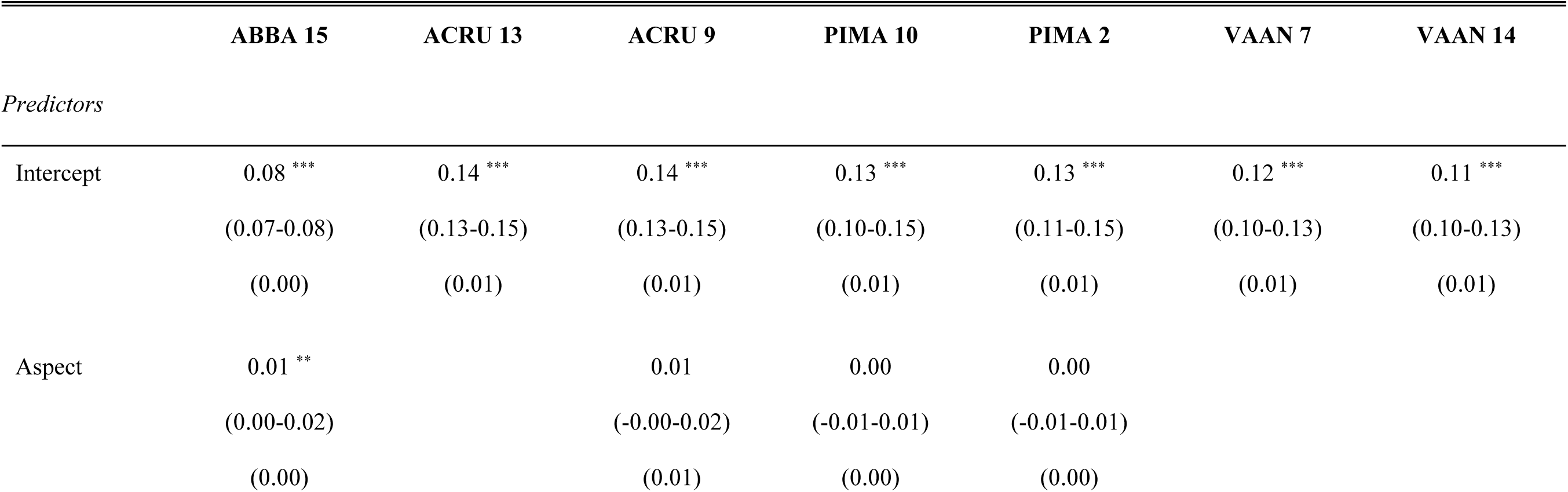
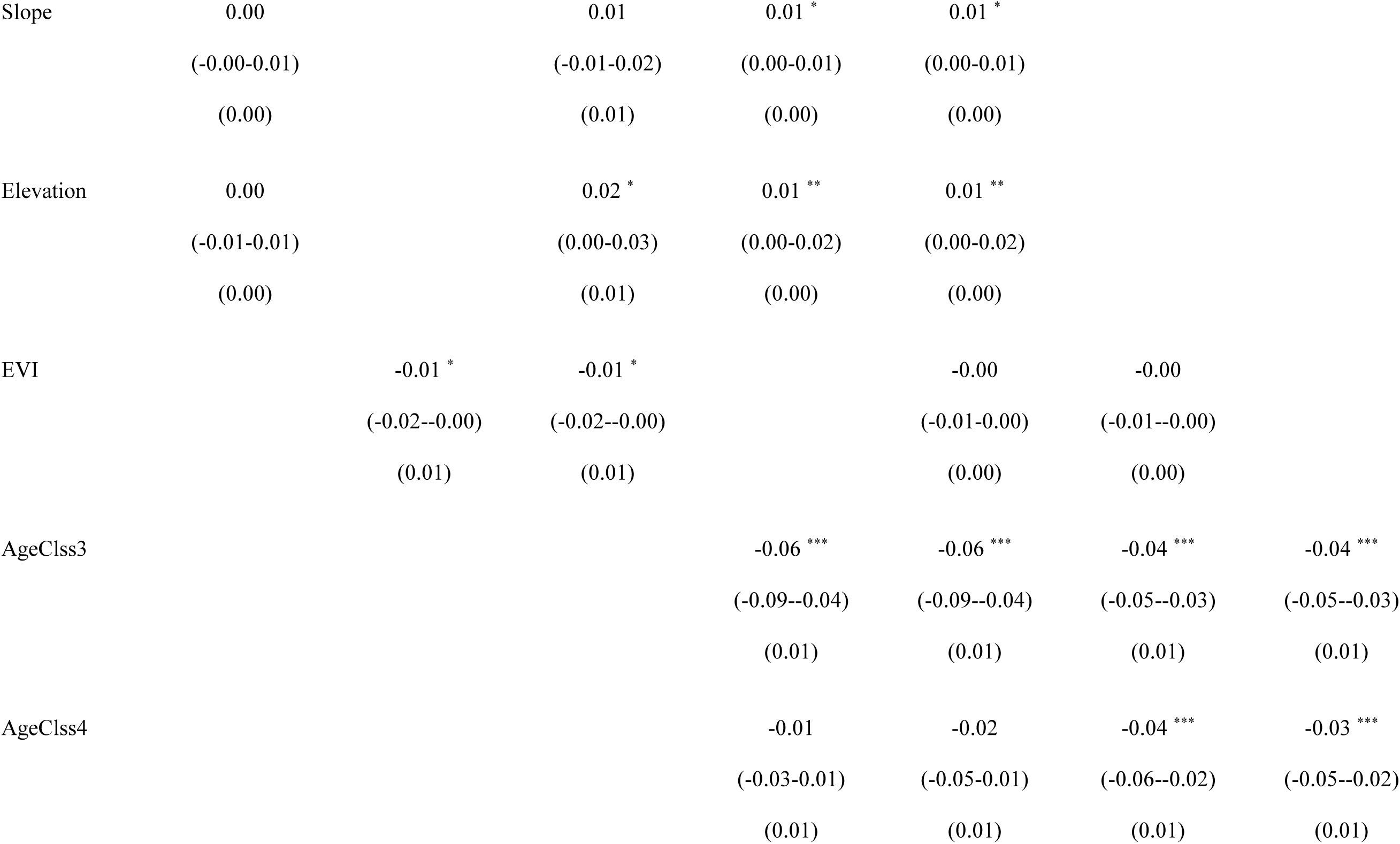
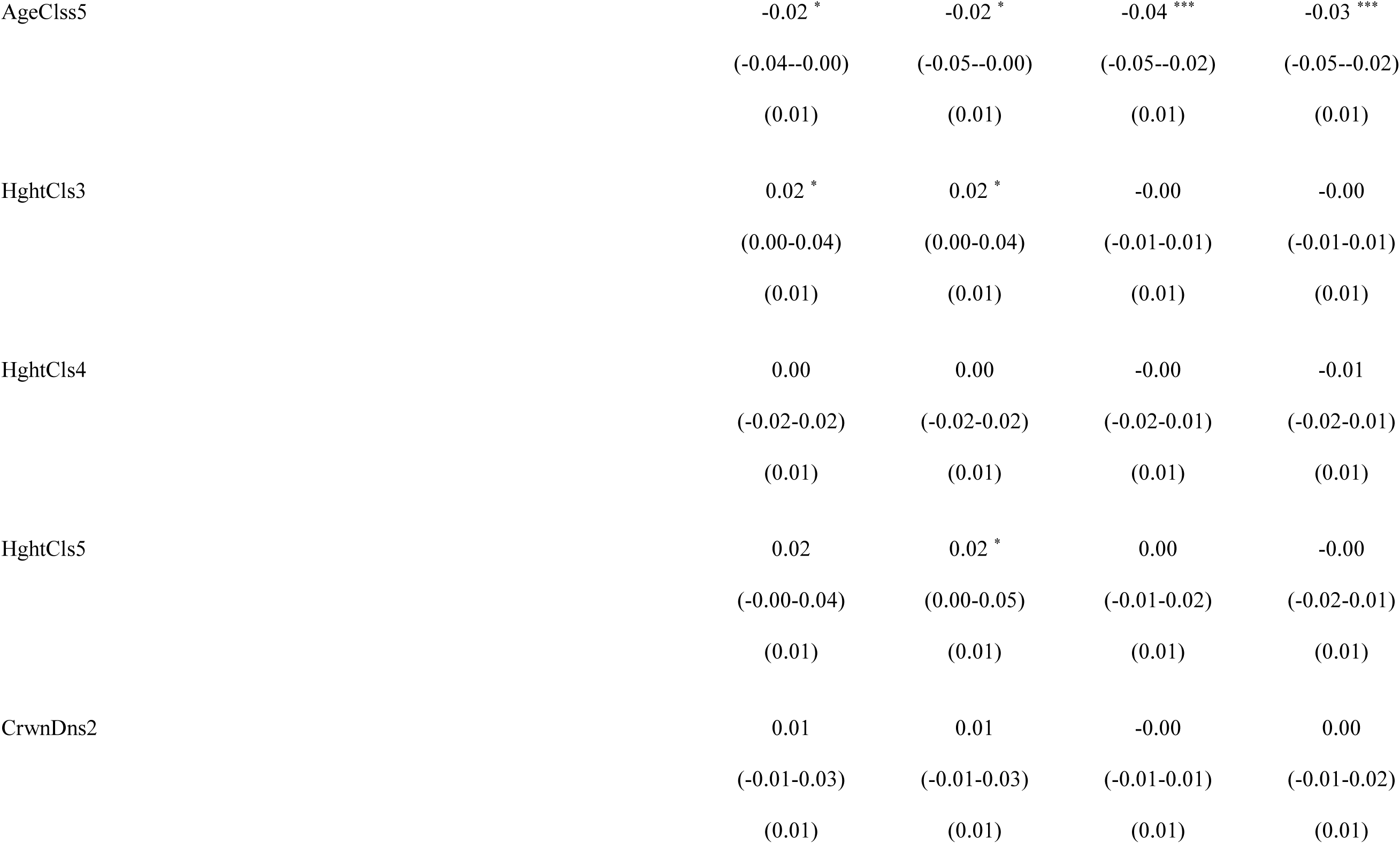
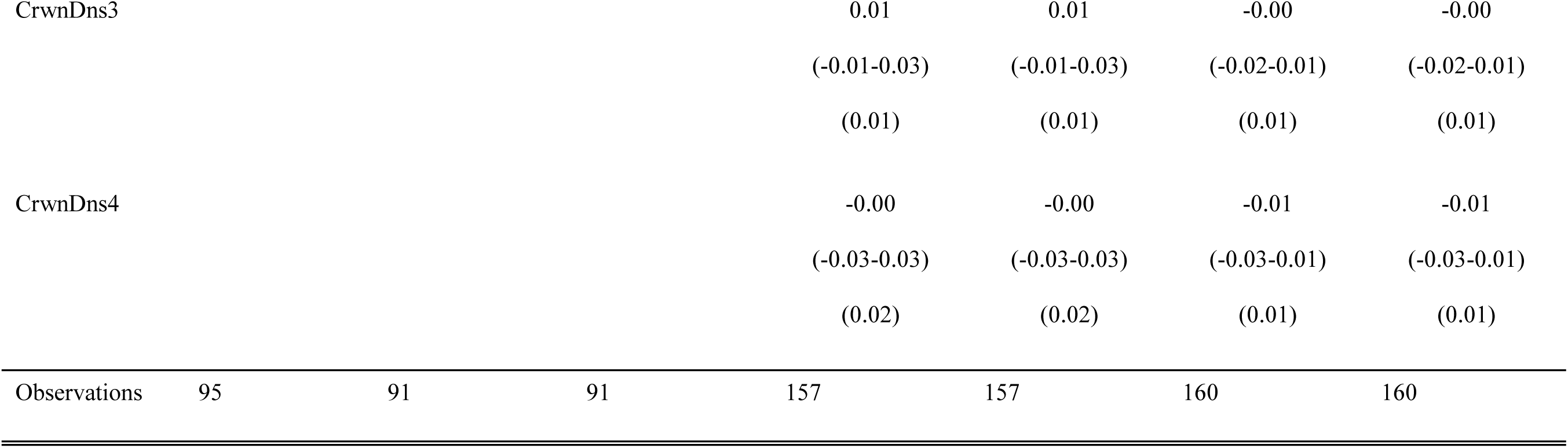
Foliar percent phosphorus trait coefficient estimates, confidence intervals, and standard error values for top ranked models (< 2 ΔAIC_c_). Species codes are used for balsam fir (ABBA), red maple (ACRU), white birch (BEPA), black spruce (PIMA, and lowbush blueberry (VAAN). If there is more than one top ranked model per species, we present in order of ΔAIC_c_ rank. Model numbers are supplied beside the species code in the top row (see Table 1 for model descriptions). Predictors include land cover (LandCover5 and LandCover6 represent deciduous and mix wood conditions), EVI (i.e., proxy for productivity), abiotic factors (aspect, slope, elevation), and biotic factors: AgeClss3 (41-60 years old), AgeClss 4 (61-80 years old), AgeClss5 (81-100 years old), HghtCls3 (6.6-9.5 m), HghtCls4 (9.6-12.5 m), HghtCls5 (12.6-15.5m), CrwnDns2 (51-75 % closed), CrwnDns3 (26-50%), CrwnDn4 (10-25 % closed). Total number of observations are provided in the bottom row. In addition, asterisks are used to indicate coefficient significance as follows: *\* p<0.05 ** p<0.01 *** p<0.001*.

|  | ABBA 15 | ACRU 13 | ACRU 9 | PIMA 10 | PIMA 2 | VAAN 7 | VAAN 14 |
| --- | --- | --- | --- | --- | --- | --- | --- |
| *Predictors* |  |  |  |  |  |  |  |
| Intercept | 0.08 ***<br>(0.07-0.08)<br>(0.00) | 0.14 ***<br>(0.13-0.15)<br>(0.01) | 0.14 ***<br>(0.13-0.15)<br>(0.01) | 0.13 ***<br>(0.10-0.15)<br>(0.01) | 0.13 ***<br>(0.11-0.15)<br>(0.01) | 0.12 ***<br>(0.10-0.13)<br>(0.01) | 0.11 ***<br>(0.10-0.13)<br>(0.01) |
| Aspect | 0.01 **<br>(0.00-0.02)<br>(0.00) |  | 0.01<br>(-0.00-0.02)<br>(0.01) | 0.00<br>(-0.01-0.01)<br>(0.00) | 0.00<br>(-0.01-0.01)<br>(0.00) |  |  |
| Slope | 0.00 |  | 0.01 | 0.01 * | 0.01 * |  |  |
|  | (-0.00-0.01) |  | (-0.01-0.02) | (0.00-0.01) | (0.00-0.01) |  |  |
|  | (0.00) |  | (0.01) | (0.00) | (0.00) |  |  |
| Elevation | 0.00 |  | 0.02 * | 0.01 ** | 0.01 ** |  |  |
|  | (-0.01-0.01) |  | (0.00-0.03) | (0.00-0.02) | (0.00-0.02) |  |  |
|  | (0.00) |  | (0.01) | (0.00) | (0.00) |  |  |
| EVI |  | -0.01 * | -0.01 * |  | -0.00 | -0.00 |  |
|  |  | (-0.02--0.00) | (-0.02--0.00) |  | (-0.01-0.00) | (-0.01--0.00) |  |
|  |  | (0.01) | (0.01) |  | (0.00) | (0.00) |  |
| AgeClss3 |  |  |  | -0.06 *** | -0.06 *** | -0.04 *** | -0.04 *** |
|  |  |  |  | (-0.09--0.04) | (-0.09--0.04) | (-0.05--0.03) | (-0.05--0.03) |
|  |  |  |  | (0.01) | (0.01) | (0.01) | (0.01) |
| AgeClss4 |  |  |  | -0.01 | -0.02 | -0.04 *** | -0.03 *** |
|  |  |  |  | (-0.03-0.01) | (-0.05-0.01) | (-0.06--0.02) | (-0.05--0.02) |
|  |  |  |  | (0.01) | (0.01) | (0.01) | (0.01) |

|  |  |  |  |  |
| --- | --- | --- | --- | --- |
| AgeCls5 | -0.02 * | -0.02 * | -0.04 *** | -0.03 *** |
|  | (-0.04--0.00) | (-0.05--0.00) | (-0.05--0.02) | (-0.05--0.02) |
|  | (0.01) | (0.01) | (0.01) | (0.01) |
| HghtCls3 | 0.02 * | 0.02 * | -0.00 | -0.00 |
|  | (0.00-0.04) | (0.00-0.04) | (-0.01-0.01) | (-0.01-0.01) |
|  | (0.01) | (0.01) | (0.01) | (0.01) |
| HghtCls4 | 0.00 | 0.00 | -0.00 | -0.01 |
|  | (-0.02-0.02) | (-0.02-0.02) | (-0.02-0.01) | (-0.02-0.01) |
|  | (0.01) | (0.01) | (0.01) | (0.01) |
| HghtCls5 | 0.02 | 0.02 * | 0.00 | -0.00 |
|  | (-0.00-0.04) | (0.00-0.05) | (-0.01-0.02) | (-0.02-0.01) |
|  | (0.01) | (0.01) | (0.01) | (0.01) |
| CrwnDns2 | 0.01 | 0.01 | -0.00 | 0.00 |
|  | (-0.01-0.03) | (-0.01-0.03) | (-0.01-0.01) | (-0.01-0.02) |
|  | (0.01) | (0.01) | (0.01) | (0.01) |
| CrwnDns3 |  |  |  | 0.01 | 0.01 | -0.00 | -0.00 |
|  |  |  |  | (-0.01-0.03) | (-0.01-0.03) | (-0.02-0.01) | (-0.02-0.01) |
|  |  |  |  | (0.01) | (0.01) | (0.01) | (0.01) |
| CrwnDns4 |  |  |  | -0.00 | -0.00 | -0.01 | -0.01 |
|  |  |  |  | (-0.03-0.03) | (-0.03-0.03) | (-0.03-0.01) | (-0.03-0.01) |
|  |  |  |  | (0.02) | (0.02) | (0.01) | (0.01) |
| Observations | 95 | 91 | 91 | 157 | 157 | 160 | 160 |

## Appendix 15

**Table A8.**
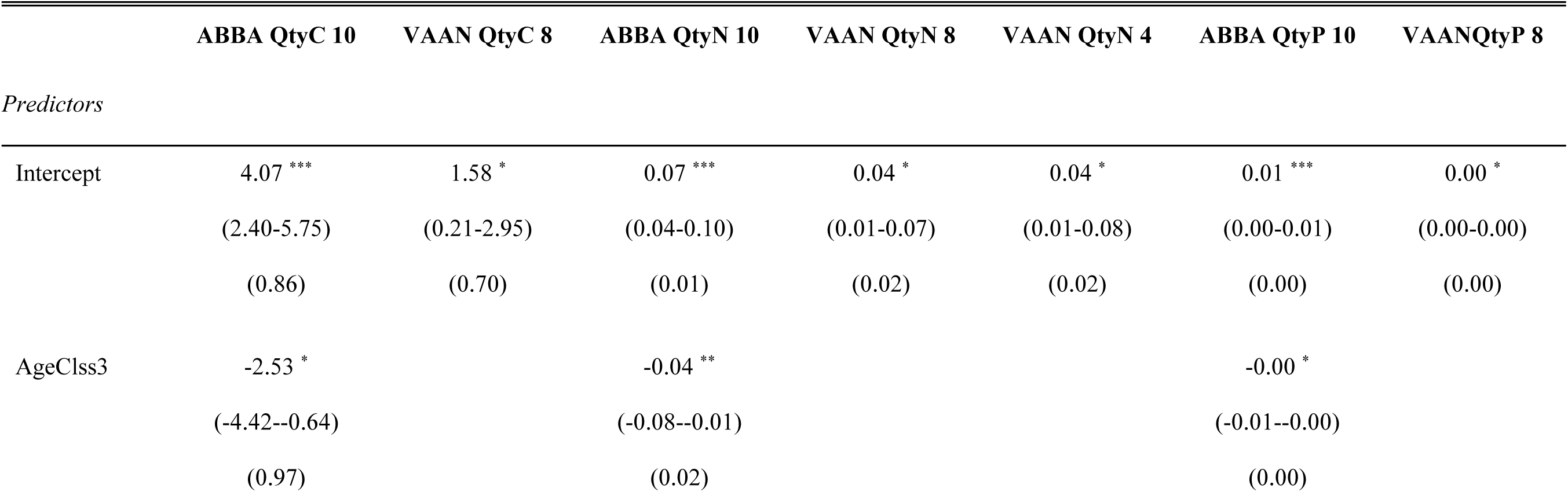
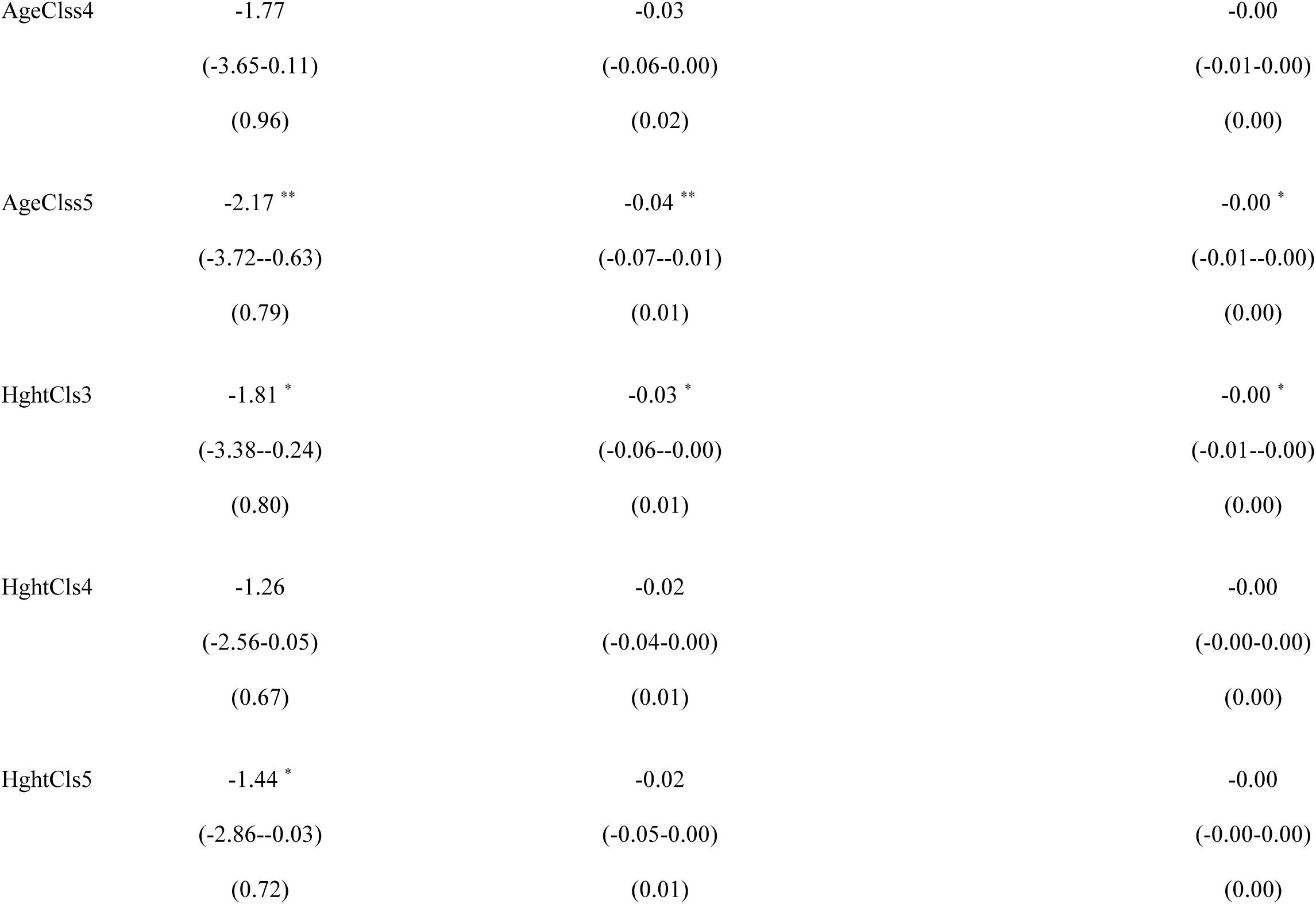
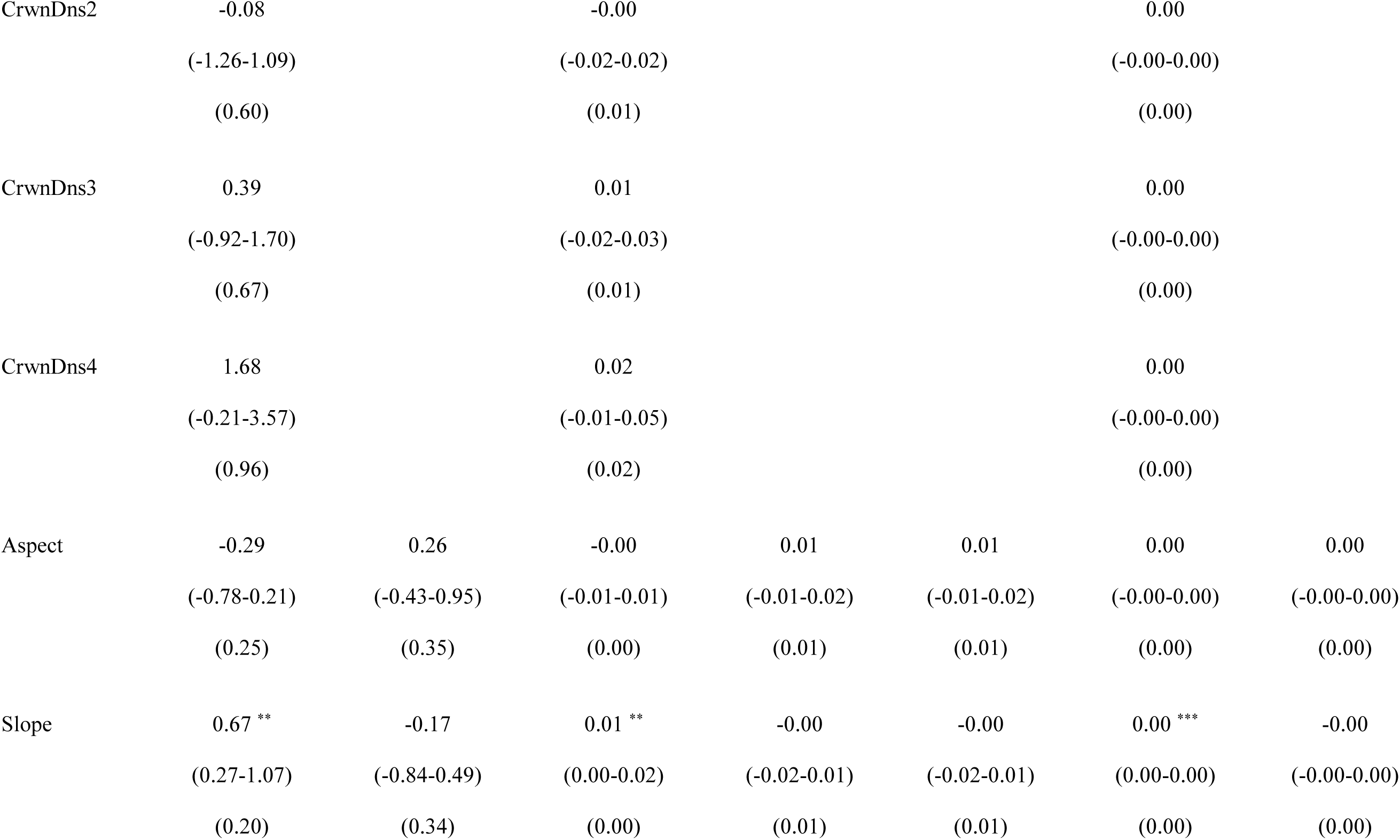
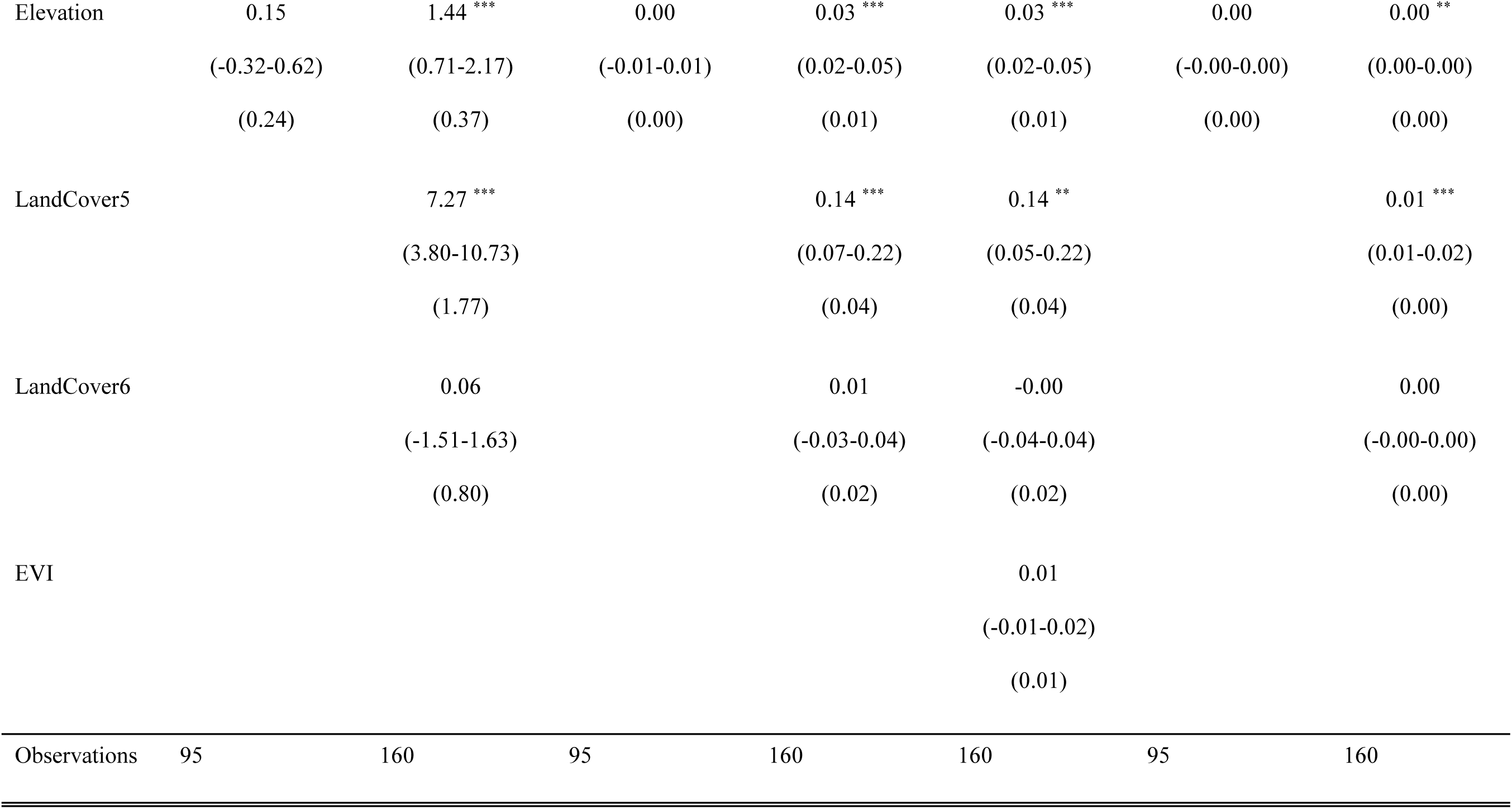
Foliar elemental quantity trait coefficient estimates, confidence intervals, and standard error values for top ranked models (< 2 ΔAIC_c_). Species codes are used for balsam fir (ABBA), red maple (ACRU), white birch (BEPA), black spruce (PIMA, and lowbush blueberry (VAAN). If there is more than one top ranked model per species, we present in order of ΔAIC_c_ rank. Model numbers are supplied beside the species code in the top row (see Table 1 for model descriptions). Predictors include land cover (LandCover5 and LandCover6 represent deciduous and mix wood conditions), EVI (i.e., proxy for productivity), abiotic factors (aspect, slope, elevation), and biotic factors: AgeClss3 (41-60 years old), AgeClss 4 (61-80 years old), AgeClss5 (81-100 years old), HghtCls3 (6.6-9.5 m), HghtCls4 (9.6-12.5 m), HghtCls5 (12.6-15.5m), CrwnDns2 (51-75 % closed), CrwnDns3 (26-50%), CrwnDn4 (10-25 % closed). In addition, asterisks are used to indicate coefficient significance as follows: *\* p<0.05 ** p<0.01 *** p<0.001*.

|  | ABBA QtyC 10 | VAAN QtyC 8 | ABBA QtyN 10 | VAAN QtyN 8 | VAAN QtyN 4 | ABBA QtyP 10 | VAANQtyP 8 |
| --- | --- | --- | --- | --- | --- | --- | --- |
| *Predictors* |  |  |  |  |  |  |  |
| Intercept | 4.07 ***<br>(2.40-5.75)<br>(0.86) | 1.58 *<br>(0.21-2.95)<br>(0.70) | 0.07 ***<br>(0.04-0.10)<br>(0.01) | 0.04 *<br>(0.01-0.07)<br>(0.02) | 0.04 *<br>(0.01-0.08)<br>(0.02) | 0.01 ***<br>(0.00-0.01)<br>(0.00) | 0.00 *<br>(0.00-0.00)<br>(0.00) |
| AgeClss3 | -2.53 *<br>(-4.42--0.64)<br>(0.97) |  | -0.04 **<br>(-0.08--0.01)<br>(0.02) |  |  | -0.00 *<br>(-0.01--0.00)<br>(0.00) |  |

|  |  |  |  |
| --- | --- | --- | --- |
| AgeClss4 | -1.77<br>(-3.65-0.11)<br>(0.96) | -0.03<br>(-0.06-0.00)<br>(0.02) | -0.00<br>(-0.01-0.00)<br>(0.00) |
| AgeClss5 | -2.17 **<br>(-3.72--0.63)<br>(0.79) | -0.04 **<br>(-0.07--0.01)<br>(0.01) | -0.00 *<br>(-0.01--0.00)<br>(0.00) |
| HghtCls3 | -1.81 *<br>(-3.38--0.24)<br>(0.80) | -0.03 *<br>(-0.06--0.00)<br>(0.01) | -0.00 *<br>(-0.01--0.00)<br>(0.00) |
| HghtCls4 | -1.26<br>(-2.56-0.05)<br>(0.67) | -0.02<br>(-0.04-0.00)<br>(0.01) | -0.00<br>(-0.00-0.00)<br>(0.00) |
| HghtCls5 | -1.44 *<br>(-2.86--0.03)<br>(0.72) | -0.02<br>(-0.05-0.00)<br>(0.01) | -0.00<br>(-0.00-0.00)<br>(0.00) |
| CrwnDns2 | -0.08 |  | -0.00 |  |  | 0.00 |  |
|  | (-1.26-1.09) |  | (-0.02-0.02) |  |  | (-0.00-0.00) |  |
|  | (0.60) |  | (0.01) |  |  | (0.00) |  |
| CrwnDns3 | 0.39 |  | 0.01 |  |  | 0.00 |  |
|  | (-0.92-1.70) |  | (-0.02-0.03) |  |  | (-0.00-0.00) |  |
|  | (0.67) |  | (0.01) |  |  | (0.00) |  |
| CrwnDns4 | 1.68 |  | 0.02 |  |  | 0.00 |  |
|  | (-0.21-3.57) |  | (-0.01-0.05) |  |  | (-0.00-0.00) |  |
|  | (0.96) |  | (0.02) |  |  | (0.00) |  |
| Aspect | -0.29 | 0.26 | -0.00 | 0.01 | 0.01 | 0.00 | 0.00 |
|  | (-0.78-0.21) | (-0.43-0.95) | (-0.01-0.01) | (-0.01-0.02) | (-0.01-0.02) | (-0.00-0.00) | (-0.00-0.00) |
|  | (0.25) | (0.35) | (0.00) | (0.01) | (0.01) | (0.00) | (0.00) |
| Slope | 0.67 ** | -0.17 | 0.01 ** | -0.00 | -0.00 | 0.00 *** | -0.00 |
|  | (0.27-1.07) | (-0.84-0.49) | (0.00-0.02) | (-0.02-0.01) | (-0.02-0.01) | (0.00-0.00) | (-0.00-0.00) |
|  | (0.20) | (0.34) | (0.00) | (0.01) | (0.01) | (0.00) | (0.00) |
| Elevation | 0.15 | 1.44 *** | 0.00 | 0.03 *** | 0.03 *** | 0.00 | 0.00 ** |
|  | (-0.32-0.62) | (0.71-2.17) | (-0.01-0.01) | (0.02-0.05) | (0.02-0.05) | (-0.00-0.00) | (0.00-0.00) |
|  | (0.24) | (0.37) | (0.00) | (0.01) | (0.01) | (0.00) | (0.00) |
| LandCover5 |  | 7.27 *** |  | 0.14 *** | 0.14 ** |  | 0.01 *** |
|  |  | (3.80-10.73) |  | (0.07-0.22) | (0.05-0.22) |  | (0.01-0.02) |
|  |  | (1.77) |  | (0.04) | (0.04) |  | (0.00) |
| LandCover6 |  | 0.06 |  | 0.01 | -0.00 |  | 0.00 |
|  |  | (-1.51-1.63) |  | (-0.03-0.04) | (-0.04-0.04) |  | (-0.00-0.00) |
|  |  | (0.80) |  | (0.02) | (0.02) |  | (0.00) |
| EVI |  |  |  |  | 0.01 |  |  |
|  |  |  |  |  | (-0.01-0.02) |  |  |
|  |  |  |  |  | (0.01) |  |  |
| Observations | 95 | 160 | 95 | 160 | 160 | 95 | 160 |

## Appendix 16

**Table A9.**
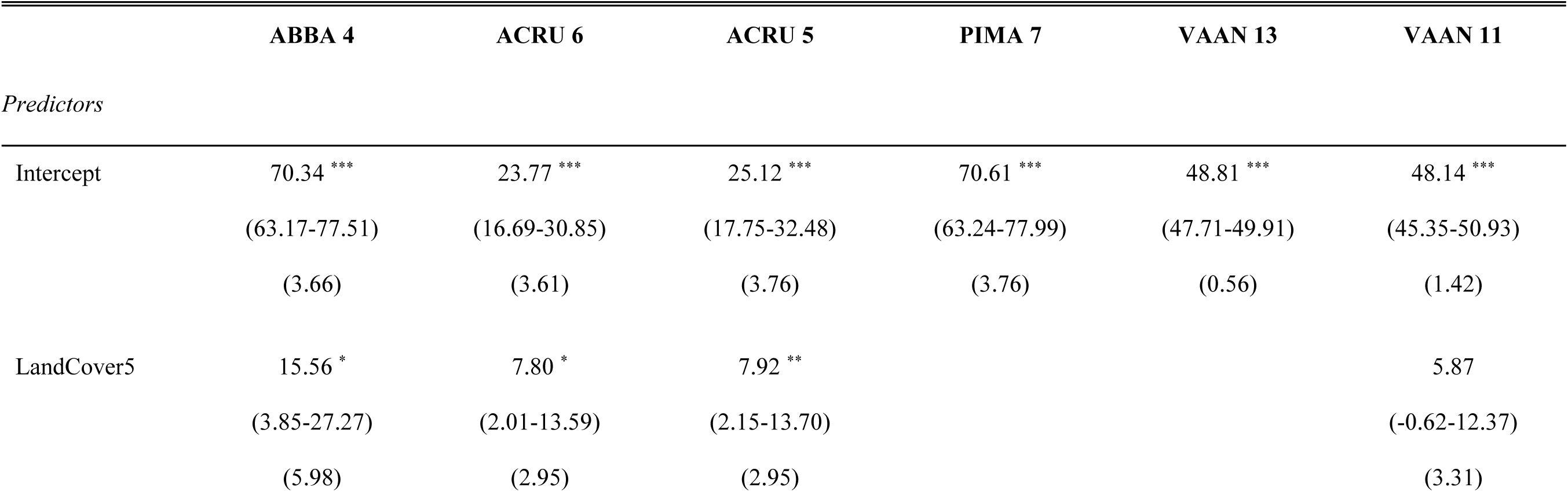
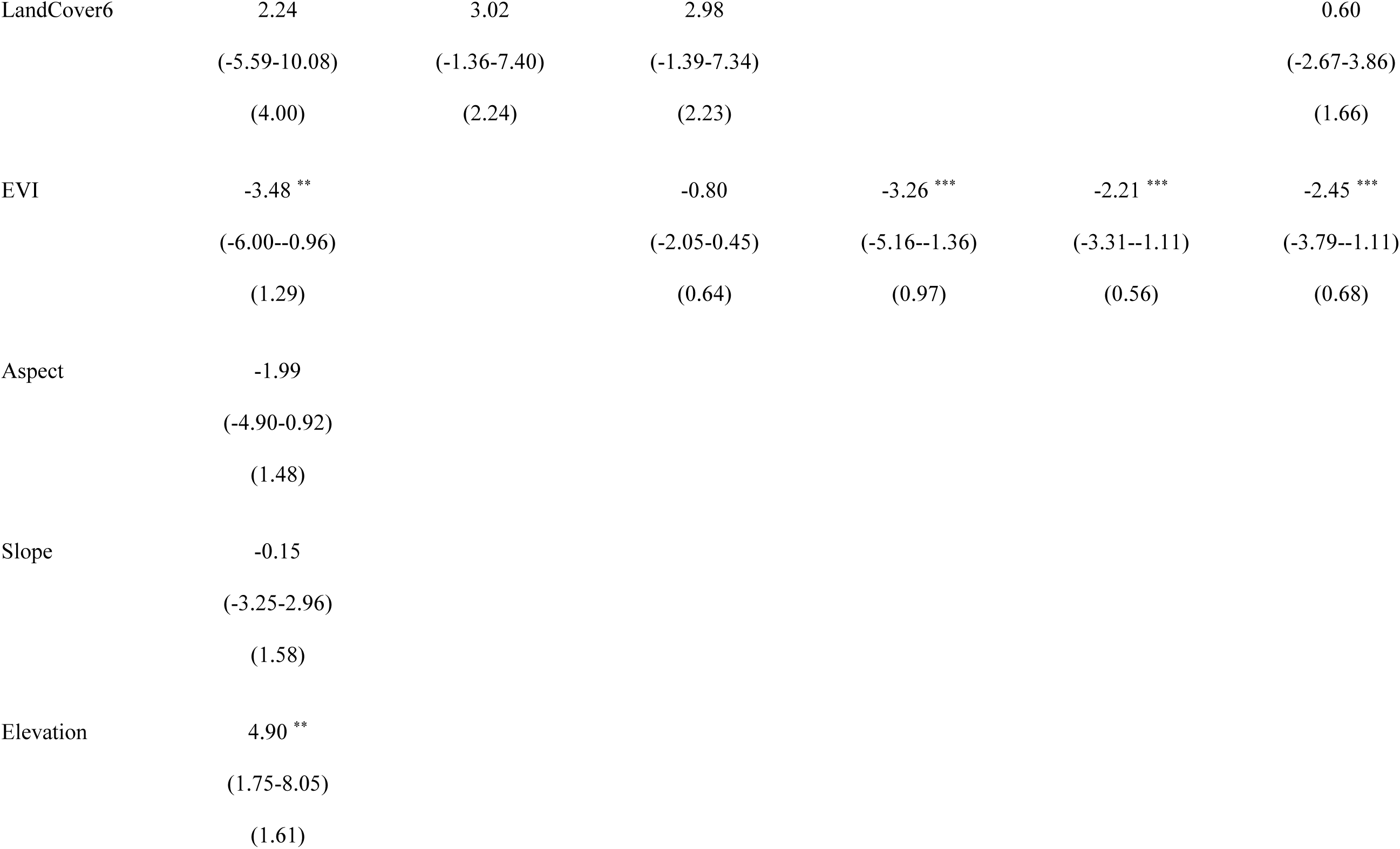
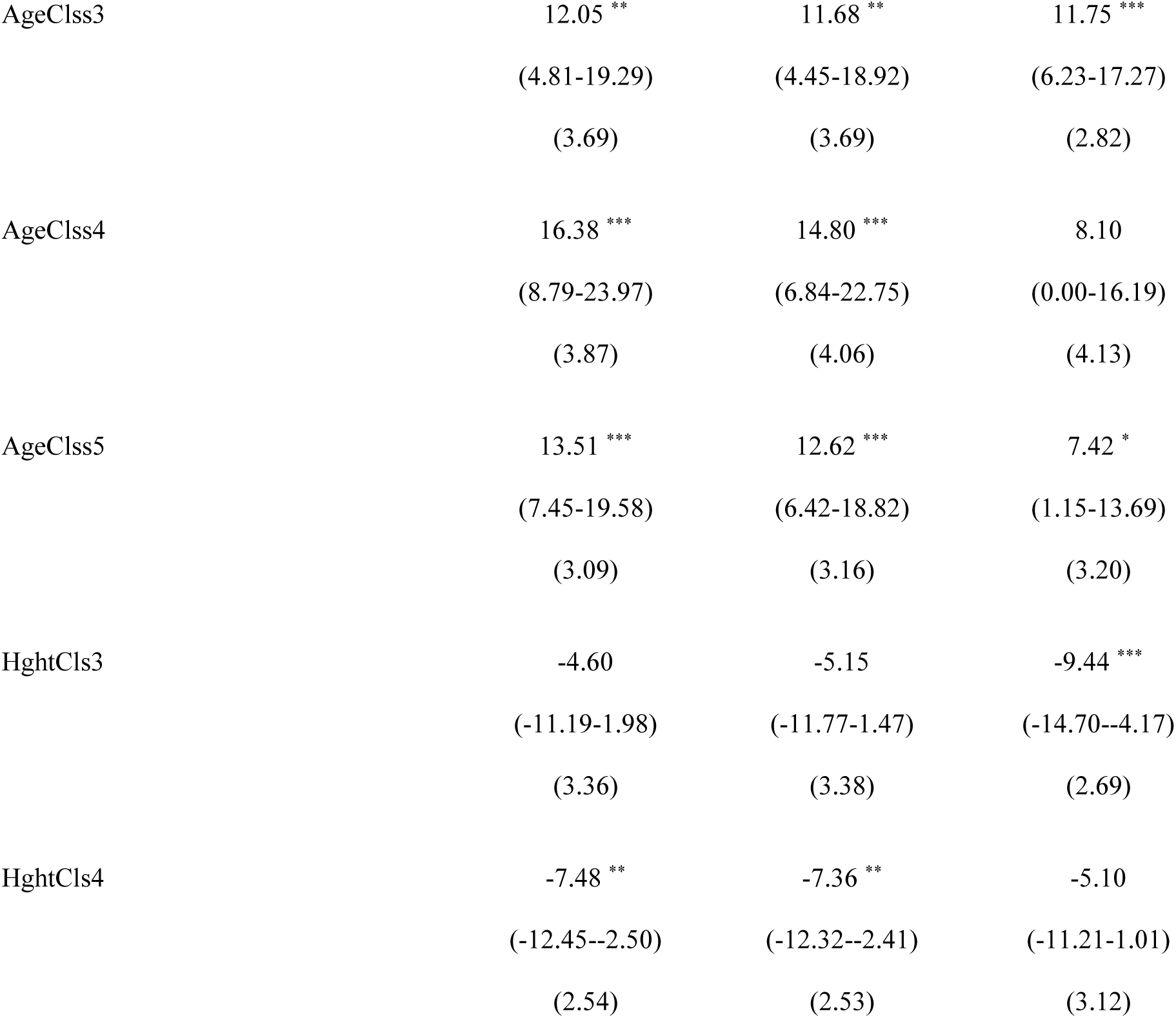
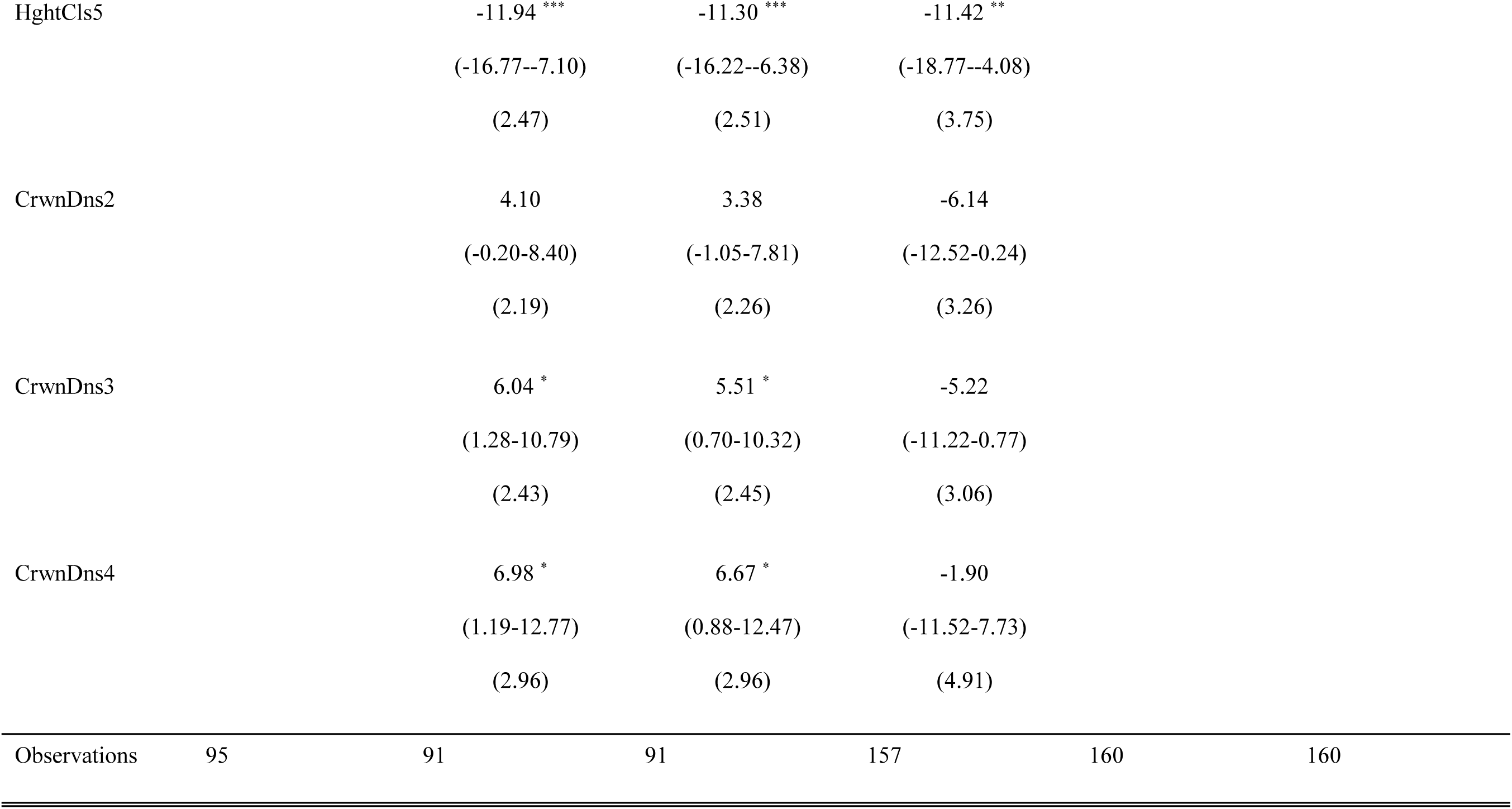
Foliar stoichiometric C:N trait coefficient estimates, confidence intervals, and standard error values for top ranked models (< 2 ΔAIC_c_). Species codes are used for balsam fir (ABBA), red maple (ACRU), white birch (BEPA), black spruce (PIMA, and lowbush blueberry (VAAN). If there is more than one top ranked model per species, we present in order of ΔAIC_c_ rank. Model numbers are supplied beside the species code in the top row (see Table 1 for model descriptions). Predictors include land cover (LandCover5 and LandCover6 represent deciduous and mix wood conditions), EVI (i.e., proxy for productivity), abiotic factors (aspect, slope, elevation), and biotic factors: AgeClss3 (41-60 years old), AgeClss 4 (61-80 years old), AgeClss5 (81-100 years old), HghtCls3 (6.6-9.5 m), HghtCls4 (9.6-12.5 m), HghtCls5 (12.6-15.5m), CrwnDns2 (51-75 % closed), CrwnDns3 (26-50%), CrwnDn4 (10-25 % closed). Total number of observations are provided in the bottom row. In addition, asterisks are used to indicate coefficient significance as follows: *\* p<0.05 ** p<0.01 *** p<0.001*.

|  | ABBA 4 | ACRU 6 | ACRU 5 | PIMA 7 | VAAN 13 | VAAN 11 |
| --- | --- | --- | --- | --- | --- | --- |
| *Predictors* |  |  |  |  |  |  |
| Intercept | 70.34 ***<br>(63.17-77.51)<br>(3.66) | 23.77 ***<br>(16.69-30.85)<br>(3.61) | 25.12 ***<br>(17.75-32.48)<br>(3.76) | 70.61 ***<br>(63.24-77.99)<br>(3.76) | 48.81 ***<br>(47.71-49.91)<br>(0.56) | 48.14 ***<br>(45.35-50.93)<br>(1.42) |
| LandCover5 | 15.56 *<br>(3.85-27.27)<br>(5.98) | 7.80 *<br>(2.01-13.59)<br>(2.95) | 7.92 **<br>(2.15-13.70)<br>(2.95) |  |  | 5.87<br>(-0.62-12.37)<br>(3.31) |
| LandCover6 | 2.24 | 3.02 | 2.98 |  |  | 0.60 |
|  | (-5.59-10.08) | (-1.36-7.40) | (-1.39-7.34) |  |  | (-2.67-3.86) |
|  | (4.00) | (2.24) | (2.23) |  |  | (1.66) |
| EVI | -3.48 ** |  | -0.80 | -3.26 *** | -2.21 *** | -2.45 *** |
|  | (-6.00--0.96) |  | (-2.05-0.45) | (-5.16--1.36) | (-3.31--1.11) | (-3.79--1.11) |
|  | (1.29) |  | (0.64) | (0.97) | (0.56) | (0.68) |
| Aspect | -1.99 |  |  |  |  |  |
|  | (-4.90-0.92) |  |  |  |  |  |
|  | (1.48) |  |  |  |  |  |
| Slope | -0.15 |  |  |  |  |  |
|  | (-3.25-2.96) |  |  |  |  |  |
|  | (1.58) |  |  |  |  |  |
| Elevation | 4.90 ** |  |  |  |  |  |
|  | (1.75-8.05) |  |  |  |  |  |
|  | (1.61) |  |  |  |  |  |

|  |  |  |  |
| --- | --- | --- | --- |
| AgeCls3 | 12.05 **<br>(4.81-19.29)<br>(3.69) | 11.68 **<br>(4.45-18.92)<br>(3.69) | 11.75 ***<br>(6.23-17.27)<br>(2.82) |
| AgeCls4 | 16.38 ***<br>(8.79-23.97)<br>(3.87) | 14.80 ***<br>(6.84-22.75)<br>(4.06) | 8.10<br>(0.00-16.19)<br>(4.13) |
| AgeCls5 | 13.51 ***<br>(7.45-19.58)<br>(3.09) | 12.62 ***<br>(6.42-18.82)<br>(3.16) | 7.42 *<br>(1.15-13.69)<br>(3.20) |
| HghtCls3 | -4.60<br>(-11.19-1.98)<br>(3.36) | -5.15<br>(-11.77-1.47)<br>(3.38) | -9.44 ***<br>(-14.70--4.17)<br>(2.69) |
| HghtCls4 | -7.48 **<br>(-12.45--2.50)<br>(2.54) | -7.36 **<br>(-12.32--2.41)<br>(2.53) | -5.10<br>(-11.21-1.01)<br>(3.12) |
| HghtCls5 |  | -11.94 *** | -11.30 *** | -11.42 ** |  |  |
|  |  | (-16.77--7.10) | (-16.22--6.38) | (-18.77--4.08) |  |  |
|  |  | (2.47) | (2.51) | (3.75) |  |  |
| CrwnDns2 |  | 4.10 | 3.38 | -6.14 |  |  |
|  |  | (-0.20-8.40) | (-1.05-7.81) | (-12.52-0.24) |  |  |
|  |  | (2.19) | (2.26) | (3.26) |  |  |
| CrwnDns3 |  | 6.04 * | 5.51 * | -5.22 |  |  |
|  |  | (1.28-10.79) | (0.70-10.32) | (-11.22-0.77) |  |  |
|  |  | (2.43) | (2.45) | (3.06) |  |  |
| CrwnDns4 |  | 6.98 * | 6.67 * | -1.90 |  |  |
|  |  | (1.19-12.77) | (0.88-12.47) | (-11.52-7.73) |  |  |
|  |  | (2.96) | (2.96) | (4.91) |  |  |
| Observations | 95 | 91 | 91 | 157 | 160 | 160 |

## Appendix 17

**Table A10.**
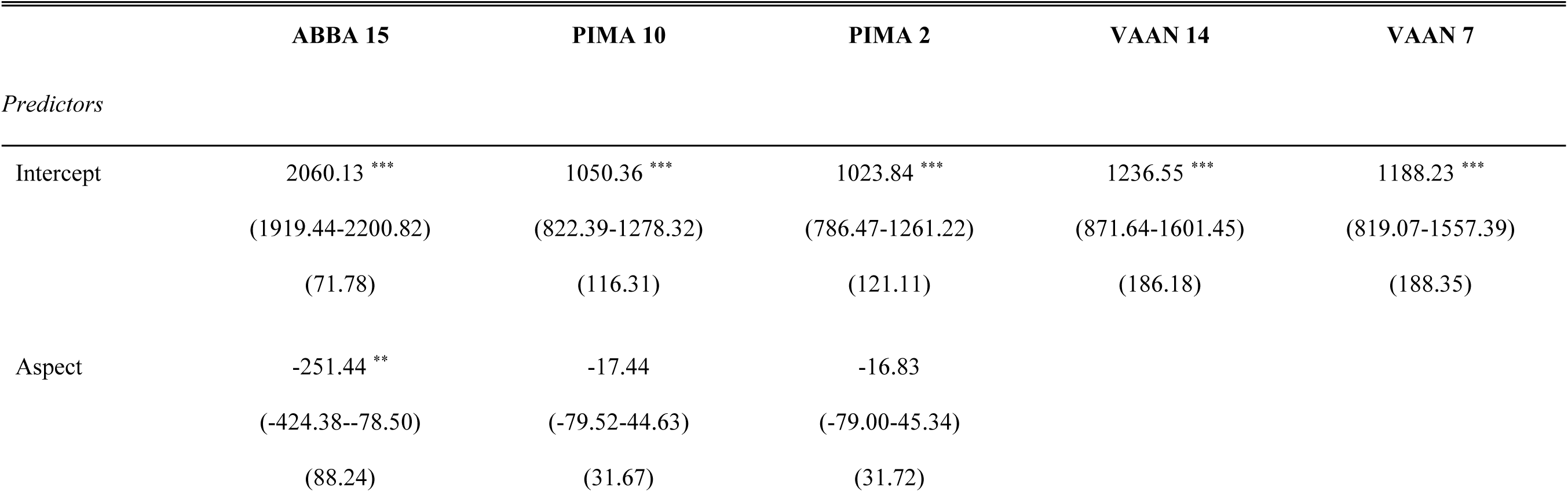
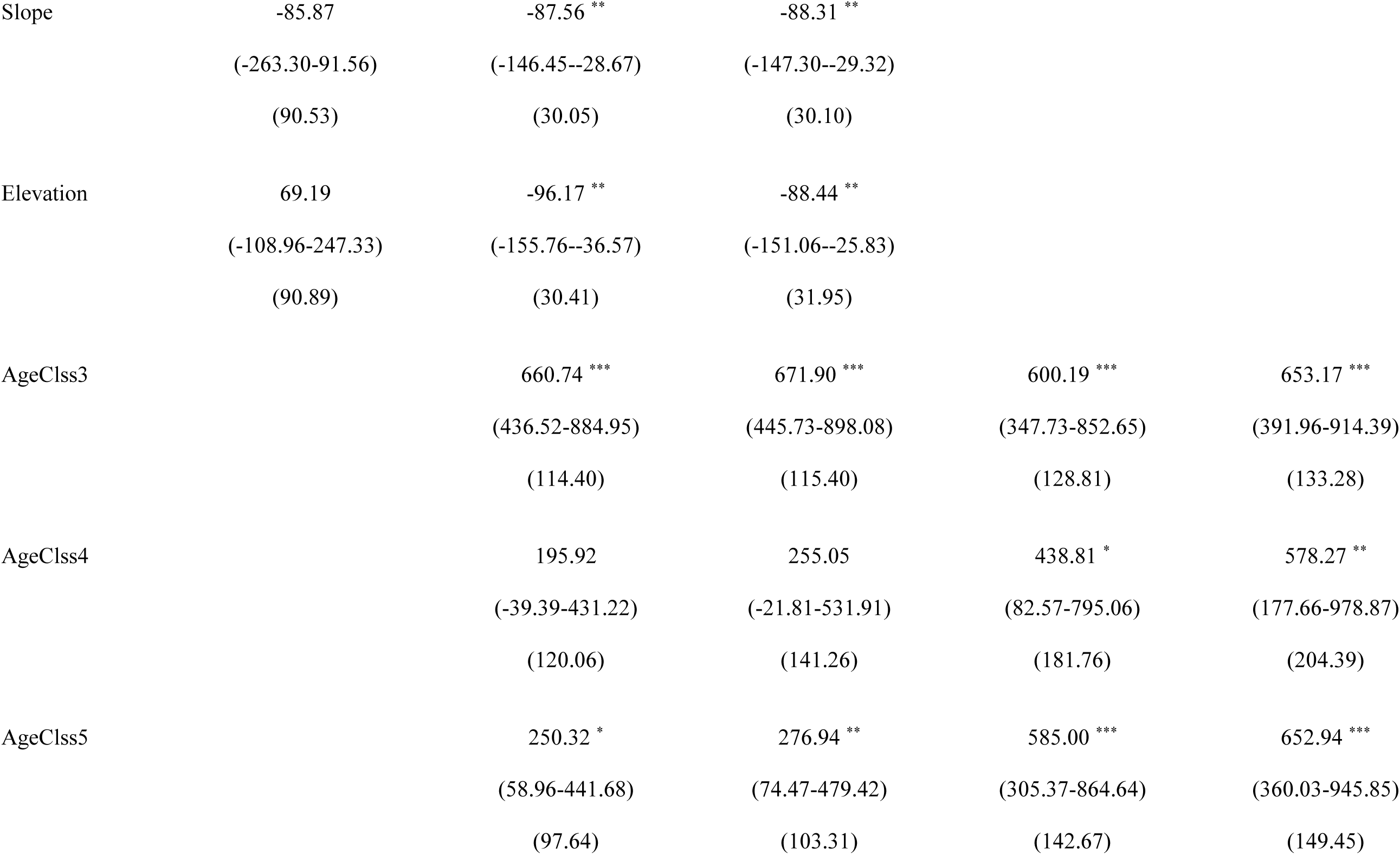
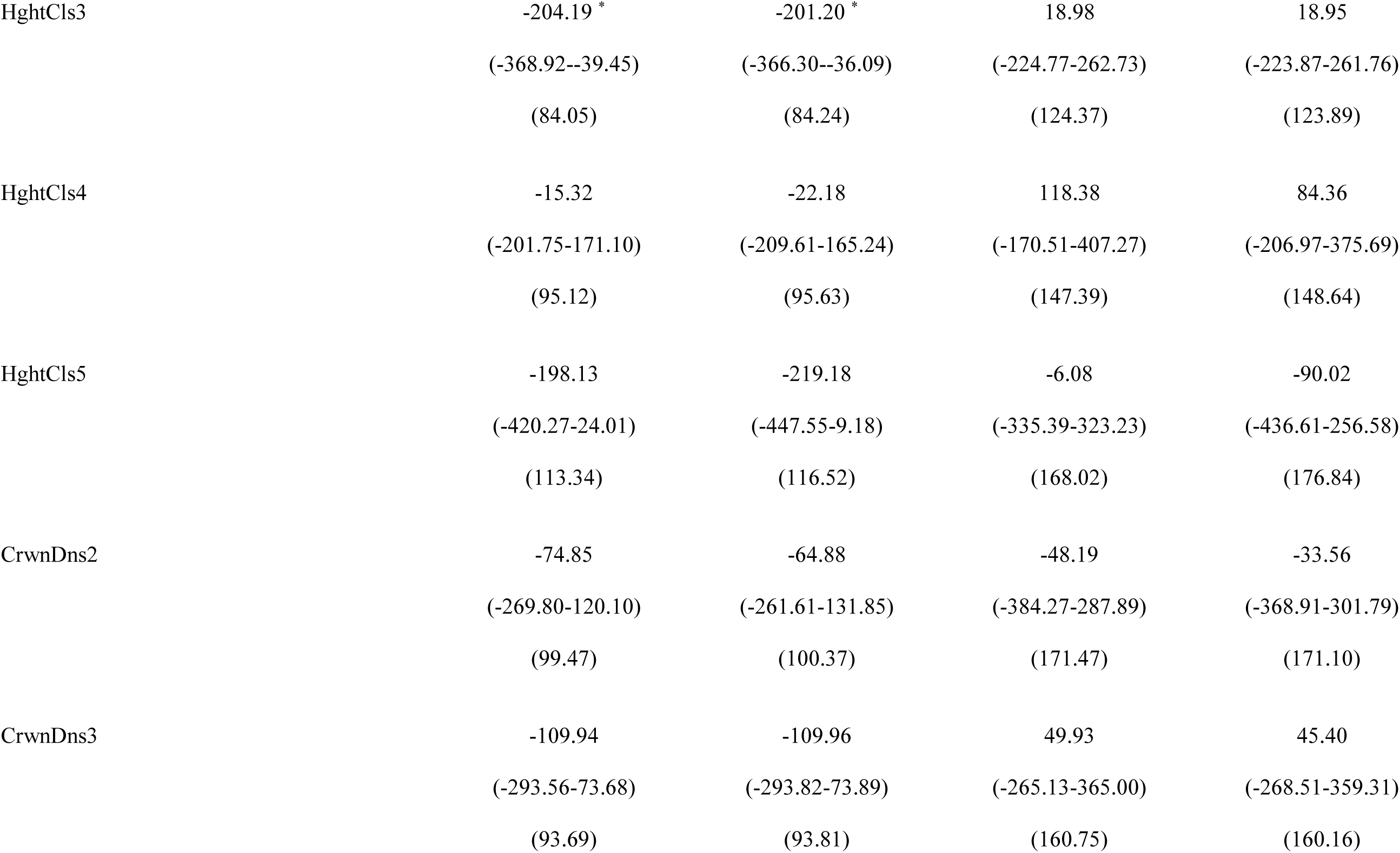
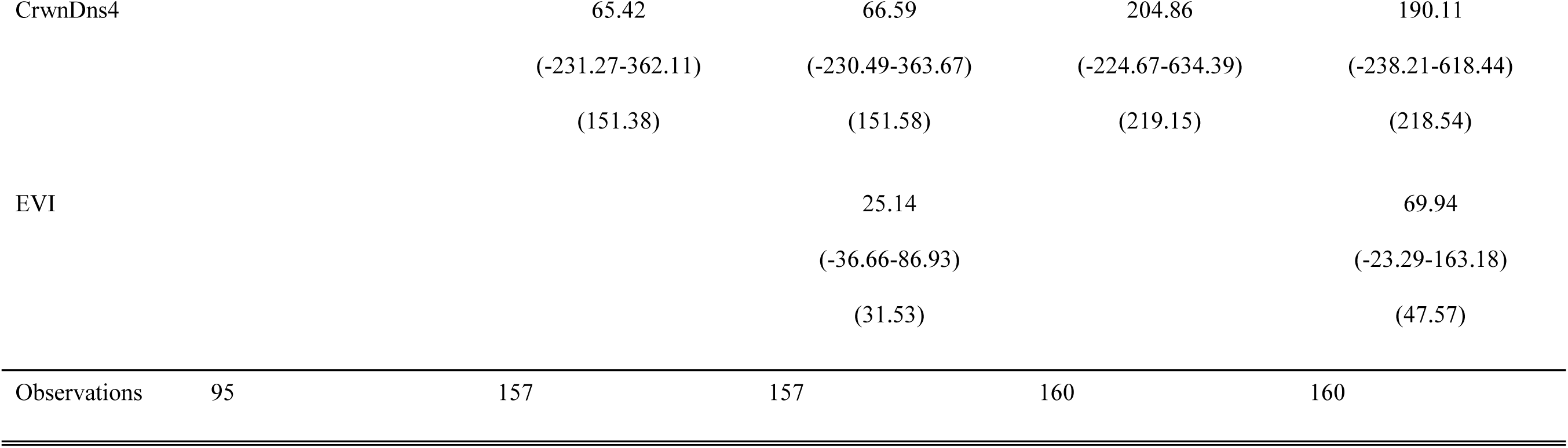
Foliar stoichiometric C:P trait coefficient estimates, confidence intervals, and standard error values for top ranked models (< 2 ΔAIC_c_). Species codes are used for balsam fir (ABBA), red maple (ACRU), white birch (BEPA), black spruce (PIMA, and lowbush blueberry (VAAN). If there is more than one top ranked model per species, we present in order of ΔAIC_c_ rank. Model numbers are supplied beside the species code in the top row (see Table 1 for model descriptions). Predictors include land cover (LandCover5 and LandCover6 represent deciduous and mix wood conditions), EVI (i.e., proxy for productivity), abiotic factors (aspect, slope, elevation), and biotic factors: AgeClss3 (41-60 years old), AgeClss 4 (61-80 years old), AgeClss5 (81-100 years old), HghtCls3 (6.6-9.5 m), HghtCls4 (9.6-12.5 m), HghtCls5 (12.6-15.5m), CrwnDns2 (51-75 % closed), CrwnDns3 (26-50%), CrwnDn4 (10-25 % closed). Total number of observations are provided in the bottom row. In addition, asterisks are used to indicate coefficient significance as follows: *\* p<0.05 ** p<0.01 *** p<0.001*.

|  | ABBA 15 | PIMA 10 | PIMA 2 | VAAN 14 | VAAN 7 |
| --- | --- | --- | --- | --- | --- |
| *Predictors* |  |  |  |  |  |
| Intercept | 2060.13 ***<br>(1919.44-2200.82)<br>(71.78) | 1050.36 ***<br>(822.39-1278.32)<br>(116.31) | 1023.84 ***<br>(786.47-1261.22)<br>(121.11) | 1236.55 ***<br>(871.64-1601.45)<br>(186.18) | 1188.23 ***<br>(819.07-1557.39)<br>(188.35) |
| Aspect | -251.44 **<br>(-424.38--78.50)<br>(88.24) | -17.44<br>(-79.52-44.63)<br>(31.67) | -16.83<br>(-79.00-45.34)<br>(31.72) |  |  |
| Slope | -85.87<br>(-263.30-91.56)<br>(90.53) | -87.56 **<br>(-146.45--28.67)<br>(30.05) | -88.31 **<br>(-147.30--29.32)<br>(30.10) |  |  |
| Elevation | 69.19<br>(-108.96-247.33)<br>(90.89) | -96.17 **<br>(-155.76--36.57)<br>(30.41) | -88.44 **<br>(-151.06--25.83)<br>(31.95) |  |  |
| AgeClss3 |  | 660.74 ***<br>(436.52-884.95)<br>(114.40) | 671.90 ***<br>(445.73-898.08)<br>(115.40) | 600.19 ***<br>(347.73-852.65)<br>(128.81) | 653.17 ***<br>(391.96-914.39)<br>(133.28) |
| AgeClss4 |  | 195.92<br>(-39.39-431.22)<br>(120.06) | 255.05<br>(-21.81-531.91)<br>(141.26) | 438.81 *<br>(82.57-795.06)<br>(181.76) | 578.27 **<br>(177.66-978.87)<br>(204.39) |
| AgeClss5 |  | 250.32 *<br>(58.96-441.68)<br>(97.64) | 276.94 **<br>(74.47-479.42)<br>(103.31) | 585.00 ***<br>(305.37-864.64)<br>(142.67) | 652.94 ***<br>(360.03-945.85)<br>(149.45) |

|  |  |  |  |  |
| --- | --- | --- | --- | --- |
| HghtCls3 | -204.19 * | -201.20 * | 18.98 | 18.95 |
|  | (-368.92--39.45) | (-366.30--36.09) | (-224.77-262.73) | (-223.87-261.76) |
|  | (84.05) | (84.24) | (124.37) | (123.89) |
| HghtCls4 | -15.32 | -22.18 | 118.38 | 84.36 |
|  | (-201.75-171.10) | (-209.61-165.24) | (-170.51-407.27) | (-206.97-375.69) |
|  | (95.12) | (95.63) | (147.39) | (148.64) |
| HghtCls5 | -198.13 | -219.18 | -6.08 | -90.02 |
|  | (-420.27-24.01) | (-447.55-9.18) | (-335.39-323.23) | (-436.61-256.58) |
|  | (113.34) | (116.52) | (168.02) | (176.84) |
| CrwnDns2 | -74.85 | -64.88 | -48.19 | -33.56 |
|  | (-269.80-120.10) | (-261.61-131.85) | (-384.27-287.89) | (-368.91-301.79) |
|  | (99.47) | (100.37) | (171.47) | (171.10) |
| CrwnDns3 | -109.94 | -109.96 | 49.93 | 45.40 |
|  | (-293.56-73.68) | (-293.82-73.89) | (-265.13-365.00) | (-268.51-359.31) |
|  | (93.69) | (93.81) | (160.75) | (160.16) |
| CrwnDns4 |  | 65.42 | 66.59 | 204.86 | 190.11 |
|  |  | (-231.27-362.11) | (-230.49-363.67) | (-224.67-634.39) | (-238.21-618.44) |
|  |  | (151.38) | (151.58) | (219.15) | (218.54) |
| EVI |  |  | 25.14 |  | 69.94 |
|  |  |  | (-36.66-86.93) |  | (-23.29-163.18) |
|  |  |  | (31.53) |  | (47.57) |
| Observations | 95 | 157 | 157 | 160 | 160 |

## Appendix 18

**Table A11.**
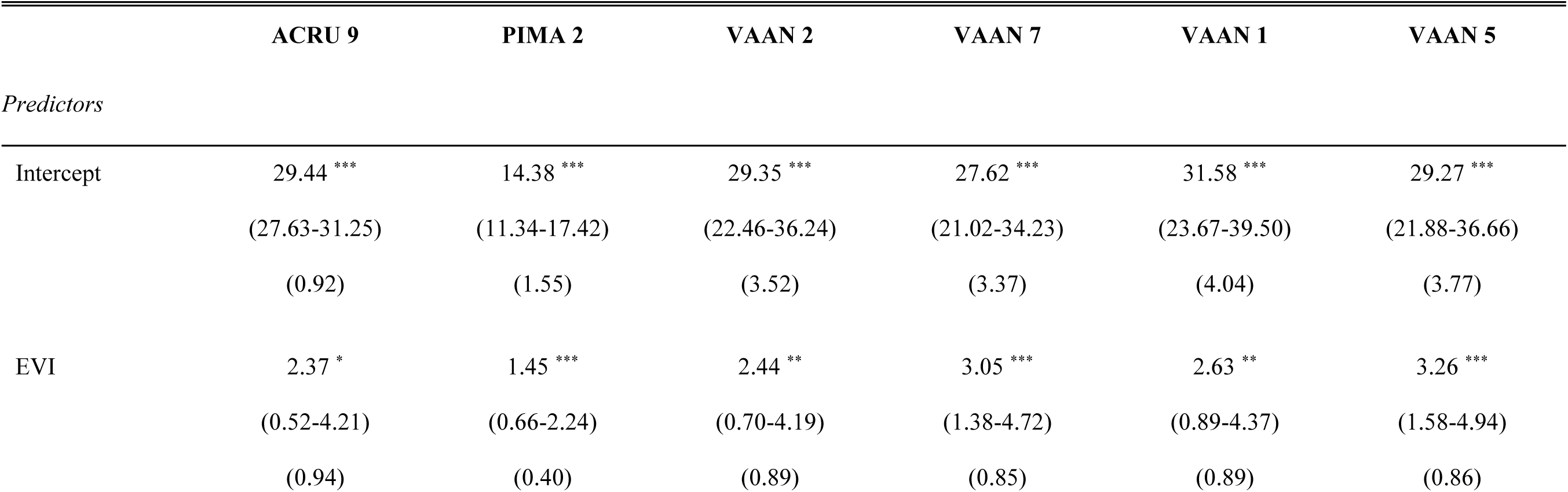
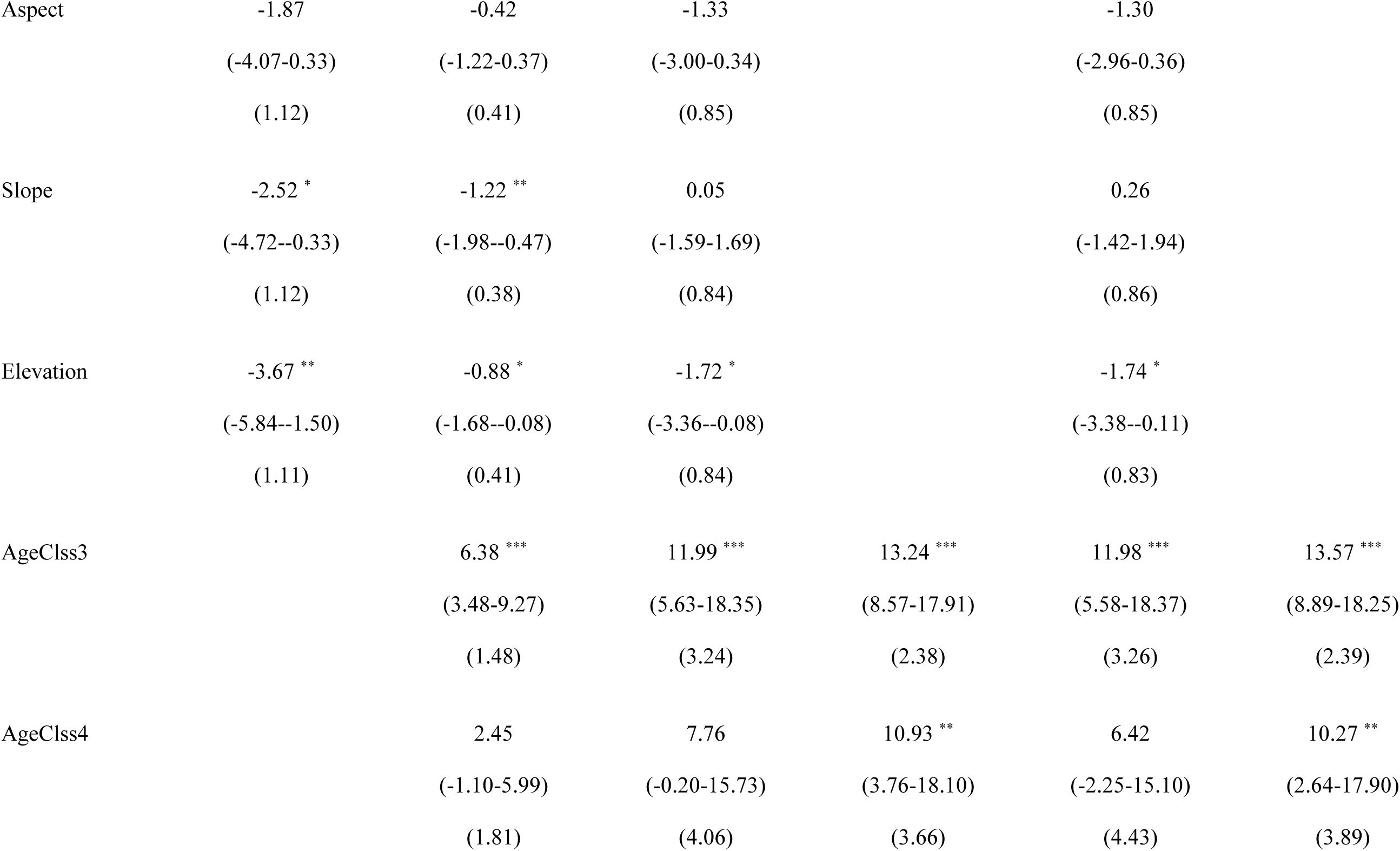
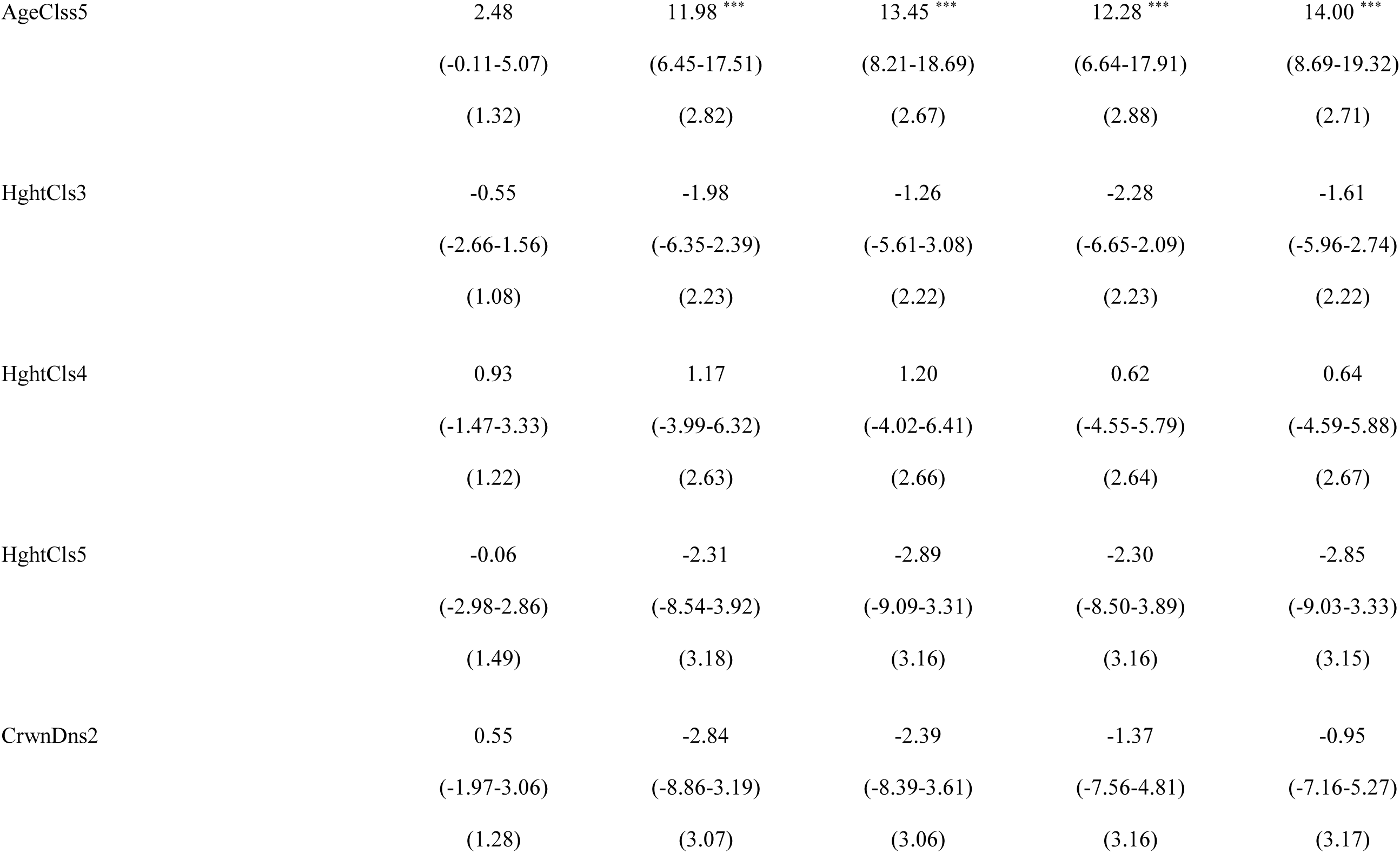
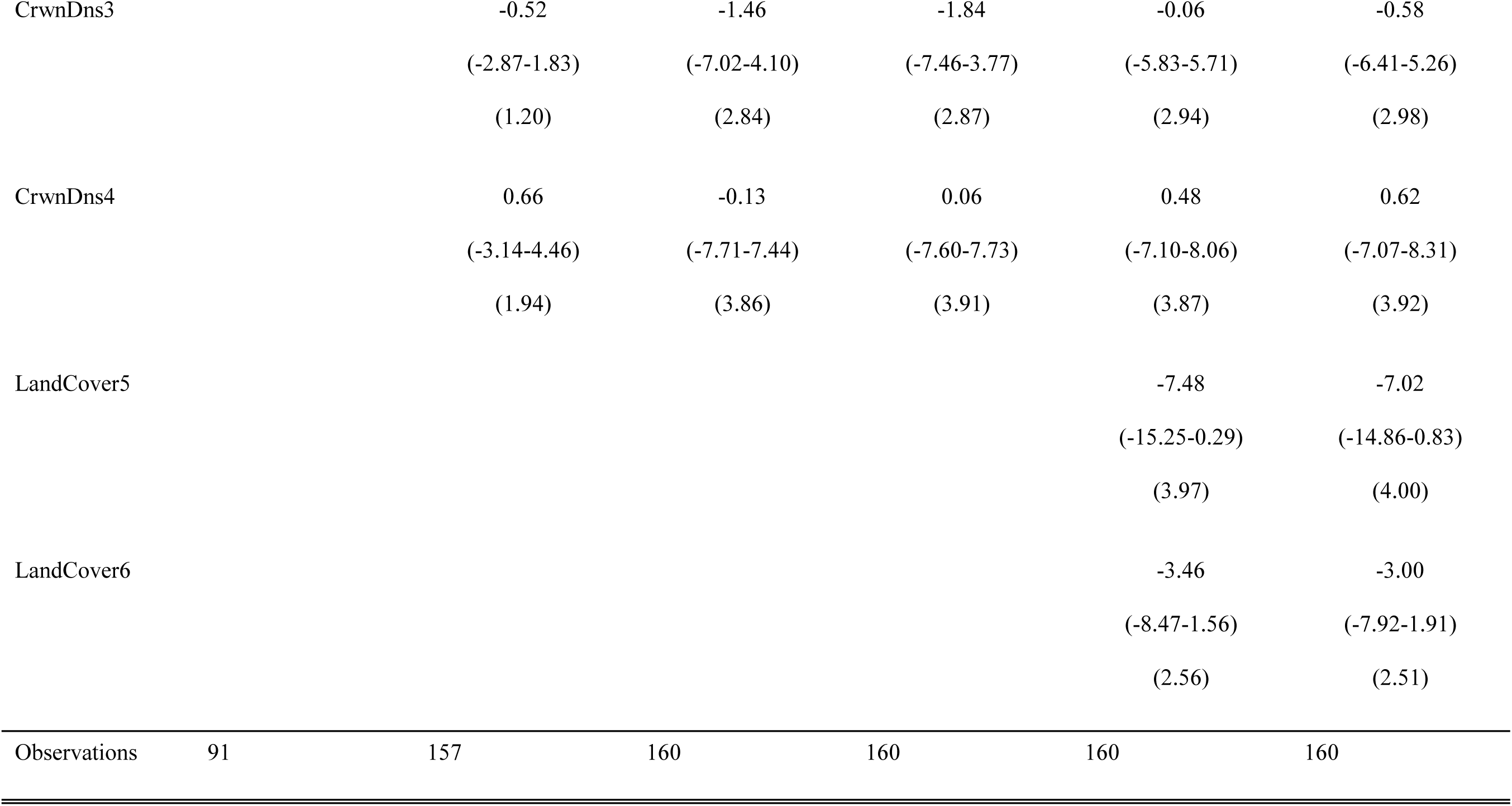
Foliar stoichiometric N:P trait coefficient estimates, confidence intervals, and standard error values for top ranked models (< 2 ΔAIC_c_). Species codes are used for balsam fir (ABBA), red maple (ACRU), white birch (BEPA), black spruce (PIMA, and lowbush blueberry (VAAN). If there is more than one top ranked model per species, we present in order of ΔAIC_c_ rank. Model numbers are supplied beside the species code in the top row (see Table 1 for model descriptions). Predictors include land cover (LandCover5 and LandCover6 represent deciduous and mix wood conditions), EVI (i.e., proxy for productivity), abiotic factors (aspect, slope, elevation), and biotic factors: AgeClss3 (41-60 years old), AgeClss 4 (61-80 years old), AgeClss5 (81-100 years old), HghtCls3 (6.6-9.5 m), HghtCls4 (9.6-12.5 m), HghtCls5 (12.6-15.5m), CrwnDns2 (51-75 % closed), CrwnDns3 (26-50%), CrwnDn4 (10-25 % closed). Total number of observations are provided in the bottom row. In addition, asterisks are used to indicate coefficient significance as follows: *\* p<0.05 ** p<0.01 *** p<0.001*.

## Appendix 19

**Table A12.**
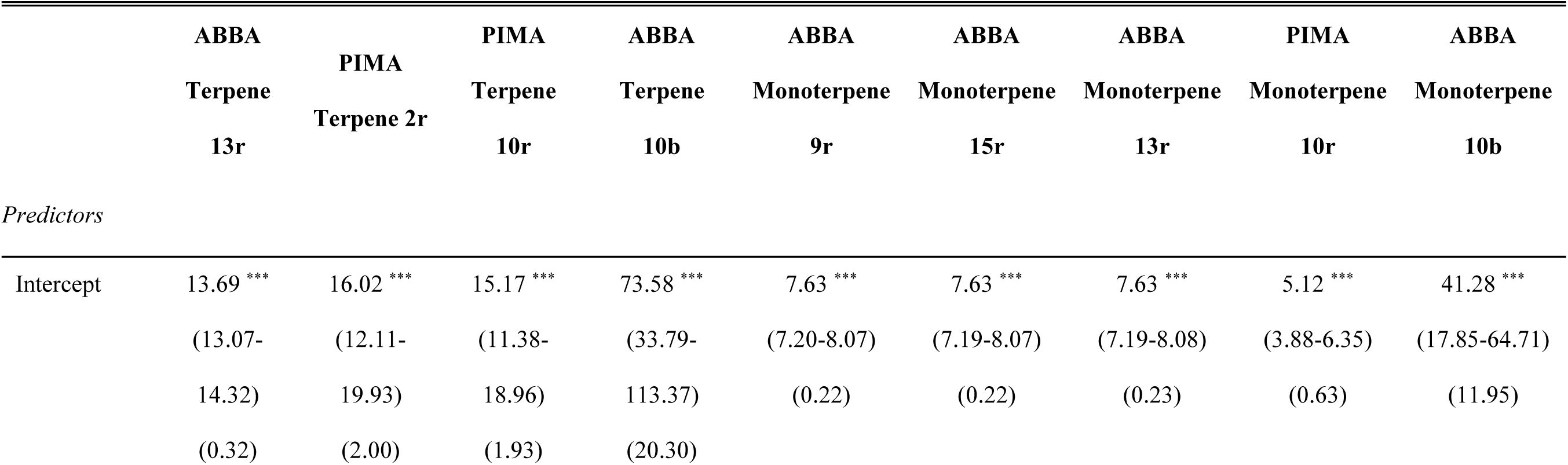
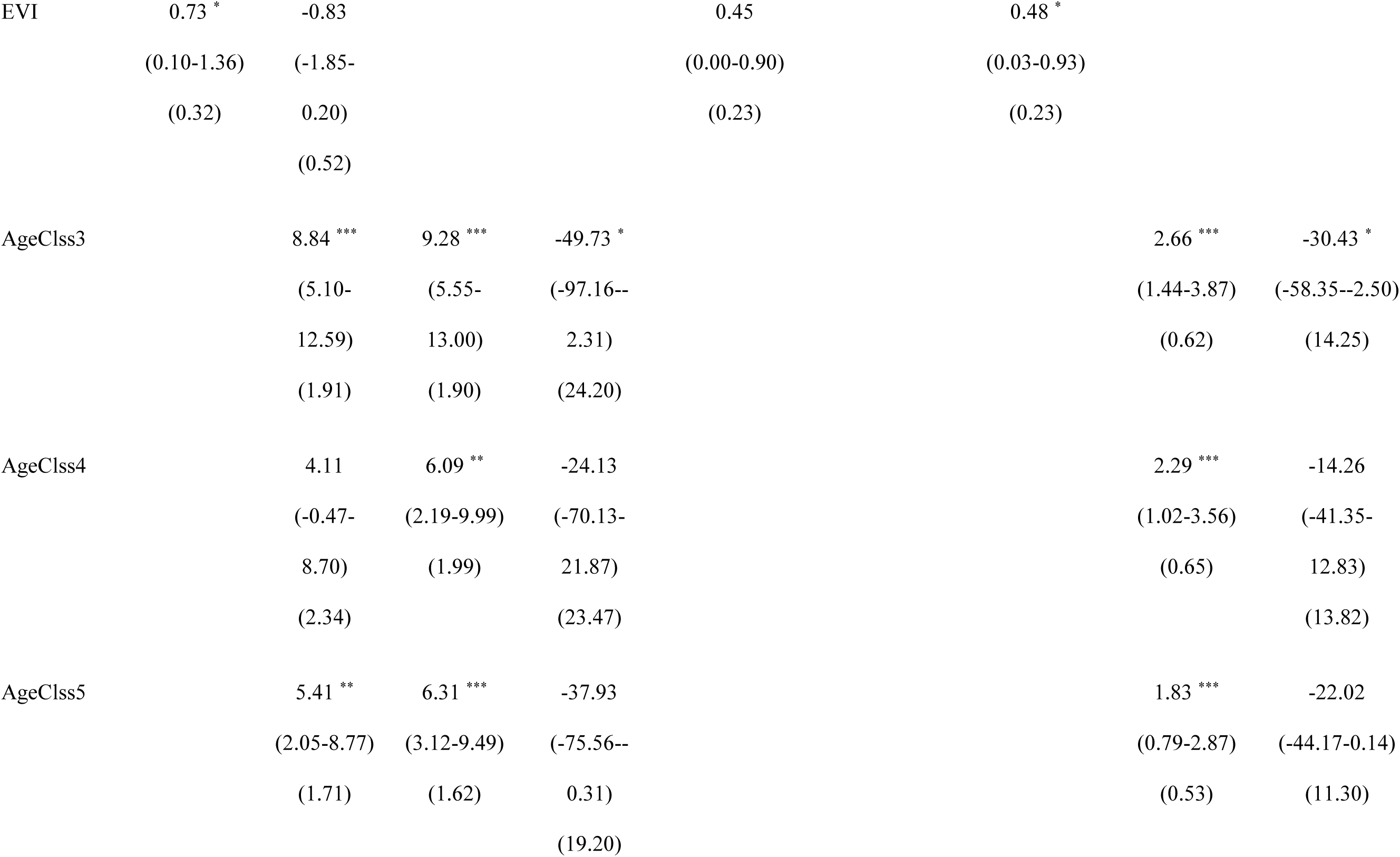
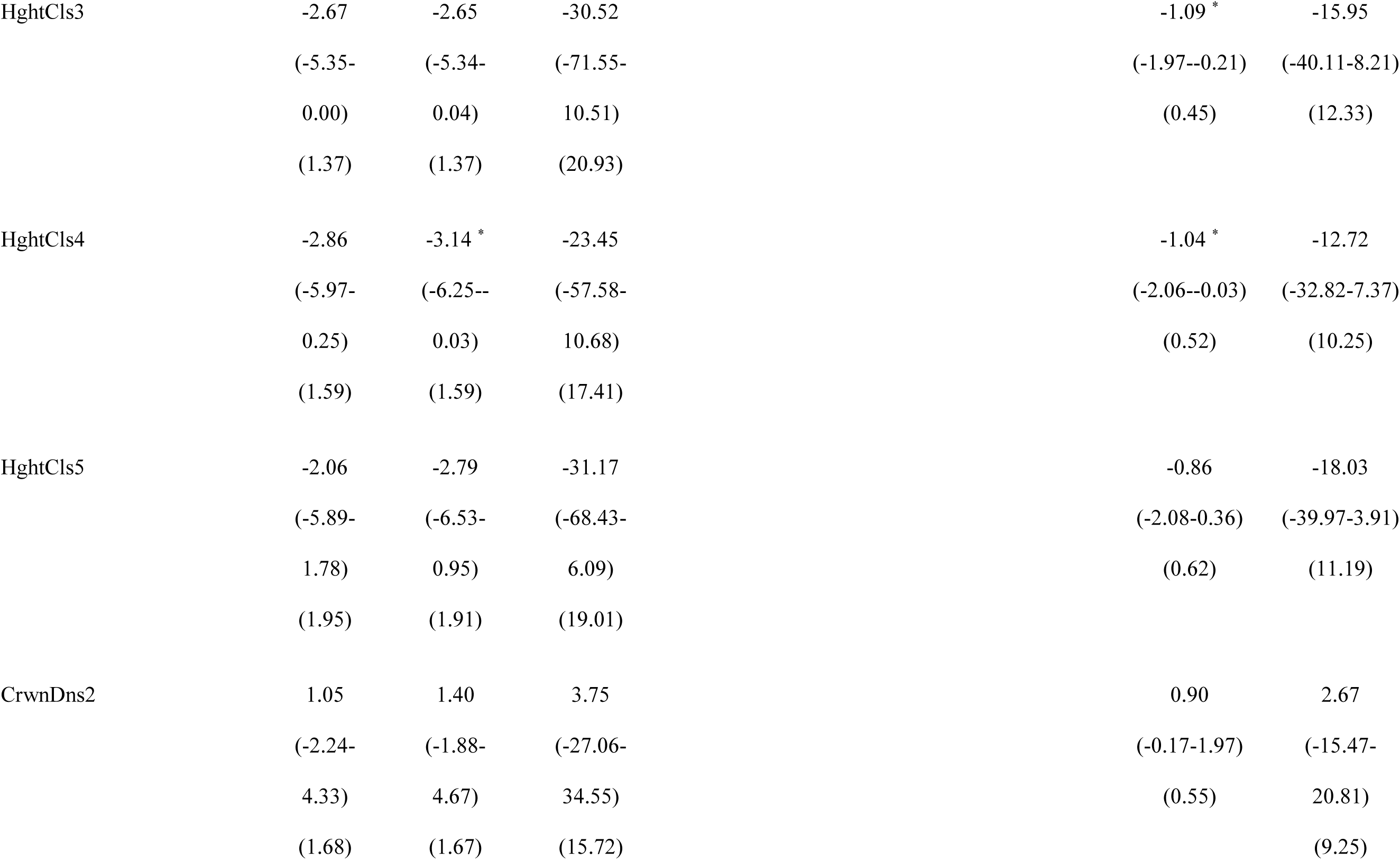
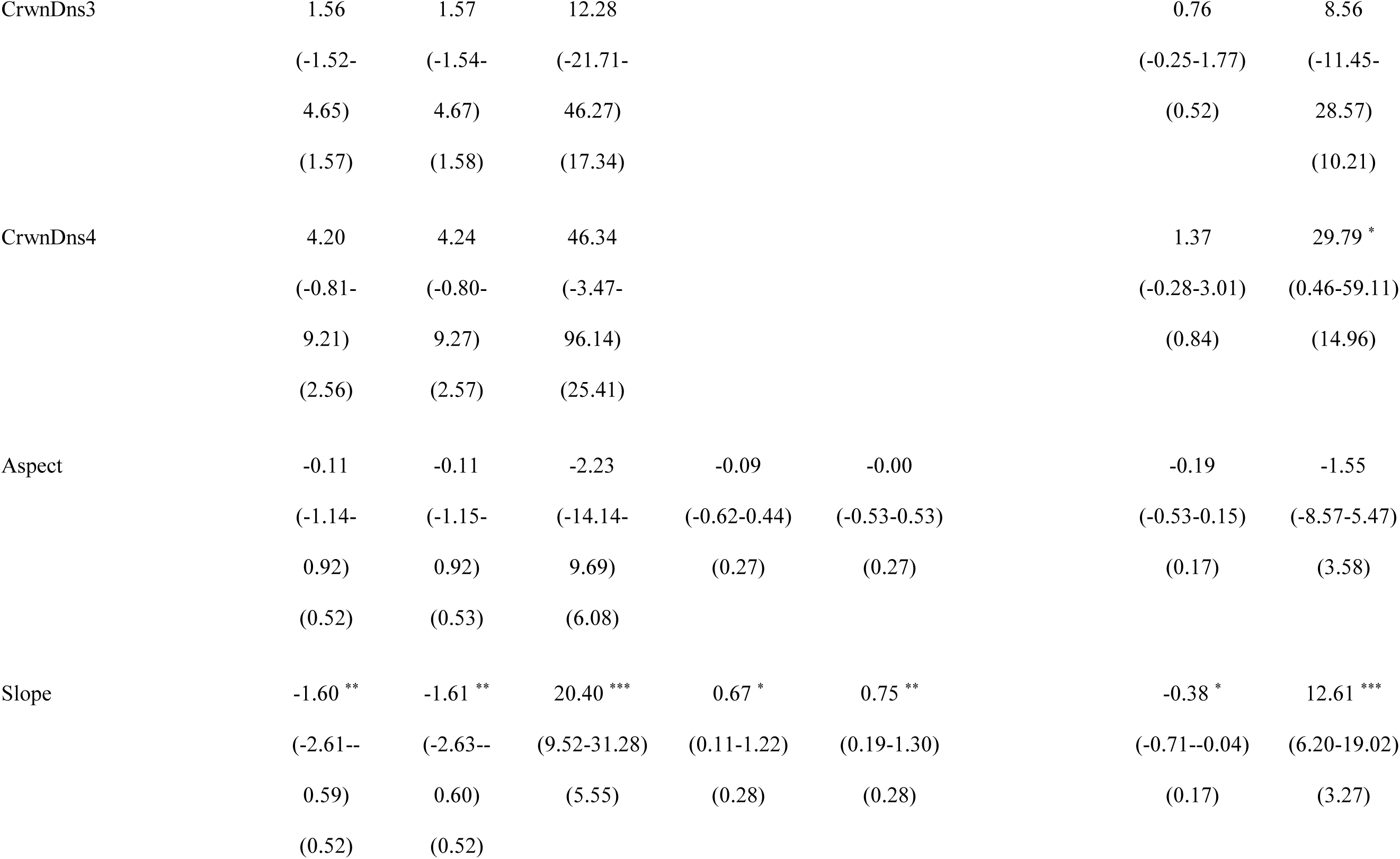
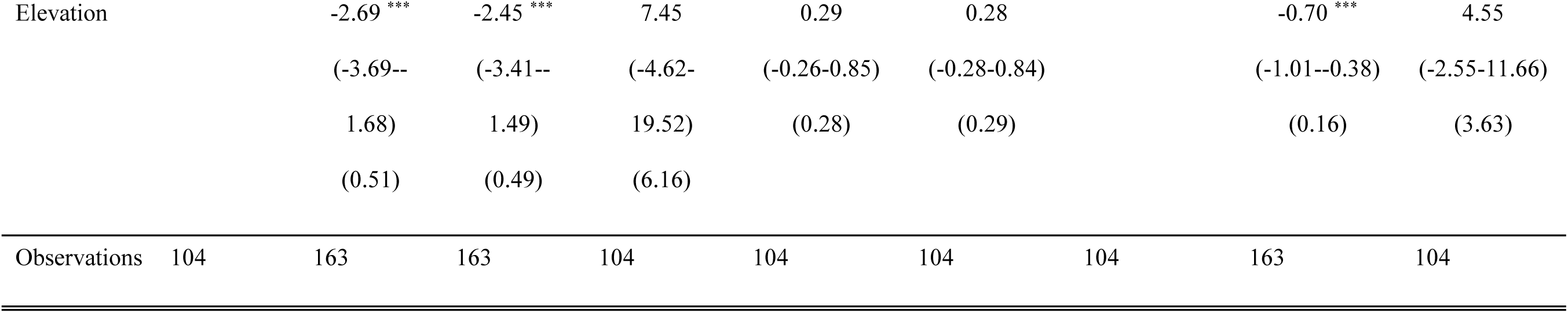
Part one of three for foliar phytochemical trait coefficient estimates, confidence intervals, and standard error values for top ranked models (< 2 ΔAIC_c_). Species codes are used for balsam fir (ABBA), red maple (ACRU), white birch (BEPA), black spruce (PIMA, and lowbush blueberry (VAAN). If there is more than one top ranked model per species, we present in order of ΔAIC_c_ rank. Model numbers are supplied beside the species code in the top row (see Table 1 for model descriptions). Models denoted with the suffix “r” and “b” represent raw and biomass basis respectively. Predictors include land cover (LandCover5 and LandCover6 represent deciduous and mix wood conditions), EVI (i.e., proxy for productivity), abiotic factors (aspect, slope, elevation), and biotic factors: AgeClss3 (41-60 years old), AgeClss 4 (61-80 years old), AgeClss5 (81-100 years old), HghtCls3 (6.6-9.5 m), HghtCls4 (9.6-12.5 m), HghtCls5 (12.6-15.5m), CrwnDns2 (51-75 % closed), CrwnDns3 (26-50%), CrwnDn4 (10-25 % closed). Total number of observations are provided in the bottom row. In addition, asterisks are used to indicate coefficient significance as follows: *\* p<0.05 ** p<0.01 *** p<0.001*.

**Table A13.**
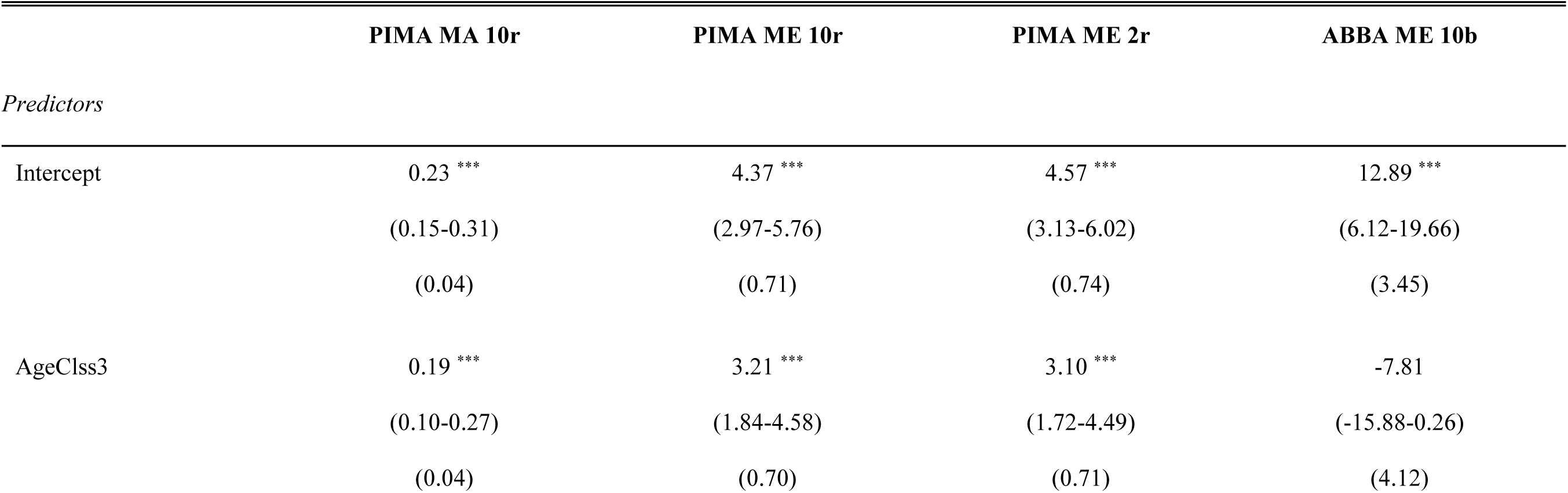
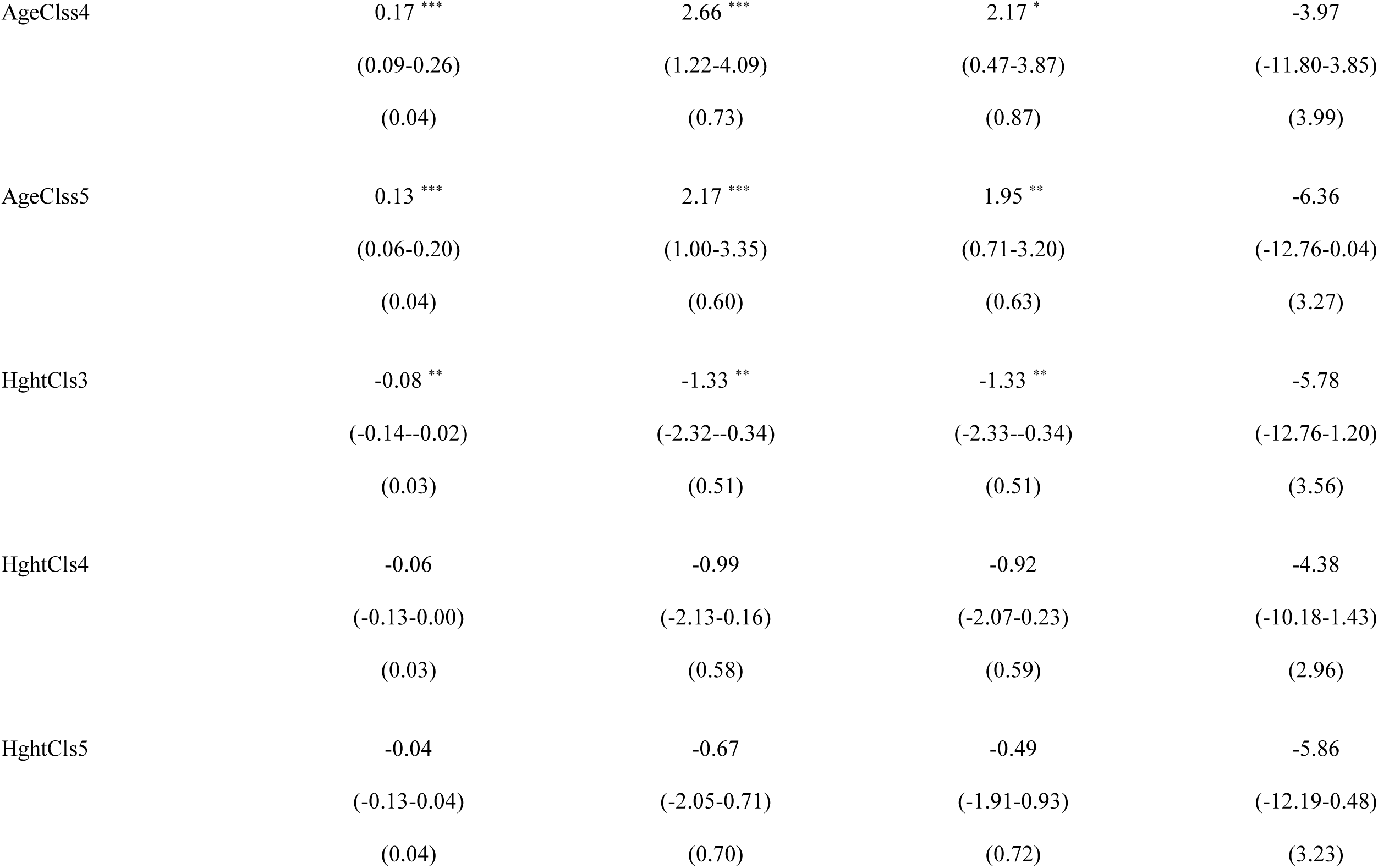
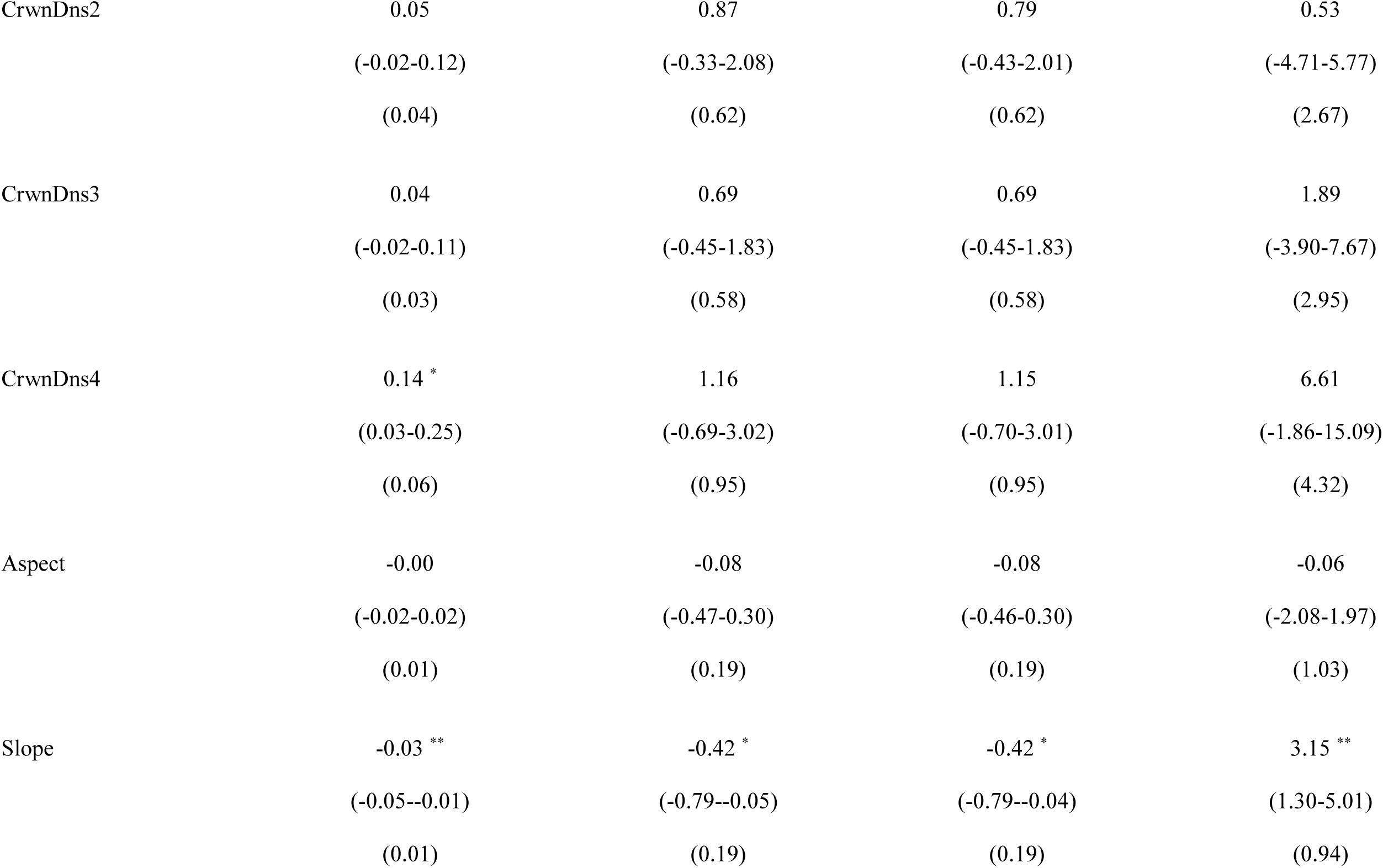
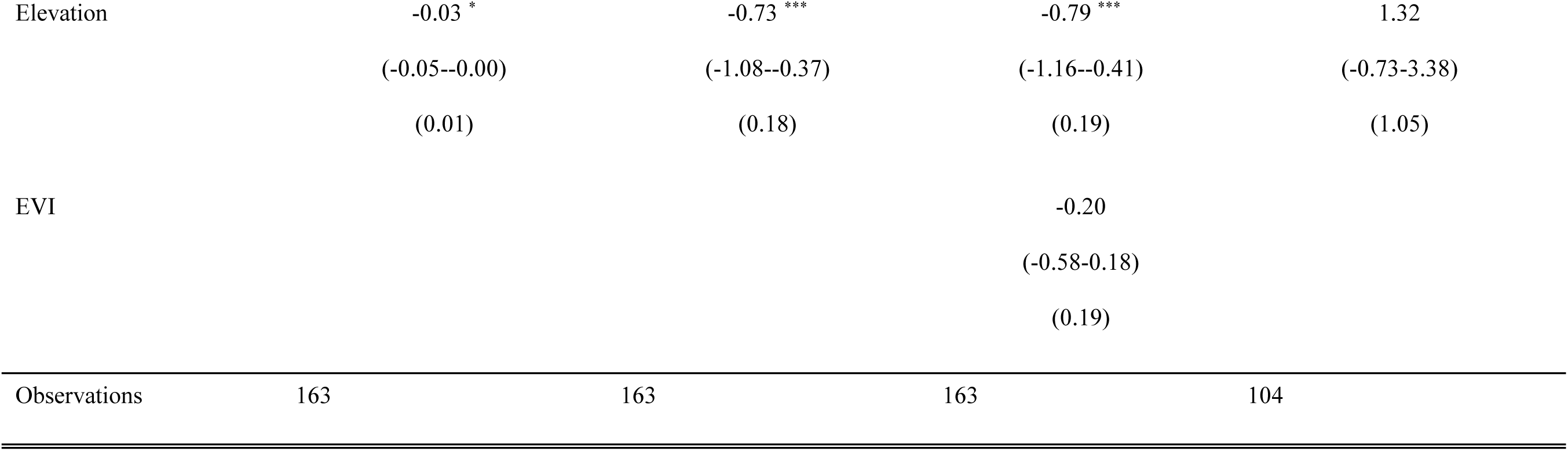
Part two of three for foliar phytochemical trait coefficient estimates, confidence intervals, and standard error values for top ranked models (< 2 ΔAIC_c_). Species codes are used for balsam fir (ABBA), red maple (ACRU), white birch (BEPA), black spruce (PIMA, and lowbush blueberry (VAAN). If there is more than one top ranked model per species, we present in order of ΔAIC_c_ rank. Model numbers are supplied beside the species code in the top row (see Table 1 for model descriptions). Codes are used for monoterpenic alcohol (MA) and monoterpenic ester (ME). Models denoted with the suffix “r” and “b” represent raw and biomass basis respectively. Predictors include land cover (LandCover5 and LandCover6 represent deciduous and mix wood conditions), EVI (i.e., proxy for productivity), abiotic factors (aspect, slope, elevation), and biotic factors: AgeClss3 (41-60 years old), AgeClss 4 (61-80 years old), AgeClss5 (81-100 years old), HghtCls3 (6.6-9.5 m), HghtCls4 (9.6-12.5 m), HghtCls5 (12.6-15.5m), CrwnDns2 (51-75 % closed), CrwnDns3 (26-50%), CrwnDn4 (10-25 % closed). Total number of observations are provided in the bottom row. In addition, asterisks are used to indicate coefficient significance as follows: *\* p<0.05 ** p<0.01 *** p<0.001*.

|  | PIMA MA 10r | PIMA ME 10r | PIMA ME 2r | ABBA ME 10b |
| --- | --- | --- | --- | --- |
| *Predictors* |  |  |  |  |
| Intercept | 0.23 ***<br>(0.15-0.31)<br>(0.04) | 4.37 ***<br>(2.97-5.76)<br>(0.71) | 4.57 ***<br>(3.13-6.02)<br>(0.74) | 12.89 ***<br>(6.12-19.66)<br>(3.45) |
| AgeCls3 | 0.19 ***<br>(0.10-0.27)<br>(0.04) | 3.21 ***<br>(1.84-4.58)<br>(0.70) | 3.10 ***<br>(1.72-4.49)<br>(0.71) | -7.81<br>(-15.88-0.26)<br>(4.12) |
| AgeClss4 | 0.17 ***<br>(0.09-0.26)<br>(0.04) | 2.66 ***<br>(1.22-4.09)<br>(0.73) | 2.17 *<br>(0.47-3.87)<br>(0.87) | -3.97<br>(-11.80-3.85)<br>(3.99) |
| AgeClss5 | 0.13 ***<br>(0.06-0.20)<br>(0.04) | 2.17 ***<br>(1.00-3.35)<br>(0.60) | 1.95 **<br>(0.71-3.20)<br>(0.63) | -6.36<br>(-12.76-0.04)<br>(3.27) |
| HghtCls3 | -0.08 **<br>(-0.14--0.02)<br>(0.03) | -1.33 **<br>(-2.32--0.34)<br>(0.51) | -1.33 **<br>(-2.33--0.34)<br>(0.51) | -5.78<br>(-12.76-1.20)<br>(3.56) |
| HghtCls4 | -0.06<br>(-0.13-0.00)<br>(0.03) | -0.99<br>(-2.13-0.16)<br>(0.58) | -0.92<br>(-2.07-0.23)<br>(0.59) | -4.38<br>(-10.18-1.43)<br>(2.96) |
| HghtCls5 | -0.04<br>(-0.13-0.04)<br>(0.04) | -0.67<br>(-2.05-0.71)<br>(0.70) | -0.49<br>(-1.91-0.93)<br>(0.72) | -5.86<br>(-12.19-0.48)<br>(3.23) |
| CrwnDns2 | 0.05<br>(-0.02-0.12)<br>(0.04) | 0.87<br>(-0.33-2.08)<br>(0.62) | 0.79<br>(-0.43-2.01)<br>(0.62) | 0.53<br>(-4.71-5.77)<br>(2.67) |
| CrwnDns3 | 0.04<br>(-0.02-0.11)<br>(0.03) | 0.69<br>(-0.45-1.83)<br>(0.58) | 0.69<br>(-0.45-1.83)<br>(0.58) | 1.89<br>(-3.90-7.67)<br>(2.95) |
| CrwnDns4 | 0.14 *<br>(0.03-0.25)<br>(0.06) | 1.16<br>(-0.69-3.02)<br>(0.95) | 1.15<br>(-0.70-3.01)<br>(0.95) | 6.61<br>(-1.86-15.09)<br>(4.32) |
| Aspect | -0.00<br>(-0.02-0.02)<br>(0.01) | -0.08<br>(-0.47-0.30)<br>(0.19) | -0.08<br>(-0.46-0.30)<br>(0.19) | -0.06<br>(-2.08-1.97)<br>(1.03) |
| Slope | -0.03 **<br>(-0.05--0.01)<br>(0.01) | -0.42 *<br>(-0.79--0.05)<br>(0.19) | -0.42 *<br>(-0.79--0.04)<br>(0.19) | 3.15 **<br>(1.30-5.01)<br>(0.94) |
| Elevation | -0.03 * | -0.73 *** | -0.79 *** | 1.32 |
|  | (-0.05--0.00) | (-1.08--0.37) | (-1.16--0.41) | (-0.73-3.38) |
|  | (0.01) | (0.18) | (0.19) | (1.05) |
| EVI |  |  | -0.20 |  |
|  |  |  | (-0.58-0.18) |  |
|  |  |  | (0.19) |  |
| Observations | 163 | 163 | 163 | 104 |

**Table A14.**
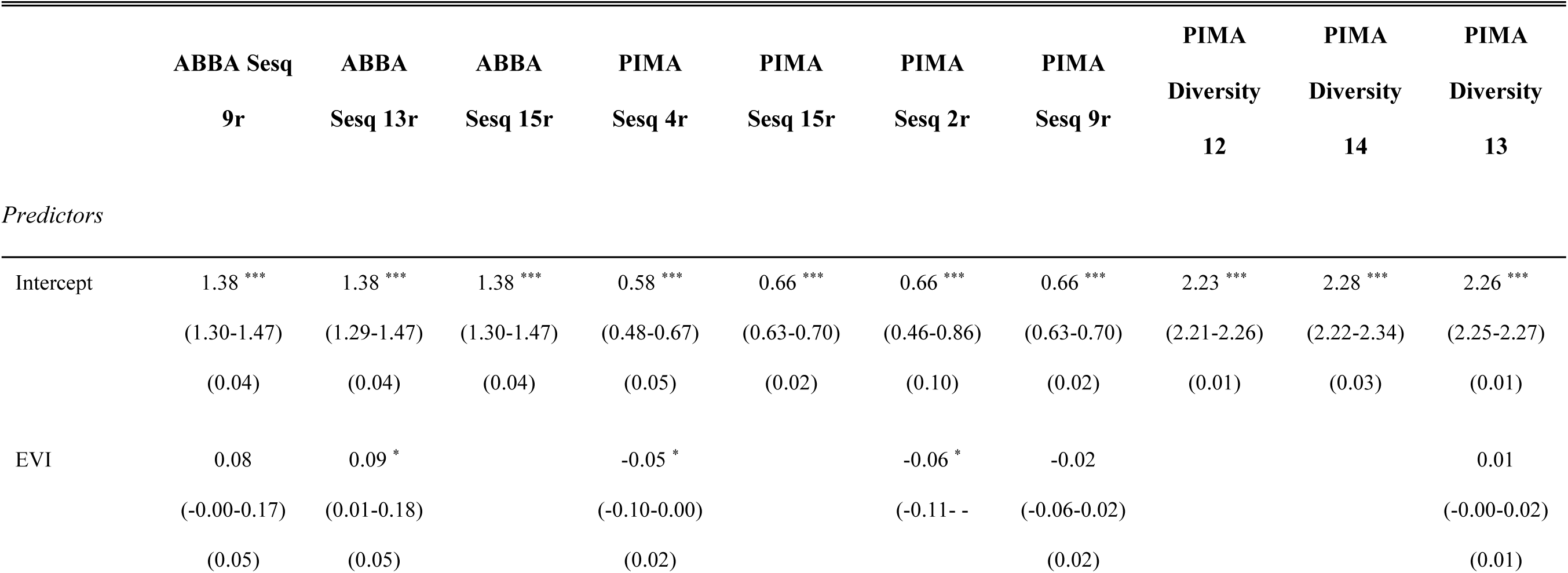
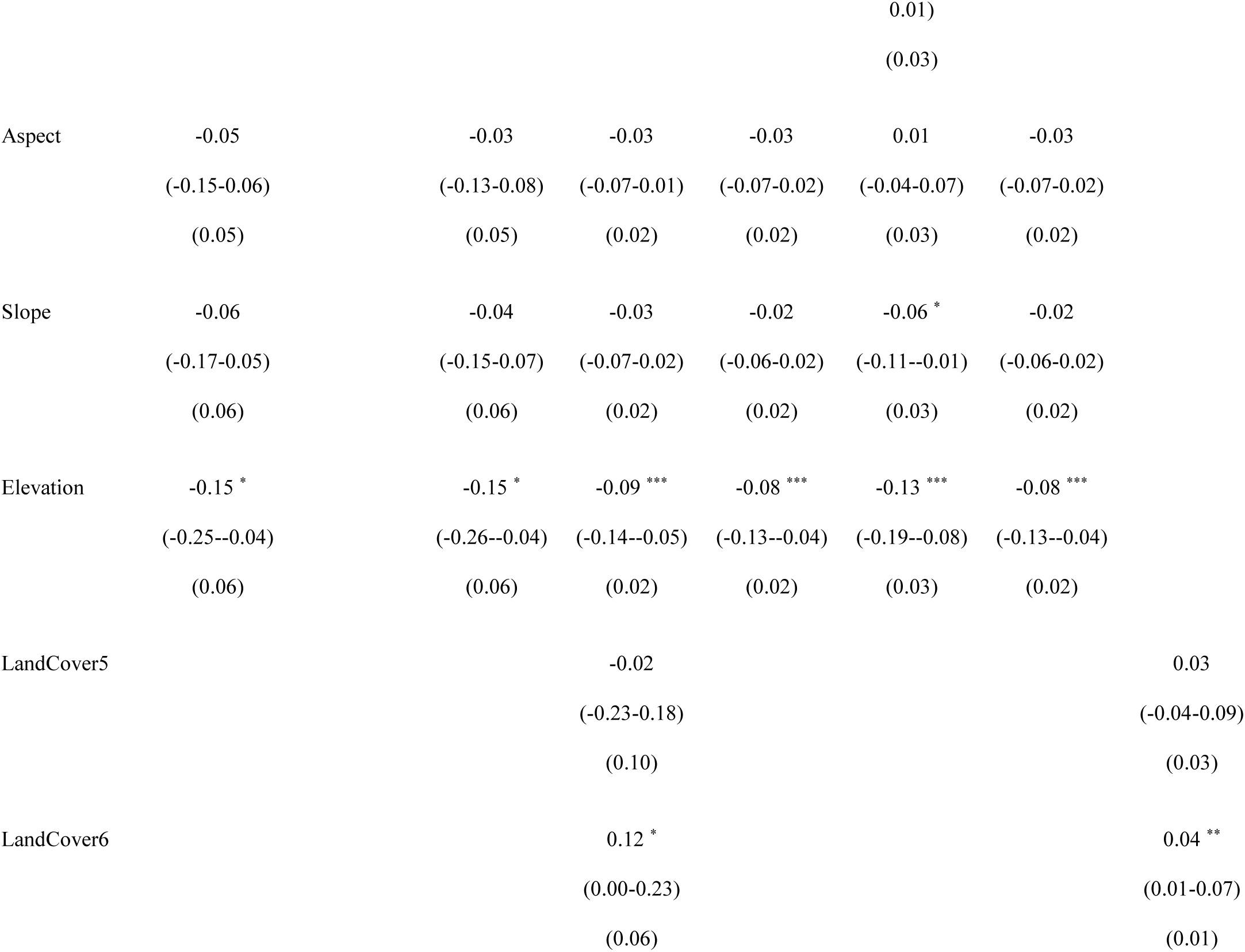
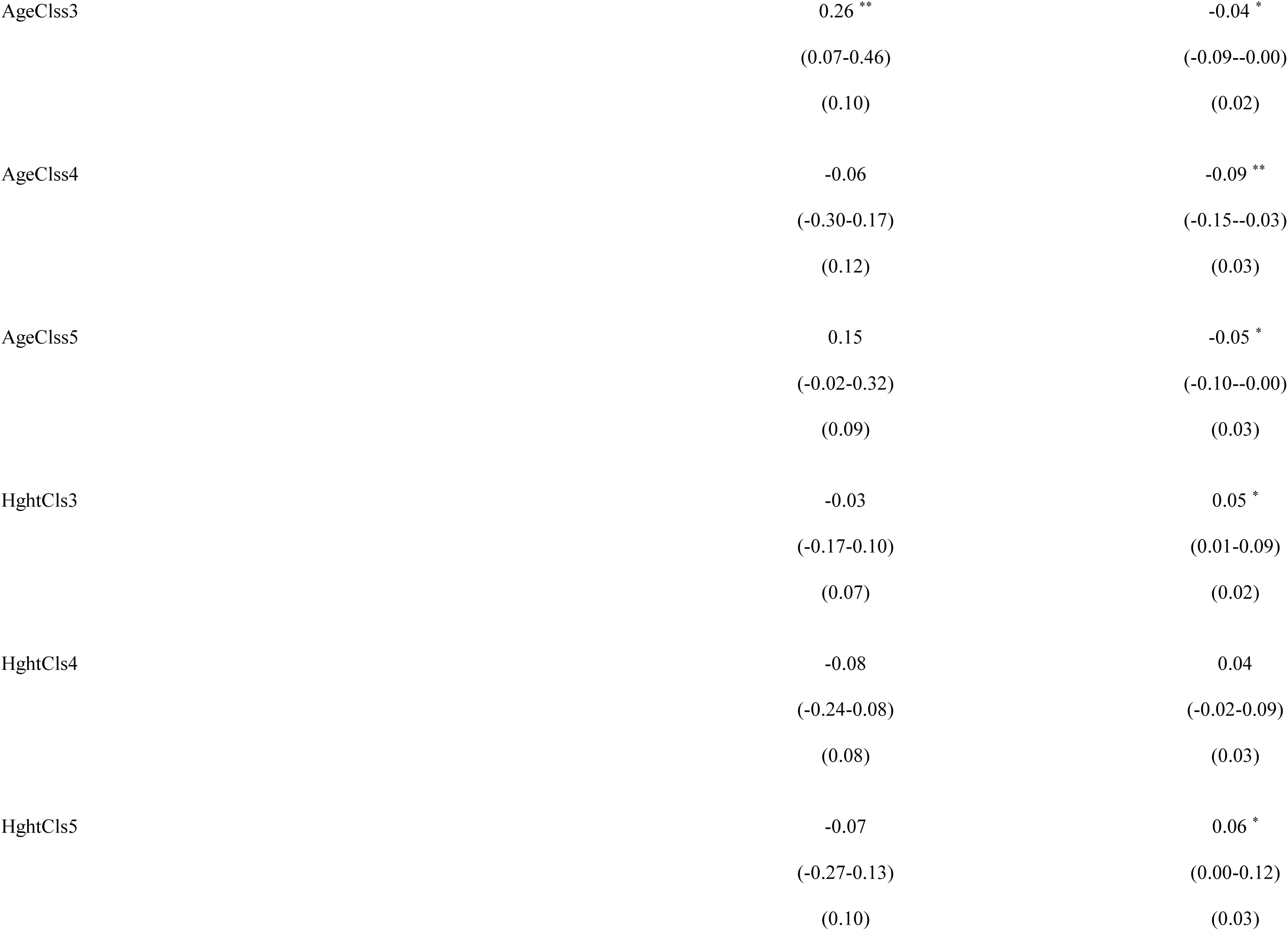
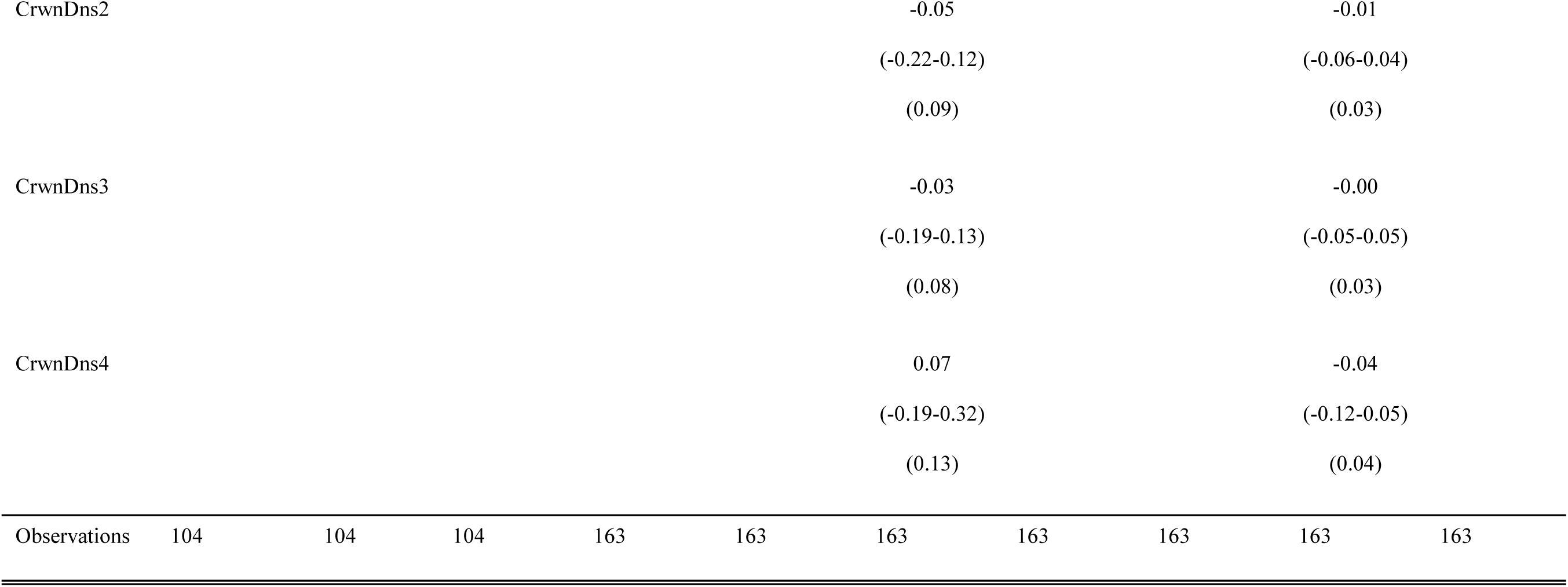
Part three of three for foliar phytochemical trait coefficient estimates, confidence intervals, and standard error values for top ranked models (< 2 ΔAIC_c_). Species codes are used for balsam fir (ABBA), red maple (ACRU), white birch (BEPA), black spruce (PIMA, and lowbush blueberry (VAAN). If there is more than one top ranked model per species, we present in order of ΔAIC_c_ rank. Model numbers are supplied beside the species code in the top row (see Table 1 for model descriptions). Sesquesterpene is truncated as sesq. Models denoted with the suffix “r” and “b” represent raw and biomass basis respectively. Predictors include land cover (LandCover5 and LandCover6 represent deciduous and mix wood conditions), EVI (i.e., proxy for productivity), abiotic factors (aspect, slope, elevation), and biotic factors: AgeClss3 (41-60 years old), AgeClss 4 (61-80 years old), AgeClss5 (81-100 years old), HghtCls3 (6.6-9.5 m), HghtCls4 (9.6-12.5 m), HghtCls5 (12.6-15.5m), CrwnDns2 (51-75 % closed), CrwnDns3 (26-50%), CrwnDn4 (10-25 % closed). Total number of observations are provided in the bottom row. In addition, asterisks are used to indicate coefficient significance as follows: *\* p<0.05 ** p<0.01 *** p<0.001*.

## Appendix 20

**Figure A6.**
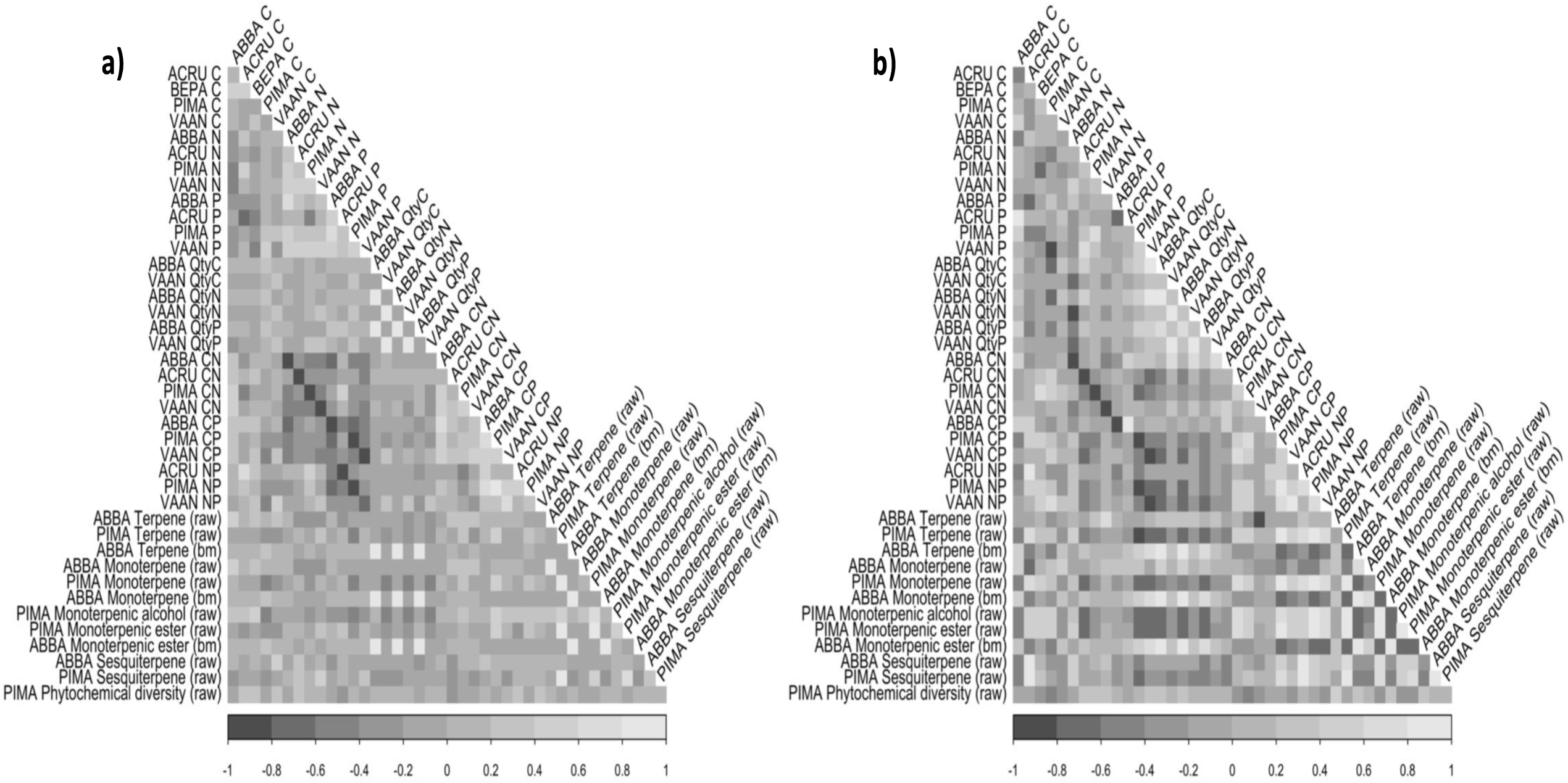
Correlation plot showing the relationships between our study species foliar elemental, stoichiometric, and phytochemical traits for top ranked models where the intercept was not with < 2 ΔAIC_c_. The left correlation plot (a) shows data space comparisons, for this we only compared plots in which all species were present (n = 29). The right correlation plot (b) shows spatial comparisons of predictive trait raster/surfaces. Correlation in data space is limited to co-occurrence of observations, while spatial correlation considers all pixels. In panel (b), we can see emergent patterns that are less apparent in data space comparisons (a).

## Notes

https://doi.org/10.6084/m9.figshare.11911455.v1

### Competing Interest Statement

The authors have declared no competing interest.

## Literature cited

Adler, P. B., Salguero-Gomez, R., Compagnoni, A., Hsu, J. S., Ray-Mukherjee, J., Mbeau-Ache, C., & Franco, M. (2014). Functional traits explain variation in plant life history strategies. Proceedings of the National Academy of Sciences, 111(2), 740–745. https://doi.org/10.1073/pnas.1315179111

Ågren, G. I. (1988). Ideal nutrient productivities and nutrient proportions in plant growth. Plant, Cell & Environment, 11(7), 613–620.

Alpert, P. (1991). Nitrogen sharing among ramets increases clonal growth in fragaria chiloensis. Ecology, 72(1), 69–80. https://doi.org/10.2307/1938903

Ashton, I. W., Miller, A. E., Bowman, W. D., & Suding, K. N. (2010). Niche complementarity due to plasticity in resource use: Plant partitioning of chemical N forms. Ecology, 91(11), 3252–3260. https://doi.org/10.1890/09-1849.1

Balluffi-Fry, J., Leroux, S. J., Wiersma, Y. F., Heckford, T. R., Rizzuto, M., Richmond, I. C., & Wal, E. V. (2020). Quantity–quality trade-offs revealed using a multiscale test of herbivore resource selection on elemental landscapes. Ecology and Evolution, *n/a*(n/a). https://doi.org/10.1002/ece3.6975

Balzotti, C. S., Asner, G. P., Taylor, P. G., Cleveland, C. C., Cole, R., Martin, R. E., Nasto, M., Osborne, B. B., Porder, S., & Townsend, A. R. (2016). Environmental controls on canopy foliar nitrogen distributions in a Neotropical lowland forest. Ecological Applications, 26(8), 2451–2464. https://doi.org/10.1002/eap.1408

Barron-Gafford, G. A., Scott, R. L., Jenerette, G. D., Hamerlynck, E. P., & Huxman, T. E. (2012). Temperature and precipitation controls over leaf- and ecosystem-level CO2 flux along a woody plant encroachment gradient. Global Change Biology, 18(4), 1389–1400. https://doi.org/10.1111/j.1365-2486.2011.02599.x

Becknell, J. M., & Powers, J. S. (2014). Stand age and soils as drivers of plant functional traits and aboveground biomass in secondary tropical dry forest. Canadian Journal of Forest Research. https://doi.org/10.1139/cjfr-2013-0331

Bernhardt, E. S., Blaszczak, J. R., Ficken, C. D., Fork, M. L., Kaiser, K. E., & Seybold, E. C. (2017). Control points in ecosystems: Moving beyond the hot spot hot moment concept. Ecosystems, 20(4), 665–682. https://doi.org/10.1007/s10021-016-0103-y

Bittebiere, A.-K., Saiz, H., & Mony, C. (2019). New insights from multidimensional trait space responses to competition in two clonal plant species. Functional Ecology, 297–307. https://doi.org/10.1111/1365-2435.13220

Blanes, M. C., Viñegla, B., Merino, J., & Carreira, J. A. (2013). Nutritional status of Abies pinsapo forests along a nitrogen deposition gradient: Do C/N/P stoichiometric shifts modify photosynthetic nutrient use efficiency? Oecologia, 171(4), 797–808. https://doi.org/10.1007/s00442-012-2454-1

Booker, F. L., & Maier, C. A. (2001). Atmospheric carbon dioxide, irrigation, and fertilization effects on phenolic and nitrogen concentrations in loblolly pine (Pinus taeda) needles. Tree Physiology, 21(9), 609–616. https://doi.org/10.1093/treephys/21.9.609

Brauer, V. S., Stomp, M., & Huisman, J. (2012). The nutrient-load hypothesis: Patterns of resource limitation and community structure driven by competition for nutrients and light. The American Naturalist, 179(6), 721–740. https://doi.org/10.1086/665650

Bryant, J. P., Chapin, F. S., & Klein, D. R. (1983). Carbon/nutrient balance of boreal plants in relation to vertebrate herbivory. Oikos, 40(3), 357. https://doi.org/10.2307/3544308

Burnham, K. P., & Anderson, D. R. (2002). Model selection and multimodel inference: A practical information-theoretic approach. Springer-Verlag.

Cachet, T., Brevard, H., Chaintreau, A., Demyttenaere, J., French, L., Gassenmeier, K., Joulain, D., Koenig, T., Leijs, H., Liddle, P., Loesing, G., Marchant, M., Merle, Ph., Saito, K., Schippa, C., Sekiya, F., & Smith, T. (2016). IOFI recommended practice for the use of predicted relative-response factors for the rapid quantification of volatile flavouring compounds by GC-FID. Flavour and Fragrance Journal, 31(3), 191–194. https://doi.org/10.1002/ffj.3311

Callis-Duehl, K., Vittoz, P., Defossez, E., & Rasmann, S. (2017). Community-level relaxation of plant defenses against herbivores at high elevation. Plant Ecology, 218(3), 291–304. https://doi.org/10.1007/s11258-016-0688-4

Canadian Digital Elevation Model: Product Specifications-Edition 1.1. (2016). [Map]. Natural Resources Canada.

Champagne, E., Moore, B. D., Côté, S. D., & Tremblay, J.-P. (2018). Spatial correlations between browsing on balsam fir by white-tailed deer and the nutritional value of neighboring winter forage. Ecology and Evolution, 8(5), 2812–2823. https://doi.org/10.1002/ece3.3878

Cornelissen, J. H. C., & Cornwell, W. K. (2014). The Tree of Life in ecosystems: Evolution of plant effects on carbon and nutrient cycling. Journal of Ecology, 102(2), 269–274. https://doi.org/10.1111/1365-2745.12217

Couture, J. J., Holeski, L. M., & Lindroth, R. L. (2014). Long-term exposure to elevated CO2 and O3 alters aspen foliar chemistry across developmental stages. Plant, Cell & Environment, 37(3), 758–765. https://doi.org/10.1111/pce.12195

Descombes, P., Marchon, J., Pradervand, J.-N., Bilat, J., Guisan, A., Rasmann, S., & Pellissier, L. (2017). Community-level plant palatability increases with elevation as insect herbivore abundance declines. Journal of Ecology, 105(1), 142–151. https://doi.org/10.1111/1365-2745.12664

Diaz, S., Hodgson, J. g., Thompson, K., Cabido, M., Cornelissen, J. h. c., Jalili, A., Montserrat-Martí, G., Grime, J. p., Zarrinkamar, F., Asri, Y., Band, S. r., Basconcelo, S., Castro-Díez, P., Funes, G., Hamzehee, B., Khoshnevi, M., Pérez-Harguindeguy, N., Pérez-Rontomé, M. c., Shirvany, F. a., … Zak, M. r. (2004). The plant traits that drive ecosystems: Evidence from three continents. Journal of Vegetation Science, 15(3), 295–304. https://doi.org/10.1111/j.1654-1103.2004.tb02266.x

Dirnböck, T., Kraus, D., Grote, R., Klatt, S., Kobler, J., Schindlbacher, A., Seidl, R., Thom, D., & Kiese, R. (2020). Substantial understory contribution to the C sink of a European temperate mountain forest landscape. Landscape Ecology, 35(2), 483–499. https://doi.org/10.1007/s10980-019-00960-2

Dunn, A. L., Wofsy, S. C., & Bright, A. V. H. (2009). Landscape heterogeneity, soil climate, and carbon exchange in a boreal black spruce forest. Ecological Applications, 19(2), 495–504. https://doi.org/10.1890/07-0771.1

Duparc, A., Garel, M., Marchand, P., Dubray, D., Maillard, D., & Loison, A. (2020). Through the taste buds of a large herbivore: Foodscape modeling contributes to an understanding of forage selection processes. Oikos, 129(2), 170–183. https://doi.org/10.1111/oik.06386

Ecological Stratification Working Group. (1996). A national ecological framework for Canada. Centre for Land and Biological Resources Research, Research Branch, Agriculture and Agri-Food Canada.

Elser, J. J., Fagan, W. F., Kerkhoff, A. J., Swenson, N. G., & Enquist, B. J. (2010). Biological stoichiometry of plant production: Metabolism, scaling and ecological response to global change: Tansley review. New Phytologist, 186(3), 593–608. https://doi.org/10.1111/j.1469-8137.2010.03214.x

Elser, James J, & Hamilton, A. (2007). Stoichiometry and the new biology: The future is now. PLoS Biology, 5(7), e181. https://doi.org/10.1371/journal.pbio.0050181

Esri (10.8). (2020). [Computer software]. https://www.esri.com/en-us/arcgis/products/arcgis-pro/

Evans, J. (2020). spatialEco. R package (1.3–3) [Computer software]. https://github.com/jeffreyevans/spatialEco

Fajardo, A., & Siefert, A. (2018). Intraspecific trait variation and the leaf economics spectrum across resource gradients and levels of organization. Ecology, 99(5), 1024–1030. https://doi.org/10.1002/ecy.2194

Fan, H., Wu, J., Liu, W., Yuan, Y., Hu, L., & Cai, Q. (2015). Linkages of plant and soil C:N:P stoichiometry and their relationships to forest growth in subtropical plantations. Plant and Soil, 392(1–2), 127–138. https://doi.org/10.1007/s11104-015-2444-2

Filipiak, M. (2018). A better understanding of bee nutritional ecology is needed to optimize conservation strategies for wild bees—The application of ecological stoichiometry. Insects, 9(3). https://doi.org/10.3390/insects9030085

Forkner, R. E., & Marquis, R. J. (2004). Uneven-aged and even-aged logging alter foliar phenolics of oak trees remaining in forested habitat matrix. Forest Ecology and Management, 199(1), 21–37. https://doi.org/10.1016/j.foreco.2004.03.044

Fyllas, N. M., Michelaki, C., Galanidis, A., Evangelou, E., Zaragoza-Castells, J., Dimitrakopoulos, P. G., Tsadilas, C., Arianoutsou, M., & Lloyd, J. (2020). Functional trait variation among and within species and plant functional types in mountainous mediterranean forests. Frontiers in Plant Science, 11. https://doi.org/10.3389/fpls.2020.00212

Gartner, T. B., & Cardon, Z. G. (2004). Decomposition dynamics in mixed-species leaf litter. Oikos, 104(2), 230–246. JSTOR.

Gimona, A., & van der Horst, D. (2007). Mapping hotspots of multiple landscape functions: A case study on farmland afforestation in Scotland. Landscape Ecology, 22(8), 1255–1264. https://doi.org/10.1007/s10980-007-9105-7

Glassmire, A. E., Jeffrey, C. S., Forister, M. L., Parchman, T. L., Nice, C. C., Jahner, J. P., Wilson, J. S., Walla, T. R., Richards, L. A., Smilanich, A. M., Leonard, M. D., Morrison, C. R., Simbaña, W., Salagaje, L. A., Dodson, C. D., Miller, J. S., Tepe, E. J., Villamarin-Cortez, S., & Dyer, L. A. (2016). Intraspecific phytochemical variation shapes community and population structure for specialist caterpillars. New Phytologist, 212(1), 208–219. https://doi.org/10.1111/nph.14038

Gosse, J., Hermanutz, L., McLaren, B., Deering, P., & Knight, T. (2011). Degradation of boreal forests by nonnative herbivores in Newfoundland’s National Parks: Recommendations for ecosystem restoration. Natural Areas Journal, 31(4), 331–339. https://doi.org/10.3375/043.031.0403

Grime, P. J., & Pierce, S. (2012). The evolutionary strategies that shape ecosystems. Wiley-Blackwell.

Hallett, R. A., & Hornbeck, J. W. (1997). Foliar and soil nutrient relationships in red oak and white pine forests. Canadian Journal of Forest Research, 27, 12.

Harpole, W. S., Ngai, J. T., Cleland, E. E., Seabloom, E. W., Borer, E. T., Bracken, M. E. S., Elser, J. J., Gruner, D. S., Hillebrand, H., Shurin, J. B., & Smith, J. E. (2011). Nutrient co-limitation of primary producer communities. Ecology Letters, 14(9), 852–862. https://doi.org/10.1111/j.1461-0248.2011.01651.x

Harvey, E., Gounand, I., Fronhofer, E. A., & Altermatt, F. (2019). Metaecosystem dynamics drive community composition in experimental, multi-layered spatial networks. Oikos. https://doi.org/10.1111/oik.07037

Harvey, E., Gounand, I., Ward, C. L., & Altermatt, F. (2017). Bridging ecology and conservation: From ecological networks to ecosystem function. Journal of Applied Ecology, 54(2), 371–379. https://doi.org/10.1111/1365-2664.12769

Hassell, M. P., Comins, H. N., & May, R. M. (1994). Species coexistence and self-organizing spatial dynamics. Nature, 370, 290–292.

Haynes, K. J., & Cronin, J. T. (2004). Confounding of patch quality and matrix effects in herbivore movement studies. Landscape Ecology, 19(2), 119–124. https://doi.org/10.1023/B:LAND.0000021721.41349.85

He, P., Fontana, S., Sardans, J., Peñuelas, J., Gessler, A., Schaub, M., Rigling, A., Li, H., Jiang, Y., & Li, M.-H. (2019). The biogeochemical niche shifts of Pinus sylvestris var. Mongolica along an environmental gradient. Environmental and Experimental Botany, 167, 103825. https://doi.org/10.1016/j.envexpbot.2019.103825

Hemming, J. D. C., & Lindroth, R. L. (1999). Effects of light and nutrient availability on aspen: Growth, phytochemistry, and insect performance. Journal of Chemical Ecology, 25(7), 1687–1714. https://doi.org/10.1023/A:1020805420160

Hessen, D. O., Ågren, G. I., Anderson, T. R., Elser, J. J., & de Ruiter, P. C. (2004). Carbon sesquestration in ecosystems: The role of stoichiometry. Ecology, 85(5), 1179–1192. https://doi.org/10.1890/02-0251

Hijmans, R., J. (2020). Raster: Geographic Data Analysis and Modeling. R package (3.4–5) [Computer software].

Hobbie, S. E. (2015). Plant species effects on nutrient cycling: Revisiting litter feedbacks. Trends in Ecology & Evolution, 30(6), 357–363. https://doi.org/10.1016/j.tree.2015.03.015

Hunter, M. D. (2016). The Phytochemical Landscape: Linking Trophic Interactions and Nutrient Dynamics. Princeton University Press.

Hunter, M. D., & Schultz, J. C. (1995). Fertilization mitigates chemical induction and herbivore responses within damaged oak trees. Ecology, 76(4), 1226–1232. https://doi.org/10.2307/1940929

Hussain, A., Rodriguez-Ramos, J. C., & Erbilgin, N. (2019). Spatial characteristics of volatile communication in lodgepole pine trees: Evidence of kin recognition and intra-species support. Science of The Total Environment, 692, 127–135. https://doi.org/10.1016/j.scitotenv.2019.07.211

Jiang, Yong, Zang, R., Lu, X., Huang, Y., Ding, Y., Liu, W., Long, W., Zhang, J., & Zhang, Z. (2015). Effects of soil and microclimatic conditions on the community-level plant functional traits across different tropical forest types. Plant and Soil, 390(1), 351–367. https://doi.org/10.1007/s11104-015-2411-y

Jiang, Yueyang, Rocha, A. V., Rastetter, E. B., Shaver, G. R., Mishra, U., Zhuang, Q., & Kwiatkowski, B. L. (2016). C–N–P interactions control climate driven changes in regional patterns of C storage on the North Slope of Alaska. Landscape Ecology, 31(1), 195–213. https://doi.org/10.1007/s10980-015-0266-5

Jobbágy, E. G., & Jackson, R. B. (2004). The uplift of soil nutrients by plants: Biogeochemical consequences across scales. Ecology, 85(9), 2380–2389. https://doi.org/10.1890/03-0245

Jones, J. W., Starbuck, M. J., & Jenkerson, C. B. (2013). Landsat surface reflectance quality assurance extraction (version 1.7): U.S. Geological Survey Techniques and Methods. https://pubs.usgs.gov/tm/11/c07

Jung, V., Violle, C., Mondy, C., Hoffmann, L., & Muller, S. (2010). Intraspecific variability and trait-based community assembly. Journal of Ecology, 98(5), 1134–1140. https://doi.org/10.1111/j.1365-2745.2010.01687.x

Kerkhoff, A. J., Enquist, B. J., Elser, J. J., & Fagan, W. F. (2005). Plant allometry, stoichiometry and the temperature-dependence of primary productivity. Global Ecology and Biogeography, 14(6), 585–598. https://doi.org/10.1111/j.1466-822X.2005.00187.x

Kessler, A. (2015). The information landscape of plant constitutive and induced secondary metabolite production. Current Opinion in Insect Science, 8, 47–53. https://doi.org/10.1016/j.cois.2015.02.002

Kichenin, E., Wardle, D. A., Peltzer, D. A., Morse, C. W., & Freschet, G. T. (2013). Contrasting effects of plant inter- and intraspecific variation on community-level trait measures along an environmental gradient. Functional Ecology, 27(5), 1254–1261. https://doi.org/10.1111/1365-2435.12116

Knops, J. M. H., Bradley, K. L., & Wedin, D. A. (2002). Mechanisms of plant species impacts on ecosystem nitrogen cycling. Ecology Letters, 5(3), 454–466. https://doi.org/10.1046/j.1461-0248.2002.00332.x

Krishna, M. P., & Mohan, M. (2017). Litter decomposition in forest ecosystems: A review. Energy, Ecology and Environment, 2(4), 236–249. https://doi.org/10.1007/s40974-017-0064-9

Land cover map of North American at 30 m resolution (1st ed.). (2017). [Map]. Commission for Environmental Cooperation.

Lavorel, S., Grigulis, K., Lamarque, P., Colace, M.-P., Garden, D., Girel, J., Pellet, G., & Douzet, R. (2011). Using plant functional traits to understand the landscape distribution of multiple ecosystem services. Journal of Ecology, 99(1), 135–147. https://doi.org/10.1111/j.1365-2745.2010.01753.x

Leroux, S. J. (2019). On the prevalence of uninformative parameters in statistical models applying model selection in applied ecology. PLoS ONE, 14(2), 12. https://doi.org/10.1371/journal.pone.0206711

Leroux, S. J., Wal, E. V., Wiersma, Y. F., Charron, L., Ebel, J. D., Ellis, N. M., Hart, C., Kissler, E., Saunders, P. W., Moudrá, L., Tanner, A. L., & Yalcin, S. (2017). Stoichiometric distribution models: Ecological stoichiometry at the landscape extent. Ecology Letters, 20(12), 1495–1506. https://doi.org/10.1111/ele.12859

Li, B., Shibuya, T., Yogo, Y., & Hara, T. (2004). Effects of ramet clipping and nutrient availability on growth and biomass allocation of yellow nutsedge. Ecological Research, 19(6), 603–612. https://doi.org/10.1111/j.1440-1703.2004.00685.x

Lindroth, R. L., Osier, T. L., Barnhill, H. R. H., & Wood, S. A. (2002). Effects of genotype and nutrient availability on phytochemistry of trembling aspen (Populus tremuloides Michx.) during leaf senescence. Biochemical Systematics and Ecology, 30(4), 297–307. https://doi.org/10.1016/S0305-1978(01)00088-6

Liu, S., Yan, Z., Chen, Y., Zhang, M., Chen, J., & Han, W. (2019). Foliar pH, an emerging plant functional trait: Biogeography and variability across northern China. Global Ecology and Biogeography, 28(3), 386–397. https://doi.org/10.1111/geb.12860

Lohbeck, M., Poorter, L., Lebrija-Trejos, E., Martínez-Ramos, M., Meave, J. A., Paz, H., Pérez-García, E. A., Romero-Pérez, I. E., Tauro, A., & Bongers, F. (2013). Successional changes in functional composition contrast for dry and wet tropical forest. Ecology, 94(6), 1211–1216. https://doi.org/10.1890/12-1850.1

Lovell, S. T., & Johnston, D. M. (2009). Designing landscapes for performance based on emerging principles in landscape ecology. Ecology and Society, 14(1). https://www.jstor.org/stable/26268059

Macek, M., Kopecký, M., & Wild, J. (2019). Maximum air temperature controlled by landscape topography affects plant species composition in temperate forests. Landscape Ecology, 34(11), 2541–2556. https://doi.org/10.1007/s10980-019-00903-x

McClain, M. E., Boyer, E. W., Dent, C. L., Gergel, S. E., Grimm, N. B., Groffman, P. M., Hart, S. C., Harvey, J. W., Johnston, C. A., Mayorga, E., McDowell, W. H., & Pinay, G. (2003). Biogeochemical hot spots and hot moments at the interface of terrestrial and aquatic ecosystems. Ecosystems, 6(4), 301–312. https://doi.org/10.1007/s10021-003-0161-9

Mendez, M., & Karlsson, S. P. (2005). Nutrient stoichiometry in pinguicula vulgaris: Nutrient availability, plant size, and reproductive status. Ecology, 86(4), 982–991.

Morquecho-Contreras, A., Zepeda-Gómez, C., & Sánchez-Sánchez, H. (2018). Plant antiherbivore defense in diverse environments. In L. Hufnagel (Ed.), Pure and Applied Biogeography. InTech. https://doi.org/10.5772/intechopen.70418

Müller, M., Oelmann, Y., Schickhoff, U., Böhner, J., & Scholten, T. (2017). Himalayan treeline soil and foliar C:N:P stoichiometry indicate nutrient shortage with elevation. Geoderma, 291, 21–32. https://doi.org/10.1016/j.geoderma.2016.12.015

Muraoka, H., Noda, H. M., Nagai, S., Motohka, T., Saitoh, T. M., Nasahara, K. N., & Saigusa, N. (2013). Spectral vegetation indices as the indicator of canopy photosynthetic productivity in a deciduous broadleaf forest. Journal of Plant Ecology, 6(5), 393–407. https://doi.org/10.1093/jpe/rts037

Niinemets, Ü., & Kull, O. (1998). Stoichiometry of foliar carbon constituents varies along light gradients in temperate woody canopies: Implications for foliage morphological plasticity. Tree Physiology, 18(7), 467–479. https://doi.org/10.1093/treephys/18.7.467

Pan, Y., Horn, J., Jenkins, J., & Birdsey, R. (2004). Importance of foliar nitrogen concentration to predict forest productivity in the mid-Atlantic region. Forest Science, 50(3), 11.

Pausas, J. G., & Bond, W. J. (2019). Humboldt and the reinvention of nature. Journal of Ecology, 107(3), 1031–1037. https://doi.org/10.1111/1365-2745.13109

Pellissier, L., Moreira, X., Danner, H., Serrano, M., Salamin, N., van Dam, N. M., & Rasmann, S. (2016). The simultaneous inducibility of phytochemicals related to plant direct and indirect defences against herbivores is stronger at low elevation. Journal of Ecology, 104(4), 1116–1125. https://doi.org/10.1111/1365-2745.12580

Philben, M., Ziegler, S. E., Edwards, K. A., Kahler, R., & Benner, R. (2016). Soil organic nitrogen cycling increases with temperature and precipitation along a boreal forest latitudinal transect. Biogeochemistry, 127(2–3), 397–410. https://doi.org/10.1007/s10533-016-0187-7

Pickett, S. T. A., & Cadenasso, M. L. (1995). Landscape ecology: Spatial heterogeneity in ecological systems. Science, 269(5222), 331–334. https://doi.org/10.1126/science.269.5222.331

Poitevin, E. (2016). Official methods for the determination of minerals and trace elements in infant formula and milk products: A review. Journal of AOAC International, 99(1), 42–52. https://doi.org/10.5740/jaoacint.15-0246

Ponette-González, A. G., Weathers, K. C., & Curran, L. M. (2010). Tropical land-cover change alters biogeochemical inputs to ecosystems in a Mexican montane landscape. Ecological Applications, 20(7), 1820–1837. https://doi.org/10.1890/09-1125.1

Poorter, L., & Bongers, F. (2006). Leaf traits are good predictors of plant performance across 53 rain forest species. Ecology, 87(7), 1733–1743. https://doi.org/10.1890/0012-9658(2006)87[1733:LTAGPO]2.0.CO;2

Pyakurel, A., & Wang, J. R. (2014). Leaf morphological and stomatal variations in paper birch populations along environmental gradients in Canada. American Journal of Plant Sciences, 05(11), 1508–1520. https://doi.org/10.4236/ajps.2014.511166

R Core Team. (2020). R: a language and environment for statistical computing. R Foundation for Statistical Computing. https://www.R-project.org/

Radwan, M. A., & Harrington, C. A. (2011). Foliar chemical concentrations, growth, and site productivity relations in western red cedar. Canadian Journal of Forest Research. https://doi.org/10.1139/x86-185

Reich, P. B., & Oleksyn, J. (2004). Global patterns of plant leaf N and P in relation to temperature and latitude. Proceedings of the National Academy of Sciences, 101(30), 11001–11006. https://doi.org/10.1073/pnas.0403588101

Requena-Mullor, J. M., López, E., Castro, A. J., Alcaraz-Segura, D., Castro, H., Reyes, A., & Cabello, J. (2017). Remote-sensing based approach to forecast habitat quality under climate change scenarios. PLOS ONE, 12(3), e0172107. https://doi.org/10.1371/journal.pone.0172107

Richards, L. A., Dyer, L. A., Forister, M. L., Smilanich, A. M., Dodson, C. D., Leonard, M. D., & Jeffrey, C. S. (2015). Phytochemical diversity drives plant–insect community diversity. Proceedings of the National Academy of Sciences of the United States of America, 112(35), 10973–10978. https://doi.org/10.1073/pnas.1504977112

Richardson, A. D. (2004). Foliar chemistry of balsam fir and red spruce in relation to elevation and the canopy light gradient in the mountains of the northeastern United States. Plant and Soil, 260(1), 291–299. https://doi.org/10.1023/B:PLSO.0000030179.02819.85

Rijkers, T., Pons, T. L., & Bongers, F. (2000). The effect of tree height and light availability on photosynthetic leaf traits of four neotropical species differing in shade tolerance. Functional Ecology, 14(1), 77–86. https://doi.org/10.1046/j.1365-2435.2000.00395.x

Rizzuto, M., Leroux, S. J., Wal, E. V., Wiersma, Y. F., Heckford, T. R., & Balluffi-Fry, J. (2019). Patterns and potential drivers of intraspecific variability in the body C, N, and P composition of a terrestrial consumer, the snowshoe hare (Lepus americanus). Ecology and Evolution, 9(24), 14453–14464. https://doi.org/10.1002/ece3.5880

Santiago, L. S., Kitajima, K., Wright, S. J., & Mulkey, S. S. (2004). Coordinated changes in photosynthesis, water relations and leaf nutritional traits of canopy trees along a precipitation gradient in lowland tropical forest. Oecologia, 139(4), 495–502. https://doi.org/10.1007/s00442-004-1542-2

Sardans, J., Alonso, R., Carnicer, J., Fernández-Martínez, M., Vivanco, M. G., & Peñuelas, J. (2016). Factors influencing the foliar elemental composition and stoichiometry in forest trees in Spain. Perspectives in Plant Ecology, Evolution and Systematics, 18, 52–69. https://doi.org/10.1016/j.ppees.2016.01.001

Sardans, Jordi, Alonso, R., Janssens, I. A., Carnicer, J., Vereseglou, S., Rillig, M. C., Fernández-Martínez, M., Sanders, T. G. M., & Peñuelas, J. (2016). Foliar and soil concentrations and stoichiometry of nitrogen and phosphorous across European *Pinus sylvestris* forests: Relationships with climate, N deposition and tree growth. Functional Ecology, 30(5), 676–689. https://doi.org/10.1111/1365-2435.12541

Schmitz, O. J., Wilmers, C. C., Leroux, S. J., Doughty, C. E., Atwood, T. B., Galetti, M., Davies, A. B., & Goetz, S. J. (2018). Animals and the zoogeochemistry of the carbon cycle. Science, 362(6419), eaar3213. https://doi.org/10.1126/science.aar3213

Sedio, B. E., Echeverri, J. C. R., P, C. A. B., & Wright, S. J. (2017). Sources of variation in foliar secondary chemistry in a tropical forest tree community. Ecology, 98(3), 616–623. https://doi.org/10.1002/ecy.1689

Shen, W., Lin, Y., Jenerette, G. D., & Wu, J. (2011). Blowing litter across a landscape: Effects on ecosystem nutrient flux and implications for landscape management. Landscape Ecology, 26(5), 629–644. https://doi.org/10.1007/s10980-011-9599-x

Shepard, E. L. C., Wilson, R. P., Rees, W. G., Grundy, E., Lambertucci, S. A., & Vosper, S. B. (2013). Energy landscapes shape animal movement ecology. The American Naturalist, 182(3), 298–312. https://doi.org/10.1086/671257

Shure, D. J., & Wilson, L. A. (1993). Patch-size effects on plant phenolics in successional openings of the southern appalachians. Ecology, 74(1), 55–67. https://doi.org/10.2307/1939501

Smith, C. K., Coyea, M. R., & Munson, A. D. (2000). Soil carbon, nitrogen, and phosphorus stocks and dynamics under disturbed black spruce forests. Ecological Applications, 10(3), 775–788. https://doi.org/10.1890/1051-0761(2000)010[0775:SCNAPS]2.0.CO;2

Smithwick, E. A. H., Harmon, M. E., & Domingo, J. B. (2003). Modeling multiscale effects of light limitations and edge-induced mortality on carbon stores in forest landscapes. Landscape Ecology, 18(7), 701–721. https://doi.org/10.1023/B:LAND.0000004254.94982.67

South, R. G. (1983). Biogeography and ecology of the island of Newfoundland (Vol. 48).

Strahan, R. T., Meador, A. J. S., Huffman, D. W., & Laughlin, D. C. (2016). Shifts in community-level traits and functional diversity in a mixed conifer forest: A legacy of land-use change. Journal of Applied Ecology, 53(6), 1755–1765. https://doi.org/10.1111/1365-2664.12737

Tang, Z., Xu, W., Zhou, G., Bai, Y., Li, J., Tang, X., Chen, D., Liu, Q., Ma, W., Xiong, G., He, H., He, N., Guo, Y., Guo, Q., Zhu, J., Han, W., Hu, H., Fang, J., & Xie, Z. (2018). Patterns of plant carbon, nitrogen, and phosphorus concentration in relation to productivity in China’s terrestrial ecosystems. Proceedings of the National Academy of Sciences, 115(16), 4033–4038. https://doi.org/10.1073/pnas.1700295114

Turner, Monica G. (1989). Landscape ecology: The effect of pattern on process. Landscape Ecology, 20, 171–197.

Turner, Monica Goigel. (2005). Landscape ecology: What is the state of the science? Annual Review of Ecology, Evolution, and Systematics, 36(1), 319–344. https://doi.org/10.1146/annurev.ecolsys.36.102003.152614

Urbina, I., Sardans, J., Grau, O., Beierkuhnlein, C., Jentsch, A., Kreyling, J., & Peñuelas, J. (2017). Plant community composition affects the species biogeochemical niche. Ecosphere, 8(5), e01801. https://doi.org/10.1002/ecs2.1801

U.S. Geological Survey. (2017). Landsat Quality Assurance ArcGIS Toolbox. doi: 10.5066/F7JM284N

Vermote, E., Justice, C., Claverie, M., & Franch, B. (2016). Preliminary analysis of the performance of the Landsat 8/OLI surface reflectance product. Remote Sensing of Environment, 185, 46–56. https://doi.org/10.1016/j.rse.2016.04.008

Vranken, I., Baudry, J., Aubinet, M., Visser, M., & Bogaert, J. (2015). A review on the use of entropy in landscape ecology: Heterogeneity, unpredictability, scale dependence and their links with thermodynamics. Landscape Ecology, 30(1), 51–65. https://doi.org/10.1007/s10980-014-0105-0

Wam, H. K., Felton, A. M., Stolter, C., Nybakken, L., & Hjeljord, O. (2018). Moose selecting for specific nutritional composition of birch places limits on food acceptability. Ecology and Evolution, 8(2), 1117–1130. https://doi.org/10.1002/ece3.3715

Waring, R. H., Coops, N. C., Fan, W., & Nightingale, J. M. (2006). MODIS enhanced vegetation index predicts tree species richness across forested ecoregions in the contiguous U.S.A. Remote Sensing of Environment, 103(2), 218–226. https://doi.org/10.1016/j.rse.2006.05.007

Zhang, L.-L., & He, W.-M. (2009). Consequences of ramets helping ramets: No damage and increased nutrient use efficiency in nurse ramets of Glechoma longituba. Flora - Morphology, Distribution, Functional Ecology of Plants, 204(3), 182–188. https://doi.org/10.1016/j.flora.2008.02.001

Zhao, N., He, N., Wang, Q., Zhang, X., Wang, R., Xu, Z., & Yu, G. (2014). The altitudinal patterns of leaf c∶n∶p stoichiometry are regulated by plant growth form, climate and soil on changbai mountain, china. PLOS ONE, 9(4), e95196. https://doi.org/10.1371/journal.pone.0095196

